# A multivalent peptide-polymer conjugate material mimics STING to therapeutically activate innate immune signaling

**DOI:** 10.64898/2026.03.24.712780

**Authors:** Justin A. Kaskow, Julia Treese, Anita Gaenko, Victoria F. Gomerdinger, Zhi Kai Tio, Margaret M. Billingsley, Aidan Kindopp, Paula T. Hammond

## Abstract

Stimulator of interferon genes (STING) is a promising therapeutic target for cancer immunotherapy, but agonists are often rendered ineffective by the loss of STING expression in cancer cells. Here we engineer a multivalent peptide-polymer conjugate material that can easily be delivered to the cytosol, where it mimics key protein interactions from the missing STING protein to directly activate downstream innate immune signaling. While previously developed STING mimicking therapeutics use nearly the full STING protein, this material contains only a 39 amino acid peptide from the STING C-terminal tail that includes interaction motifs for downstream kinase TBK1 and transcription factor IRF3. Conjugation of multiple peptide copies to a negatively charged polymer backbone mimics the multivalent protein-protein interactions of the oligomerized STING signaling complex, activating TBK1 and IRF3 as well as the transcription of downstream genes in both STING-proficient and STING-silenced cancer cell lines. We optimize a lipid nanoparticle formulation to deliver this conjugate material intracellularly, allowing for its application as an immunotherapy for ovarian cancer. Treatment with the STING mimicking conjugate material promoted the production of type I interferons, repolarization of myeloid cells to an anti-tumor phenotype, and recruitment of T cells to tumors in mice. This treatment ultimately led to tumor regression and extended survival in multiple mouse models of metastatic ovarian cancer. Overall, this work highlights the potential of peptide-polymer conjugate mimics of STING to therapeutically activate innate immune signaling.

## INTRODUCTION

Drugging the stimulator of interferon genes (STING) signaling pathway shows great promise in treating cancer. Activated by the leakage of DNA into the cytosol which has been observed in cancer cells,^1^ this innate immune pathway is central to the spontaneous immune mediated elimination of early tumors by initiating a type I interferon (IFN) response.^2,3^ Pharmacological targeting of this pathway has primarily focused on developing small molecule agonists of the STING protein. STING agonists have demonstrated strong efficacy in preclinical models of cancer, leading to cancer-specific immune responses and the complete elimination of tumors.^4–9^ However, translation of STING agonists to the clinic has thus far not been successful.^10–13^

One major biological limitation of STING agonism as a therapeutic strategy is that STING expression is commonly lost in human cancer,^14–19^ presumably as a mechanism to escape immune surveillance. While host immune cells retain functional STING and targeting immune cells has demonstrated therapeutic benefit,^5,20,21^ there is evidence that STING activation directly in cancer cells is required for optimal anti-cancer immune responses.^22–24^ STING activation in cancer cells is correlated with increased cancer cell immunogenicity ^22,24–28^ and the secretion of additional anti-tumor cytokines.^23,24,29^ Further, experiments using STING-deficient mice implanted with STING-proficient tumors demonstrate that activation of STING signaling in cancer cells alone can generate potent adaptive immune responses and inhibit tumor progression, even without STING activity in host immune cells.^22^ Thus, there is a strong rationale to develop new therapeutic strategies capable of activating STING signaling in cancer cells where traditional STING agonists fail. Direct activation of the downstream transcription factor interferon regulatory factor 3 (IRF3) would overcome this challenge by bypassing the need for STING expression in cancer; however, IRF3 is an intracellular target that is typically activated only through multiple interactions with the molecules present in the STING signaling complex. Inspired by the polymeric signaling complex formed during natural STING activation,^30^ we develop a multivalent peptide-polymer conjugate material that can be delivered to the cytosol and synthetically mimic key protein-protein interactions of the STING signaling complex. This multivalent peptide conjugate is capable of scaffolding the signaling molecules required to activate IRF3, enabling the desirable downstream outcomes of STING activation even in STING-deficient cancer.

When functioning properly, STING signaling (**Figure 1A**) begins when cytosolic DNA activates cGAMP synthase (cGAS), which catalyzes the production of the cyclic dinucleotide 2’,3’-cGAMP.^31–33^ 2’,3’-cGAMP binds to a pocket in the cytosolic domain of the endoplasmic reticulum resident STING protein, inducing conformational changes that initiate STING trafficking,^34^ multimerization,^30,35,36^ and free STING’s flexible C-terminal tail.^30^ Activated STING multimers act as a scaffold presenting many copies of the C-terminal tail (**Figure 1B**), which contains a PLPLRT/SD motif that recruits the kinase TANK-binding kinase 1 (TBK1).^35^ The generation of clusters of multiple TBK1 molecules on this scaffold allows for their coordinated trans-autophosphorylation and activation. TBK1 then phosphorylates a pLxIS motif on the C-terminal tail of a neighboring STING molecule.^37^ The phosphorylated pLxIS motif recruits the transcription factor IRF3, which itself is phosphorylated by TBK1 and converted into its dimeric active state.^38^ Active IRF3 enters the nucleus and drives the transcription of Type I IFNs and interferon stimulated genes, that ultimately drive the downstream innate immune response. A wide variety of strategies to disrupt STING signaling have been observed in cancer, ^39–41^ but one of the most common is the epigenetic silencing of upstream proteins cGAS and STING (**Figure 1A**).^14–19^ Loss of STING expression in particular is observed in 50% of patients with metastatic melanoma ^15^ and over 70% of patients with metastatic ovarian cancer.^16^

**Figure 1:**
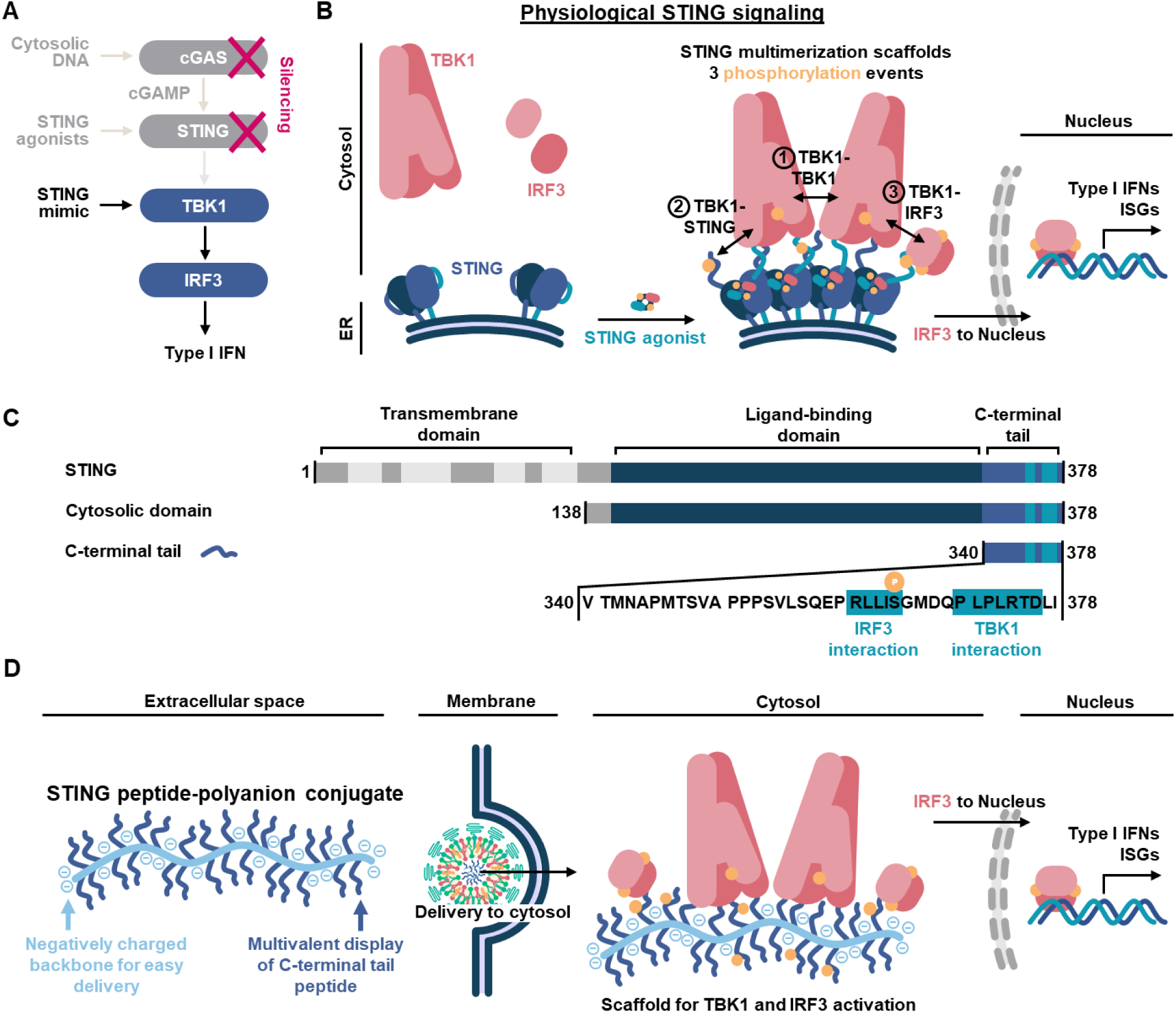
Designing a material to mimic protein-protein interactions of the STING signaling complex. (**A**) A schematic of the STING signaling cascade, noting that upstream molecules cGAS and STING are frequently epigenetically silenced in cancer, leaving cytosolic DNA or STING agonists incapable of activating downstream signaling. A STING mimicking therapeutic instead acts directly on downstream TBK1 and IRF3 to enable activity in STING silenced cells. (**B**) Binding a STING agonist frees the STING C-terminal tail and results in STING multimerization. Multimerized STING acts as a scaffold to promote protein-protein interactions that enable the autophosphorylation of TBK1, allows TBK1 to phosphorylate the STING C-terminal tail to enable IRF3 recruitment, and then allows TBK1 to phosphorylate IRF3. Active phosphorylated IRF3 transits to the nucleus where it results in the transcription of IRF3 controlled genes such as Type I IFNs and interferon stimulated genes (ISGs). (**C**) Schematic of mouse STING protein, comparing full STING to the cytosolic domain that is used in existing STING mimicking therapeutics and the STING C-terminal tail that was used in this work. The TBK1 and IRF3 interaction motifs in the C-terminal tail are highlighted, noting that phosphorylation of S365 is required for IRF3 interaction. (**D**) The STING peptide-polyanion conjugate was designed with a polyanion backbone to promote electrostatic interactions with positively charged delivery vehicles and allow easy delivery to the cytosol. Multiple copies of the STING C-terminal tail peptide are conjugated to this backbone to mimic the high-valency display of the STING C-terminal tail found in the multimerized active state of STING. In the cytosol, the STING peptide-polyanion conjugate mimics the protein-protein interaction motifs of multimerized STING to enable TBK1 and IRF3 activation, and the resulting transcription of IRF3 controlled genes.

Two main approaches have been developed to overcome STING dysregulation in cancer: epigenetic reprogramming and STING mimicking therapeutics. Existing epigenetic reprogramming strategies utilize inhibitors of proteins involved in the epigenetic silencing of STING, and have been shown to reactivate STING expression and improve the efficacy of STING agonists.^18,22,27,42^ However, current epigenetic reprogramming strategies risk off-target effects on cancer and host cell phenotypes and still require subsequent treatment with a STING agonist, complicating treatment regimens. Instead, a more direct strategy is to bypass the need for endogenous STING in cancer cells entirely, by “replacing” its function with mimics to directly activate TBK1 and IRF3. Existing STING mimicking therapeutic strategies deliver an activated version of STING to the cytosol where it can interact with endogenous TBK1 and IRF3, either by delivering STING protein directly ^43,44^ or a nucleic acid encoding STING. ^45–47^ To induce the multimerization required for STING activation, these therapies deliver STING that has been premixed with its ligand,^43^ aggregated by interactions with a delivery vehicle,^44^ mutated to drive constitutive multimerization,^45,46^ or fused to a self-oligomerizing protein domain.^47^ Though these therapeutics have demonstrated efficacy in murine models of cancer, ^43,47,48^ they face some limitations. The strategies to induce multimerization are currently limited to lower valences of 4 to 8 ^43,47^ or left to uncontrolled processes inside the cell.^44–46^ For protein STING mimics, delivery of functional protein to the cytosol can be challenging and would be expected to require time-consuming development of a bespoke nanoparticle carrier. Further, existing designs use the full STING protein including the folded ligand-binding domain (**Figure 1C**). Mechanistically, the ligand-binding domain controls STING multimerization, making it redundant when used alongside these alternative means of inducing multimerization. This redundancy is supported by studies that show STING protein fragments containing the C-terminal tail maintain their ability to activate TBK1 and IRF3 even after the ligand-binding domain is thermally denatured ^44^ or removed from the protein entirely.^38^ Ultimately, we believe that the design of STING mimics could be simplified while developing more effective therapeutics and improving deliverability.

We hypothesized that a molecule presenting multiple copies of only the STING C-terminal tail would contain the protein interaction motifs necessary to engage and strongly activate downstream proteins TBK1 and IRF3 (**Figure 1D**), so we built a therapeutic on a polymer scaffold to facilitate high-valency peptide display. Using a negatively charged, polyanion backbone promoted electrostatic interactions to enable straightforward encapsulation and delivery using existing nanocarriers designed to deliver negatively charged nucleic acids. When delivered to the cytosol, this multivalent peptide-polyanion conjugate was observed to phosphorylate and activate TBK1 and IRF3, resulting in the TBK1-dependent expression of IRF3 controlled genes. Peptide-polyanion conjugate treatment initiated signaling comparable to a high dose of the STING agonist ADU-S100 in a cancer cell line with functional STING signaling and also retained activity even in a STING-silenced cancer cell line that showed no response to ADU-S100. To enable application as a therapeutic, we developed an ionizable lipid nanoparticle (LNP) formulation capable of effectively delivering this conjugate material into the cytosol. As a proof of concept, we used conjugate-loaded LNP to treat ovarian cancer, a disease that has responded poorly to traditional immunotherapies and has an extremely high frequency of STING inactivation that may limit existing STING agonists.^16^ In mice, administration of conjugate-loaded LNP increased IRF3 controlled cytokine levels and repolarized the tumor microenvironment towards an inflamed state. Ultimately, STING peptide-polyanion conjugate therapy was able to shrink tumors and prolong survival in multiple mouse models of metastatic ovarian cancer, demonstrating the therapeutic potential of this strategy.

## RESULTS

### Designing a Multivalent Peptide-Polyanion Conjugate Material

Despite prior biochemical work demonstrating that the STING C-terminal tail alone is capable of activating TBK1 and IRF3, it is still challenging to apply this strategy therapeutically. The initial biochemical demonstration of activity occurred in an *in vitro* reconstitution system, where high molecular weight aggregates of the STING C-terminal tail were directly added to a solution containing TBK1 and IRF3.^38^ A therapeutic would additionally need to cross the cell membrane. In our prior work, we attempted to deliver the STING C-terminal tail into the cytosol of live cells using a transfection reagent;^44^ however, we observed no detectible activation of IRF3. This result implied that the STING C-terminal tail failed to be delivered into the cytosol effectively, failed to multimerize into an active form, or both. Here, we overcome both these challenges by designing a multivalent STING C-terminal tail peptide-polyanion conjugate for straightforward delivery to the cytosol and high intrinsic valency.

We built the conjugate material on a poly(L-glutamate) backbone, as this polyanion has a high negative charge density, is biocompatible, and a controlled number of carboxylate side chains can be modified to generate reactive alkyne handles. A 300-repeat unit polymer with alkyne groups grafted onto 10% of side chains was selected to enable high valency display of 30 peptides on each molecule while retaining a strong negative charge. The murine STING C-terminal tail, mSTING(340-378), is only 39 amino acids, allowing for production via solid-phase peptide synthesis (**Figure 2A**). An azido-lysine was added as a reactive handle at the N-terminus of the peptide to prevent steric interference with the known protein interaction motifs at the C-terminus. Peptide-polyanion conjugates were synthesized by copper-catalyzed azide-alkyne cycloaddition with excess azide-peptide (**Figure 2B**). Complete conversion of alkyne groups on the polyanion to peptide was confirmed by measuring the disappearance of detectable alkyne groups in the product using a fluorescence-based alkyne detection assay and via NMR (**Figure S1-S5**). The removal of excess peptide after purification was confirmed using size exclusion chromatography, which also demonstrated the expected increase in size of the peptide-polyanion product compared to peptide and polyanion starting materials (**Figure 2C**). The mSTING(340-378) peptide showed a random coil secondary structure as measured by circular dichroism, both alone and after conjugation to poly(L-glutamate) (**Figure 2D**), as expected given the known flexibility of the STING C-terminal tail.^37,49,50^

**Figure 2:**
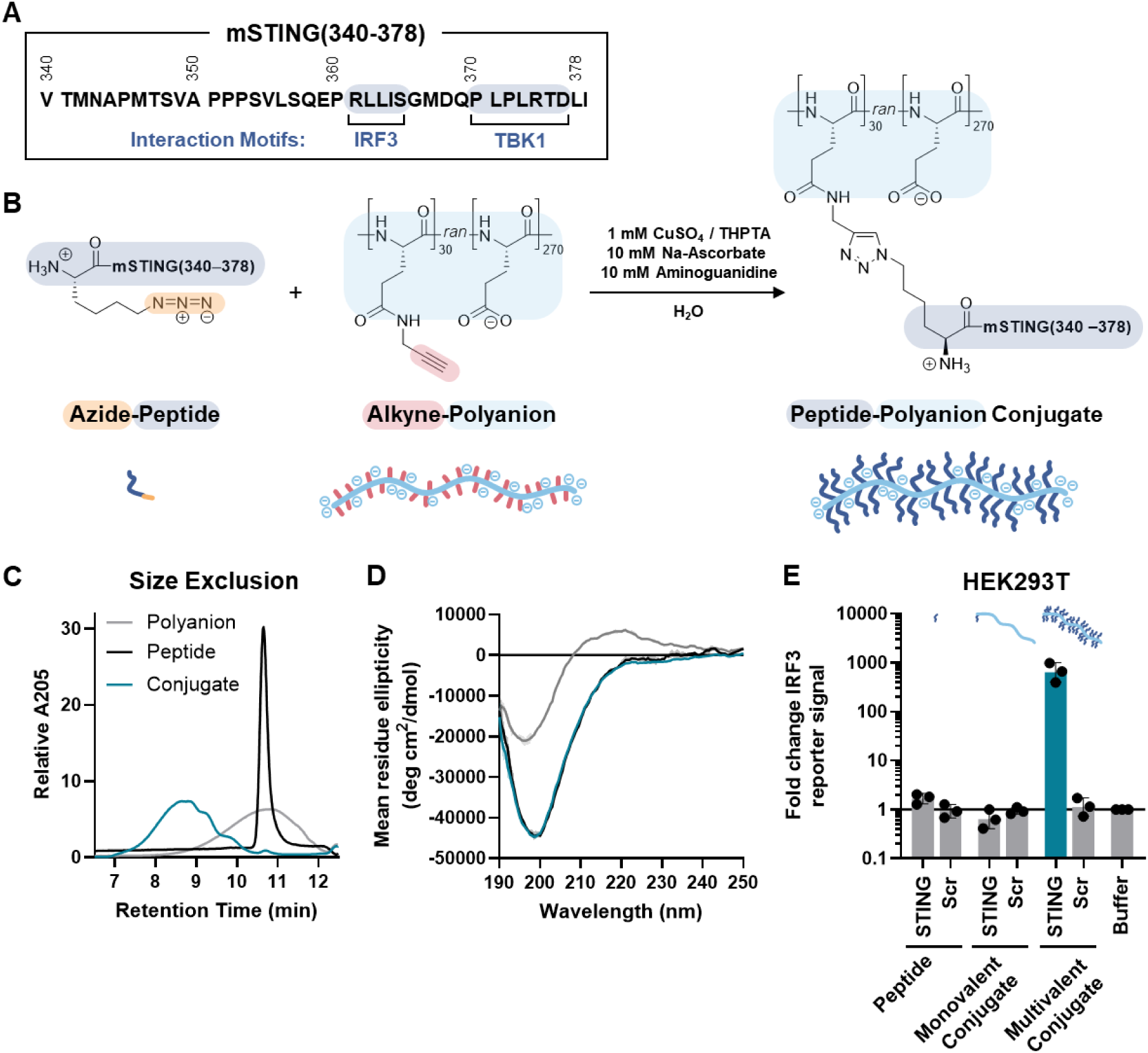
Synthesis and characterization of a STING mimicking peptide-polyanion conjugate. (**A**) Sequence of the murine STING C-terminal tail peptide mSTING(340-378), highlighting motifs known to interact with TBK1 and IRF3. (**B**) Peptide-polyanion conjugate was synthesized through copper-catalyzed azide-alkyne cycloaddition of an azidolysine-mSTING(340-378) peptide and the polyanion poly(L-glutamate)-graft-alkyne. (**C**) HPLC chromatogram using size exclusion column of azidolysine-mSTING(340-378), poly(L-glutamate)-graft-alkyne, and poly(L-glutamate)-graft-mSTING(340-378). (**D**) Circular dichroism spectrum of azidolysine-mSTING(340-378), poly(L-glutamate)-graft-alkyne, and poly(L-glutamate)-graft-mSTING(340-378) in 1× PBS. (**E**) Activity of the peptide-polyanion conjugate was examined using a HEK293T-derived reporter with luciferase expression under the control of target transcription factor IRF3. IRF3 reporter signal relative to buffer treatment for HEK293T reporter cells treated with 2 μM STING or Scr peptide alone, conjugated to only C-terminus of size-matched poly(L-glutamate) to generate a monovalent peptide-polyanion conjugate control, or conjugated to side chains as specified in (**B**) to generate a multivalent peptide-polyanion conjugate. All treatments were transfected using TransIT-X2, activity was measured 24 h post treatment (N = 3 biological replicates). Data represented as geometric mean ± SD.

To confirm that the STING C-terminal tail peptide-polyanion conjugate is capable of activating STING signaling, the commercial transfection reagent TransIT-X2 was used to deliver it to the cytosol of a HEK293T-derived reporter cell line. This reporter expresses luciferase under the control of the target transcription factor IRF3, allowing IRF3 activation to be quantified. Treatment with a multivalent peptide-polyanion conjugate containing 30-copies of the mSTING(340-378) peptide resulted in strong IRF3 activation, nearly 1000-fold over baseline levels (**Figure 2E**). As a control, we synthesized conjugates with an identical design but using a scrambled (Scr) peptide. No increase in IRF3 activation was observed after treatment with a Scr control conjugate, indicating that this activity depends on the mSTING(340-378) peptide sequence and is not a result of non-specific effects of the transfection reagent or the introduction of exogenous material into the cytosol. A monovalent conjugate with only a single copy of the mSTING(340-378) peptide conjugated to the C-terminus of a size-matched 300 repeat unit poly(L-glutamate) was synthesized to control for the valency of peptide display (**Figure S6-S8**). At an equal molar dose of peptide, the monovalent conjugate showed no increase in IRF3 activity, confirming that multivalent display is required for activity (**Figure 2E**). Multiple independently synthesized batches of multivalent peptide-polyanion conjugate showed consistent IRF3 activity in HEK293T (**Figure S9**). This conjugate material appears resistant to thermal denaturation, as it maintains consistent IRF3 activity in HEK293T after exposure to a 95 *°*C thermal stress of up to 10 min (**Figure S10**). Overall, these results confirm that multivalent display of the STING C-terminal tail sequence on a polyanion can activate the target transcription factor IRF3, supporting further examination of this multivalent conjugate.

### Conjugate Mimics STING to Activate Downstream Innate Immune Signaling

To confirm that the STING peptide-polyanion conjugate is capable of activating TBK1 and IRF3, we treated HEK293T cells using TransIT-X2 for intracellular delivery and examined the resulting cellular response. Confocal microscopy showed puncta of TBK1 co-localized with puncta of the conjugate, providing evidence that the conjugate is capable of recruiting TBK1 in the cytosol (**Figure 3A, Figure S11**). Phosphorylation of TBK1 and IRF3 was observed by Western blot, confirming on-target activation of these molecules (**Figure 3B, Figure S12**). Pretreatment of cells with the specific TBK1 inhibitor MRT67307 completely eliminated IRF3 activation after STING peptide-polyanion conjugate treatment (**Figure 3C**), confirming that TBK1 is required for activity of the conjugate. Overall, these results provide evidence that cytosolically delivered peptide-polyanion conjugate is capable of mimicking STING to activate TBK1 and IRF3.

**Figure 3:**
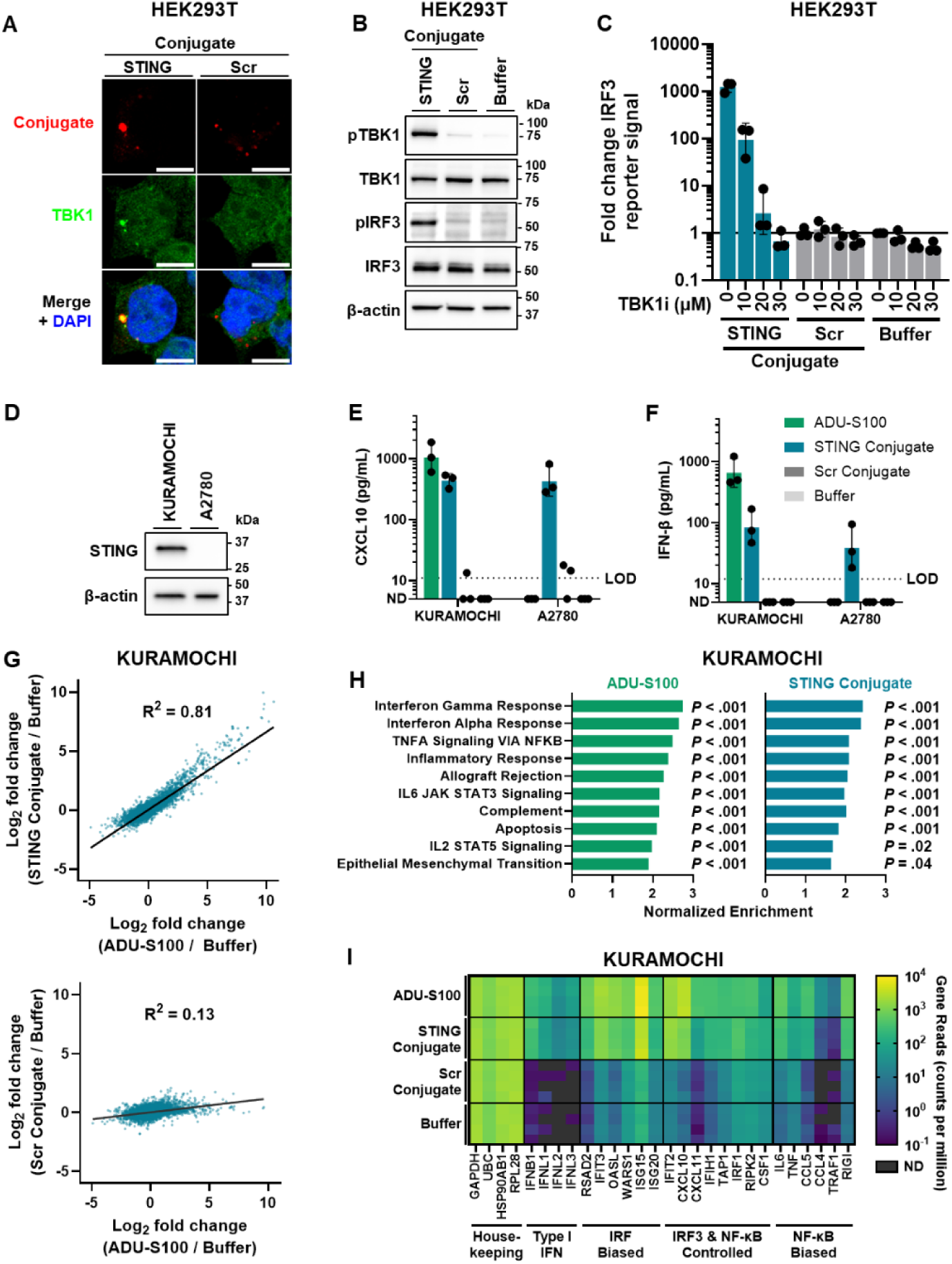
STING peptide-polyanion conjugate activates innate immune signaling even in STING-deficient cancer cells. (**A**) HEK293T reporter cells were treated with 6.0 μg/mL STING or Scr conjugate using TransIT-X2 as a vehicle, cells were imaged by confocal microscopy at 6 h post treatment to examine conjugate colocalization with target TBK1 (representative of N = 3 biological replicates). Scale bar is 10 μm. (**B**) HEK293T reporter cells were treated with 8.3 μg/mL STING or Scr conjugate using TransIT-X2 as a vehicle, Western blot was performed at 6 h post treatment to examine TBK1 and IRF3 phosphorylation (representative of N = 3 biological replicates). (**C**) IRF3 reporter signal relative to buffer treatment for HEK293T reporter cells pretreated for 6 h with TBK1 inhibitor MRT67307 (TBK1i) and then treated with 8.3 μg/mL STING or Scr conjugate delivered using TransIT-X2, measured 24 h post treatment (N = 3 biological replicates). (**D**) Western blot of STING and β-actin expression in ovarian cancer cell lines KURAMOCHI and A2780. (**E-F**) KURAMOCHI and A2780 ovarian cancer cell lines were treated with 5.0 μg/mL STING or Scr conjugate using TransIT-X2 as a vehicle or 50 μM STING agonist ADU-S100. (**E**) CXCL10 and (**F**) IFN-β in supernatant was measured by ELISA 24 h post treatment (N = 3 biological replicates). Replicates where analyte was below the limit of detection (LOD) are labeled as not detected (ND), no summary statistics were computed if any replicate was ND. (**G-I**) KURAMOCHI cells were treated with 5.0 μg/mL STING or Scr conjugate using TransIT-X2 as a vehicle or 50 μM ADU-S100, mRNA sequencing was performed at 6 h post treatment (N = 4 biological replicates). (**G**) Plot of Log2 fold change of STING conjugate or ADU-S100 treatment compared to Buffer, showing high correlation between treatments. Plot of Log2 fold change of Scr conjugate or ADU-S100 treatment compared to Buffer is displayed below as a control, showing greatly reduced correlation. The coefficient of determination R^2^ for line of best fit is displayed. (**H**) Gene set enrichment analysis was performed on MSigDB Hallmark gene set, normalized enrichment and adjusted *P* value are displayed for the 10 gene sets significantly enriched (*P* < .05) when comparing STING to Scr conjugate. Normalized enrichment and adjusted *P* values for the same 10 gene sets are displayed for the comparison of ADU-S100 to Buffer (**I**) Heatmap of gene expression for selected genes. Replicates where a given gene was not detected are labeled ND. Data represented as geometric mean ± SD.

We then examined if the STING peptide-polyanion conjugate could activate IRF3 in cancer cells with silenced STING. We compared treatment with the clinically advanced STING agonist ADU-S100 ^10,11^ to treatment with the STING peptide-polyanion conjugate in two ovarian cancer cell lines: KURAMOCHI cells that express functional STING and A2780 cells that have no detectable STING expression (**Figure 3D, Figure S13**). As expected, the STING agonist ADU-S100 only induced secretion of the IRF3 controlled cytokines CXCL10 and IFN-β in STING-proficient KURAMOCHI cells, while no detectable cytokines were observed after treatment of STING-deficient A2780 cells (**Figure 3E-F**). Treatment with the STING peptide-polyanion conjugate resulted in secretion of CXCL10 and IFN-β in both KURAMOCHI and A2780 (**Figure 3E-F**). This demonstrates that the STING peptide-polyanion conjugate is capable of activating desired innate immune signaling even in STING-deficient cancer cells that do not respond to STING agonists.

The cellular response to treatment with the STING peptide-polyanion conjugate was examined in greater detail using mRNA sequencing. Treatment with the STING peptide-polyanion conjugate was compared to treatment with the STING agonist ADU-S100 in the STING-proficient cell line KURAMOCHI. At 6 h after treatment, over 500 differentially expressed genes were observed for both STING peptide-polyanion conjugate and ADU-S100 treatment compared to a buffer control (**Figure S14**). Transcriptional changes were highly similar between the STING agonist and STING mimic treatment. Compared to a buffer treated control, differential gene expression following STING peptide-polyanion conjugate treatment was well correlated with gene expression following ADU-S100 treatment (line of best fit R^2^ = 0.81), with no obvious outliers that showed large changes in expression with one treatment but not the other (**Figure 3G**). To contrast this, differential gene expression following Scr peptide-polyanion conjugate treatment was only weakly correlated with gene expression following ADU-S100 treatment (line of best fit R^2^ = 0.13), with genes that were strongly upregulated by ADU-S100 seeing no notable change after treatment with the control conjugate (**Figure 3G**). Gene set enrichment analysis (GSEA) using the MSigDB hallmark gene sets ^51^ showed 10 gene sets significantly enriched (adjusted *P* ≤ .05) when comparing STING to Scr peptide-polyanion conjugate treatment. These 10 enriched gene sets were the same as the top 10 gene sets enriched (*P* ≤ .05) by ADU-S100 treatment (**Figure 3H**). Combined, these results provide evidence that STING peptide-polyanion conjugate treatment activates on-target STING signaling without any major induction of off-target signaling.

MSigDB hallmark gene sets strongly enriched after STING peptide-polyanion conjugate treatment include “IFNα response” and “IFNγ response”, consistent with activation of the transcription factor IRF3 (**Figure 3H**). Other enriched gene sets include “TNFα signaling via NF-κB” and “IL6 JAK STAT3 signaling”, consistent with activation of the transcription factor NF-κB. NF-κB activation is known to follow STING activation, downstream of TBK1 but independent of canonical IRF3 activation.^52^ Strong increases in specific IRF3 controlled genes such as *RSAD2*, *ISG15*, and *ISG20* and NF-κB controlled genes such as *IL6*, *CCL5*, and *RIGI* are observed following both STING peptide-polyanion conjugate and ADU-S100 treatment (**Figure 3I**). Activation of both IRF3 and NF-κB transcription factors was confirmed in a THP1 monocyte dual transcription factor reporter cell line, where treatment with the STING peptide-polyanion conjugate resulted in dose-dependent activation of IRF3 and NF-κB, with no notable bias towards stronger IRF3 or NF-κB activation compared to ADU-S100 (**Figure S15**). These results demonstrate that the STING peptide-polyanion conjugate effectively engages therapeutically relevant innate immune signaling downstream of STING, including both IRF3 and NF-κB transcription factors.

### Designing Lipid Nanoparticles for Effective Conjugate Delivery

The therapeutic effect of the STING peptide-polyanion conjugate requires successful delivery to the cytosol for the conjugate to engage its targets TBK1 and IRF3. Development of an effective delivery vehicle is crucial for the STING peptide-polyanion conjugate to be utilized as a therapeutic *in vivo*. A wide variety of delivery vehicles have been developed to encapsulate and deliver nucleic acids intracellularly, often by leveraging electrostatic interactions between the negatively charged nucleic acid cargo and a positively charged nanocarrier. We hypothesized that designing the STING peptide-polyanion conjugate with a strongly negatively charged backbone would mimic the physicochemical properties of nucleic acids, thereby allowing for encapsulation and delivery using existing positively charged nanocarriers designed for nucleic acid delivery. To test this hypothesis, we attempted to deliver the STING peptide-polyanion conjugate using a panel of nucleic acid carriers and measured IRF3 activation in a HEK293T-based reporter cell line as a functional metric of successful intracellular delivery. We tested lipid nanoparticle (LNP) formulations using ionizable lipids ALC-0315, MC3, and SM-102 from three clinically approved LNP therapeutics, using 50:10:38.5:1.5 (mol ratio) ionizable lipid:DOPE:cholesterol:DMG-PEG2000 to reflect the composition featured in clinically-approved LNP formulations.^53–55^ We also examined cationic polymer-based delivery vehicles including linear poly(ethyleneimine) (PEI), the poly(β-amino ester) (PBAE) “Poly2”,^56,57^ and the commercial transfection reagent TransIT-X2. All vehicles achieved functional delivery of the conjugate, demonstrated by a significant increase in IRF3 activation compared to a Scr control delivered using the same vehicle (**Figure 4A**). All vehicles showed a >200-fold increase in IRF3 activation relative to buffer treated controls, with the exception of PEI which showed a weaker 40-fold increase in IRF3 activation. The SM-102 LNP formulations showed an approximately 10-fold increase in IRF3 activation, even when loaded with the inactive Scr peptide-polyanion conjugate. This is expected as LNPs themselves are known to activate innate immune signaling through interaction of the ionizable lipid with pattern recognition receptors and disruption caused during endosomal escape, with stronger innate immune activation being observed with SM-102 in prior literature.^58–61^ Overall, these results demonstrate that the STING peptide-polyanion conjugate can be effectively delivered to the cytosol using a variety of “off-the-shelf” nucleic acid delivery vehicles.

**Figure 4:**
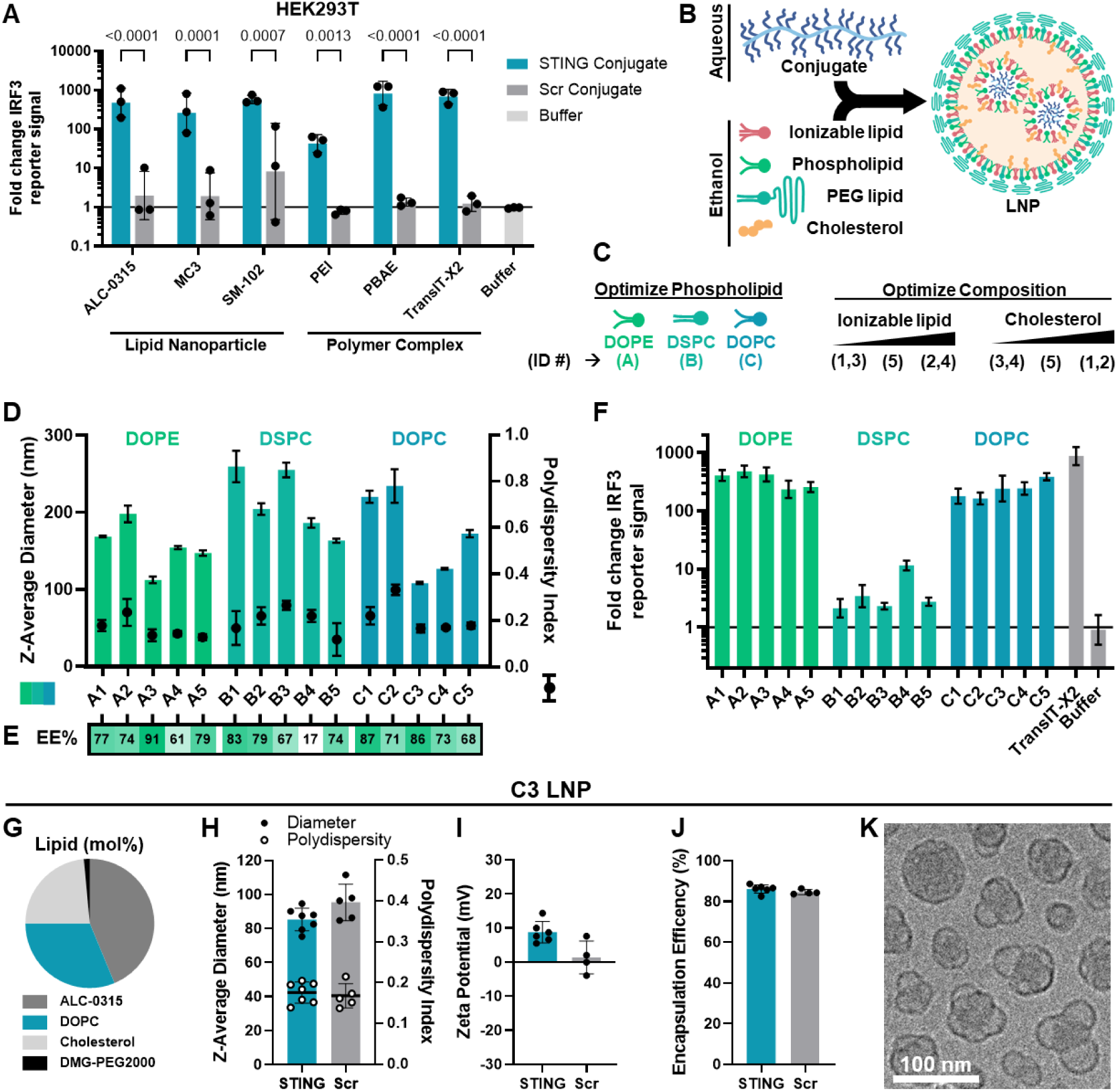
Designing a LNP carrier for peptide-polyanion conjugate delivery. (**A**) Ease of peptide-polyanion conjugate delivery was tested using a variety of established nucleic acid delivery methods: LNPs prepared with the ionizable lipids ALC-0315, MC3, and SM-102 or polymer complexes prepared using PEI, the PBAE “Poly2”, or the transfection reagent TransIT-X2. IRF3 reporter signal relative to buffer treatment for HEK293T reporter cells treated with 4.2 μg/mL STING or Scr conjugate delivered using specified delivery method, 24 h post treatment (N = 3 biological replicates). *P* values computed with a two-way ANOVA on log-transformed IRF3 reporter signal followed by Šídák’s post-hoc test are displayed above each figure, comparing STING to Scr conjugate for each delivery method. (**B**) Schematic of LNP formation method, where conjugate in aqueous phase is mixed with lipids in ethanol phase to generate nanoparticles. (**C**) An ALC-0315-based LNP formulation was optimized by screening a library of LNPs with varied phospholipid components (DOPE, DSPC, DOPC) and molar ratios of ALC-0315, phospholipid, and cholesterol. (**D**) LNP Z-Average diameter and polydispersity index measured by DLS (N = 3 technical replicates). (**E**) Encapsulation efficiency (EE%) measured by native PAGE assay. (**F**) IRF3 reporter signal relative to buffer treatment for HEK293T reporter cells treated with 3.3 μg/mL conjugate delivered by each LNP formulation or TransIT-X2 control, 24 h post treatment (N = 3 technical replicates). (**G**) C3 LNP formulation selected for further examination is composed of 43.8 : 31.3 : 23.5 : 1.5 (mol ratio) ALC-0315 : DOPC : Cholesterol : DMG-PEG2000. The C3 formulation was loaded with STING or Scr conjugate and tested for (**H**) LNP Z-Average diameter and polydispersity index measured by DLS (N = 7 (STING) or N = 5 (Scr) LNP batches), (**I**) zeta potential (N = 6 (STING) or N = 4 (Scr) LNP batches), and (**J**) Encapsulation efficiency by native PAGE assay (N = 6 (STING) or N = 4 (Scr) LNP batches). (**K**) Cryo-TEM image of C3 LNP formulation containing STING conjugate (representative of N = 2 LNP batches). Data represented as mean ± SD for data on linear scale and geometric mean ± SD for data on log-scale.

We decided to advance a LNP formulation for peptide-polyanion conjugate delivery (**Figure 4B**) due to this vehicle’s widespread use in the clinic. ALC-0315 was selected for its strong functional delivery ability and low off-target IRF3 activation of the LNP itself compared to SM-102 (**Figure 4A**). The initial 50:10:38.5:1.5 (mol ratio) ALC-0315:DOPE:cholesterol:DMG-PEG2000 formulation (labeled formulation A2) showed strong functional delivery in HEK293T *in vitro*. However, at 200 nm LNP size was larger than typical nucleic acid formulations, size increased during the centrifugal filtration process used to purify and concentrate LNPs, and the encapsulation efficiency (EE) was lower than ideal at 74% (**Figure S16**). Thus, we optimized the LNP formulation, aiming to decrease size and increase encapsulation efficiency. To this end, we screened a library of LNPs with varying phospholipid components and compositions (**Figure 4C**). Ultimately, LNPs with high phospholipid and reduced cholesterol and ALC-0315 content, containing either DOPE (A3) or DOPC (C3), led to smaller diameters at approximately 100 nm, polydispersity indices < 0.2 (**Figure 4D**), and EEs >85% (**Figure 4E, Figure S17**). All DOPE and DOPC based formulations performed similarly in the HEK293T functional delivery assay, leading to a >100-fold increase in IRF3 activation (**Figure 4F**). The inclusion of DSPC on the other hand rendered LNPs ineffective at delivering the conjugate to the cytosol, leading to only 2- to 10- fold increases in IRF3 activation (**Figure 4F**). We moved forward with the C3 formulation over the A3 formulation because the A3 formulation showed signs of instability during purification by centrifugal filtration, increasing in diameter by 20 nm (**Figure S16**).

The final C3 LNP formulation was prepared at 43.8:31.3:23.5:1.5 (mol ratio) ALC-0315:DOPC:cholesterol:DMG-PEG2000 (**Figure 4G, Figure S18**). Validation experiments generating LNPs loaded with both STING and Scr peptide-polyanion conjugate showed that this formulation consistently produces 80-100 nm diameter LNPs with a polydispersity index < 0.2 (**Figure 4H**), a near neutral zeta potential (**Figure 4I**), and an EE of 85% (**Figure 4J, Figure S19**). LNPs showed an apparent pK_a_ of 6.4 that is consistent with other ALC-0315 LNP formulations loaded with mRNA (**Figure S20**).^62^ CryoTEM showed that STING peptide-polyanion conjugate loaded C3 LNPs have an electron-dense core typical of LNPs containing nucleic acid cargos, with a high degree of blebbing (**Figure 4K, Figure S21**).

### Conjugate Treatment Activates On-Target Innate Immune Response *In Vivo*

We selected ovarian cancer as a proof of concept to apply the STING peptide-polyanion conjugate LNP as a therapeutic. There is a great clinical need to improve treatments for ovarian cancer, as a majority of patients are diagnosed after metastasis when existing therapeutics have limited benefit.^63,64^ T cell directed immunotherapies like checkpoint blockade have enabled durable tumor control in other metastatic cancers ^65^ but have thus far shown limited efficacy in ovarian cancer patients,^66,67^ correlated with poor T cell infiltration and the immunosuppressive microenvironments typical of ovarian cancer.^68,69^ STING signaling shows promise to repolarize immunosuppressive ovarian cancer tumors and promote anti-tumor immune responses. As STING expression is lost in over 70% of patients with metastatic disease,^16^ there may be a benefit of using STING mimicking therapy. We selected intraperitoneal (IP) administration for STING peptide-polyanion conjugate LNP therapy. This route of administration is used clinically for ovarian cancer treatment ^70,71^ and has been observed to significantly improve accumulation in abdominal tumors compared to intravenous administration.^70,72^

We first examined the pharmacokinetics of IP administered LNP in the BPPNM syngeneic mouse model of ovarian cancer. BPPNM mimics common mutations found in patients with homologous recombination-deficient high-grade serous ovarian carcinoma (*<u>B</u>rca1*^−/−^ *Tr<u>p</u>53*^−/−*R*172*H*^ *<u>P</u>ten*^−/−^ *<u>N</u>f1*^−/−^ *<u>M</u>yc*^OE^).^73^ This model recapitulates an immunosuppressive tumor microenvironment, resistance to checkpoint blockade therapies, and the abdominal metastasis pattern commonly observed in ovarian cancer patients.^73–75^ Tracking the fluorescence of Cyanine5-labeled peptide-polyanion conjugate, this therapy had a 2.5 h serum half-life that aligns with the half-life of other LNP platforms (**Figure 5A**).^76–78^ Next, biodistribution was measured at 4 h post IP administration via *ex vivo* IVIS. BPPNM tumor bioluminescence showed that the largest tumors were present on the omentum, with smaller tumor nodules on the upper genital tract (UGT) and intestines (**Figure S22**). The majority of measured peptide-polyanion conjugate fluorescence was observed in highly tumored organs, with 25%, 13%, and 25% in the omentum, UGT, and intestines, respectively (**Figure S22).** Primary sites of off-target accumulation were the liver and kidneys, with 14% and 10% of recovered conjugate fluorescence. Minimal (< 2%) accumulation was observed in the spleen, lungs, or heart (**Figure S22)**. To examine biodistribution at the cellular level, fluorescent signal from cells in the omental tumor and ascites (peritoneal fluid) was measured using flow cytometry (**Figure S23**). Conjugate association was expectedly high in the ascites as this fluid is the site of injection, with association observed in 56% of BPPNM cancer cells and 74% of stromal cells (CD45^-^, GFP^-^) (**Figure S23**). Conjugate association in the tumor was highest in BPPNM cancer cells (15%), stromal cells (47%), dendritic cells (DCs) (14%), and macrophages, with higher accumulation in M1-polarized (35%) compared to M2-polarized (9.1%) macrophages (**Figure S23**). Conjugate association with lymphoid cells was low overall, with the exception of B cells that showed moderate association in both the ascites (17%) and tumor (5.4%) (**Figure S23**).

**Figure 5:**
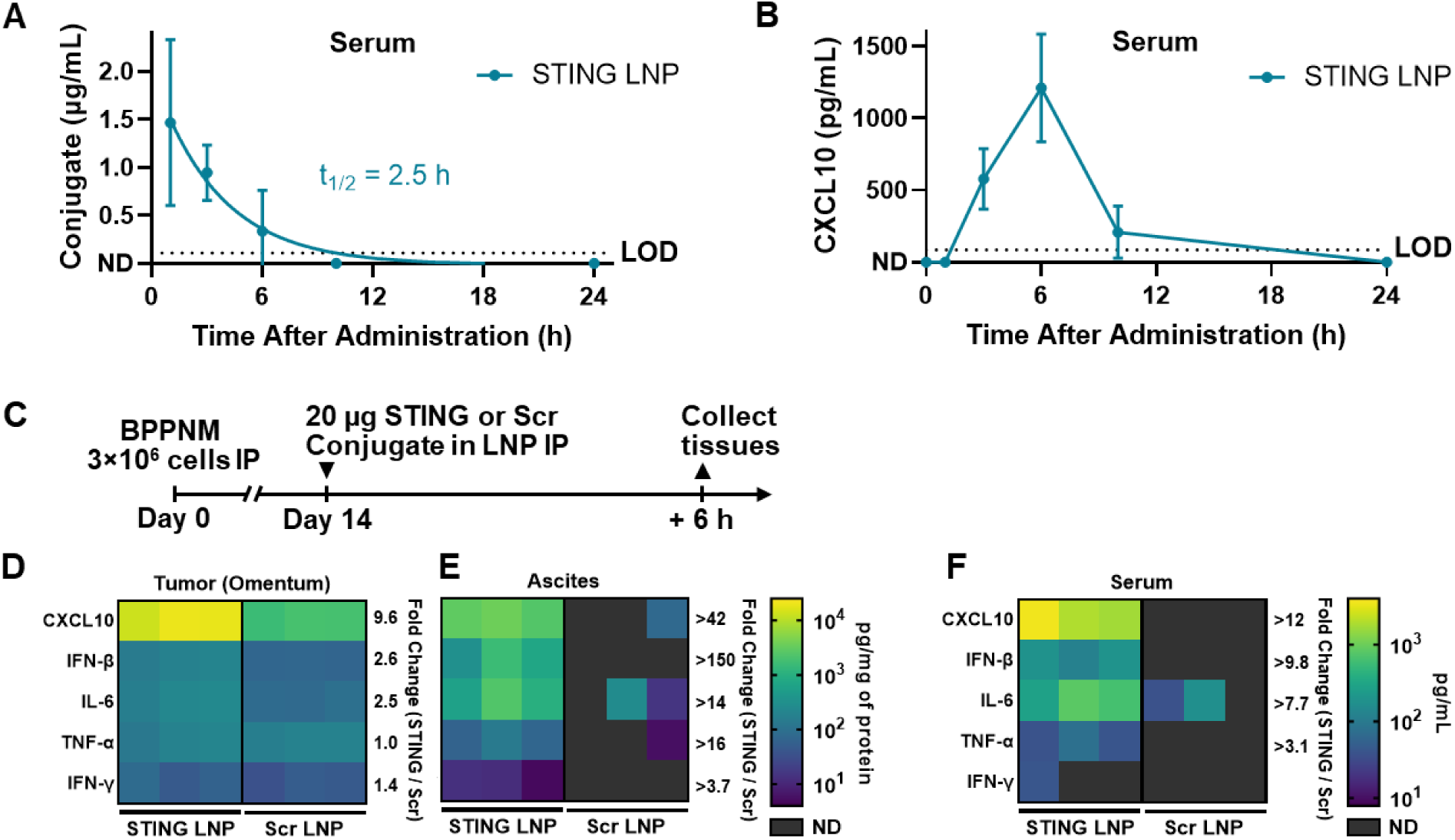
STING peptide-polyanion conjugate treatment induces innate immune cytokine production in mice. (**A-B**) Mice were dosed with 20 μg of STING conjugate delivered by LNP IP and serum was collected at 0, 1, 3, 6, 10, 24, and 50 h. Serum was analyzed to measure (**A**) STING conjugate concentration by Cy5 fluorescence (showing one phase exponential decay fit to data) and (**B**) CXCL10 concentration by ELISA (N = 3 mice). Conditions where analyte was below the limit of detection (LOD) are labeled as not detected (ND). (**C**) Mice were inoculated with 3×10^6^ BPPNM cells IP and dosed with 20 μg of STING or Scr conjugate (N = 3 mice) delivered by LNP IP at 14 days after inoculation. (**D**) Omental tumor, (**E**) ascites, and (**F**) serum were collected 6 h after dosing. Concentrations of CXCL10, IFN-β, IL-6, TNF-α, and IFN-γ were measured by ELISA and are reported relative to total protein concentration in tumor and ascites (**D-E**) or relative to volume in serum (**F**). Conditions where analyte was below the LOD are labeled as ND. Average fold change increases in cytokine concentration for STING conjugate treatment compared to Scr conjugate are displayed. Where cytokine level was ND, a lower bound on the fold change was computed by setting all ND replicates as the LOD. Data represented as mean ± SD.

On-target biological activity was validated *in vivo* by measuring the presence of IRF3 and NF-κB controlled cytokines. Serum levels of IRF3 controlled chemokine CXCL10 are below the ELISA assay limit of detection (LOD) at baseline, but rise to detectable levels between 3 and 10 h, hitting a maximum at 6 h after treatment with STING peptide-polyanion conjugate LNP (**Figure 5B**). Cytokine levels in an omental tumor, ascites, and serum were examined in more detail 6 h post treatment with STING or Scr peptide-polyanion conjugate (**Figure 5C, Figure S24**). BPPNM tumors had elevated baseline cytokine levels as observed in prior work;^73^ treatment resulted in significant increases in CXCL10 (9.6-fold), IFN-β (2.6-fold), and IL-6 (2.5-fold) (**Figure 5D**). In ascites, treatment resulted in large increases in CXCL10 (>42-fold), IFN-β (>150-fold), IL-6 (>14-fold), TNF-α (>16-fold), and IFN-γ (>3.7 fold) levels above the scramble control, where cytokine levels for the scramble control were below the LOD for many replicates or low otherwise (**Figure 5E**). Similar increases in cytokine levels were observed in the serum to a lesser magnitude, with CXCL10 (>12-fold), IFN-β (>9.8-fold), IL-6 (>7.7-fold) and TNF-α (>3.1-fold) levels increasing above the Scr control where most cytokines were below the LOD (**Figure 5F**).

### Conjugate Shows Therapeutic Efficacy in Metastatic Ovarian Cancer

With evidence that STING peptide-polyanion conjugate LNPs lead to innate immune activation, we examined therapeutic efficacy in the BPPNM model of metastatic ovarian cancer (**Figure 6A**). Treatment with STING peptide-polyanion conjugate LNP resulted in a significant survival improvement compared to controls (*P* = .006 vs PBS, *P* = .006 vs Scr LNP), increasing median survival to 37 days compared to 31 or 34 days for PBS or Scr controls, respectively (**Figure 6B**). Treatment resulted in a partial response with BPPNM luminescence shrinking over an order of magnitude; however, tumor growth resumed when treatment ended (**Figure 6C-E**). To examine the generalizability of this treatment, we examined efficacy in the KPCA.C syngeneic mouse model of metastatic ovarian cancer (**Figure 6F**), which mimics mutations found in a different subset of patients, homologous recombination-proficient high-grade serous ovarian carcinoma (*<u>K</u>RAS*^G12V^*Tr<u>p</u>53*^−/−R172H^*<u>C</u>cne1*^OE^*<u>A</u>kt2*^OE^).^73^ The KPCA.C model has a highly immunosuppressive tumor microenvironment, and is more resistant to immune checkpoint blockade than the BPPNM model.^73^ Treatment with STING peptide-polyanion conjugate LNP resulted in a significant survival improvement compared to controls (*P* = .02 vs PBS, *P* = .03 vs Scr LNP), increasing median survival to 33 days compared to 25 days for both PBS or Scr controls (**Figure 6G**). Treatment resulted in a partial response with KPCA.C luminescence shrinking over an order of magnitude; however, tumor growth resumed after treatment (**Figure 6H-J**). Observing therapeutic efficacy in two challenging to treat and immunosuppressive ovarian cancer models demonstrates the promise of STING peptide-polyanion conjugate as a standalone immunotherapy.

**Figure 6:**
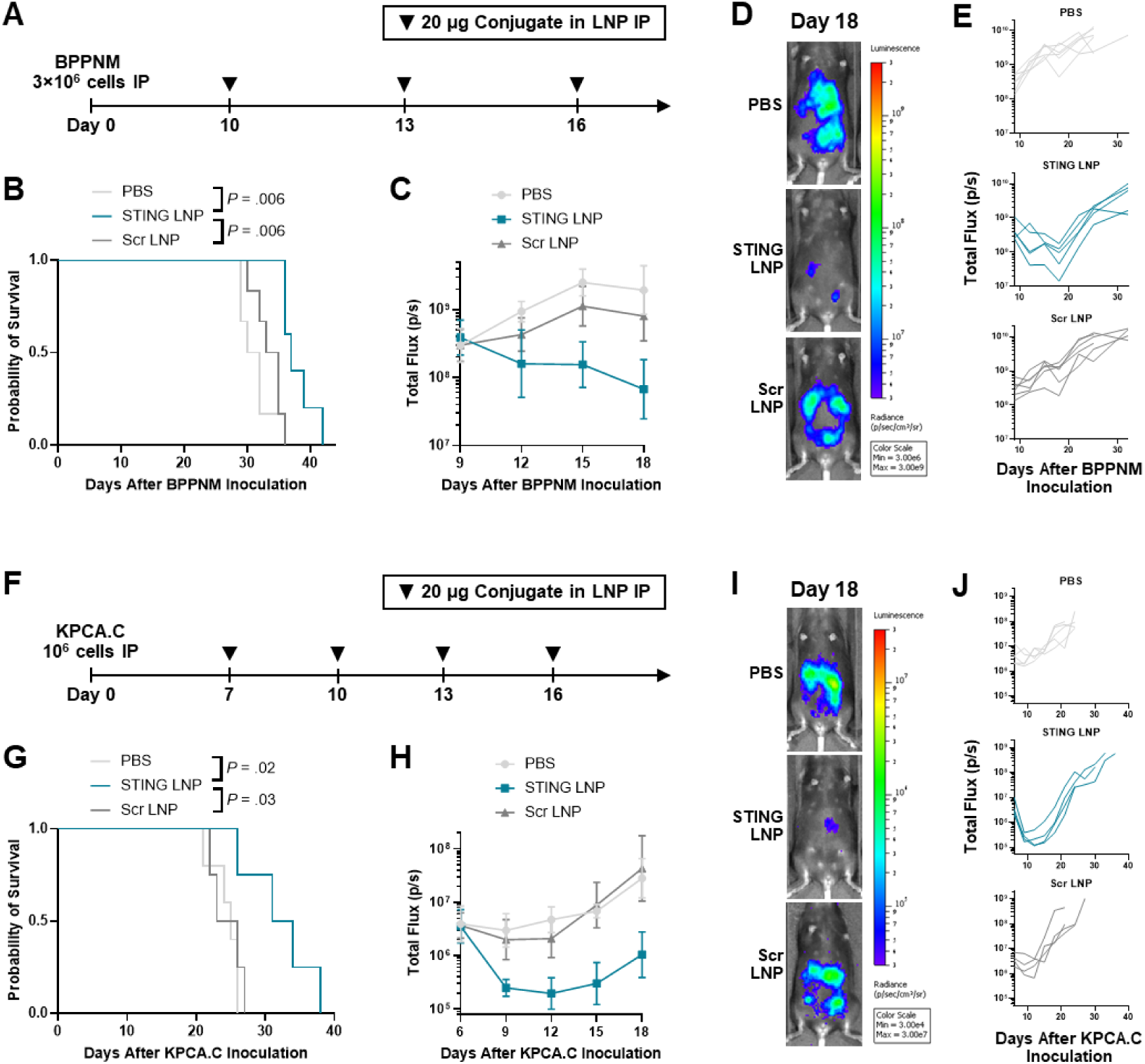
STING peptide-polyanion conjugate treatment shrinks tumors and prolongs survival in metastatic ovarian cancer models. (**A**) Mice were inoculated with 3×10^6^ BPPNM cells IP and dosed with 20 μg of STING or Scr conjugate delivered by LNP IP at 10, 13, and 16 days after inoculation. Groups included N = 6 (PBS, Scr LNP) or N = 5 (STING LNP) mice. (**B**) Survival plot with *P* values determined by log(rank) (Mantel–Cox) test. (**C-E**) Tumor burden measured by IVIS, displaying (**C**) geometric mean ± SD bioluminescent intensity over the treatment period, (**D**) an image of the mouse with the median bioluminescent intensity from each group at the measurement following the end of treatment (day 18), and (**E**) individual mouse bioluminescent intensity values. (**F**) Mice were inoculated with 10^6^ KPCA.C cells IP and dosed with 20 μg of STING or Scr conjugate delivered by LNP IP at 7, 10, 13, and 16 days after inoculation. Groups included N = 5 (PBS), or N = 4 (STING LNP, Scr LNP) mice. (**G**) Survival plot with *P* values determined by log(rank) (Mantel–Cox) test. (**H-J**) Tumor burden measured by IVIS, displaying (**H**) geometric mean ± SD bioluminescent intensity over the treatment period, (**I**) an image of the mouse with the median bioluminescent intensity from each group at the measurement following the end of treatment (day 18), and (**J**) individual mouse bioluminescent intensity values.

Administration of STING peptide-polyanion conjugate LNPs was well tolerated over the course of treatment of both BPPNM and KPCA.C models. Mice lost an average of 5-10% initial body mass in the day following treatment, but this effect was transient, and mice recovered to their initial mass within 1-2 days of each dose (**Figure S25**). Acute toxicity was examined by performing complete blood counts and a serum chemistry panel 24 h after a single dose of STING peptide-polyanion conjugate LNP or controls. Results were consistent with the known toxicity profiles of innate immune activation observed in previously examined STING activating therapies.^79–82^ A decrease in total white blood cell counts was observed, driven by decreases in blood lymphocytes and monocytes (**Figure S26**). A minor (<20%) but significant decrease in platelet counts was also observed (**Figure S26**); however, platelet counts remained within a normal range for mice and changes were small compared to those observed after systemic administration of a small molecule STING agonist.^81^ No significant changes in red blood cell counts or quality were observed with STING peptide-polyanion conjugate treatment (**Figure S26**). We also did not observe any signs of major liver or kidney damage, with no significant changes in liver enzymes, total protein, blood urea nitrogen, or creatinine (**Figure S27**).

### Conjugate Repolarizes the Tumor Microenvironment to Improve Response to Checkpoint Blockade

We evaluated the tumor immune response to the STING peptide-polyanion conjugate LNP in greater detail, focusing on the BPPNM model where there was more potential to improve therapeutic response. Four doses of STING peptide-polyanion conjugate LNP (**Figure 7A**) resulted in major changes in the immune composition of the tumor compared to PBS and Scr controls, as measured by flow cytometry (**Figure 7B, Figure S28-S29**). STING peptide-polyanion conjugate LNP treatment decreased the fraction of macrophages in the tumor, while increasing CD86 expression (**Figure 7C**) and decreasing CD206 expression (**Figure 7D**), indicating a shift from an immunosuppressive M2-phenotype to an inflammatory M1-phenotype. Treatment decreased the fraction of DCs in the tumor (**Figure 7B**), while increasing CD86 expression (**Figure 7E**), indicating DC activation. Treatment resulted in an increase in the fraction of CD4^+^ and CD8^+^ T cells present in the tumor (**Figure 7B**) but also increased PD-1 expression on both types of T cells (**Figure 7F, Figure S30**), providing evidence of T cell recruitment but suggesting that the PD-1 immune checkpoint may limit T cell effector function. Overall, these results demonstrate that STING peptide-polyanion conjugate LNP treatment can shift the BPPNM tumor from an immunosuppressive to an inflamed phenotype, both activating innate immune cells and recruiting T cells. We also observed changes that appeared to be caused by the presence of the LNP vehicle itself independently of its therapeutic cargo, that were observed with STING and Scr peptide-polyanion conjugate LNP treatment but not PBS control. LNP vehicle effects included an increase in the fraction of myeloid cells (CD11b^+^) in the ascites (**Figure S31**) and MDSC (CD11b^+^ Gr1^hi^) recruitment to the tumor (**Figure 7B**). Myeloid recruitment is a previously demonstrated outcome of LNP administration,^61^ caused by the immunogenic nature of the LNP ionizable lipid and endosomal escape.^58–61^ The presence of lipid particles has also previously been shown to induce MDSC-like cell populations.^61,83^

**Figure 7:**
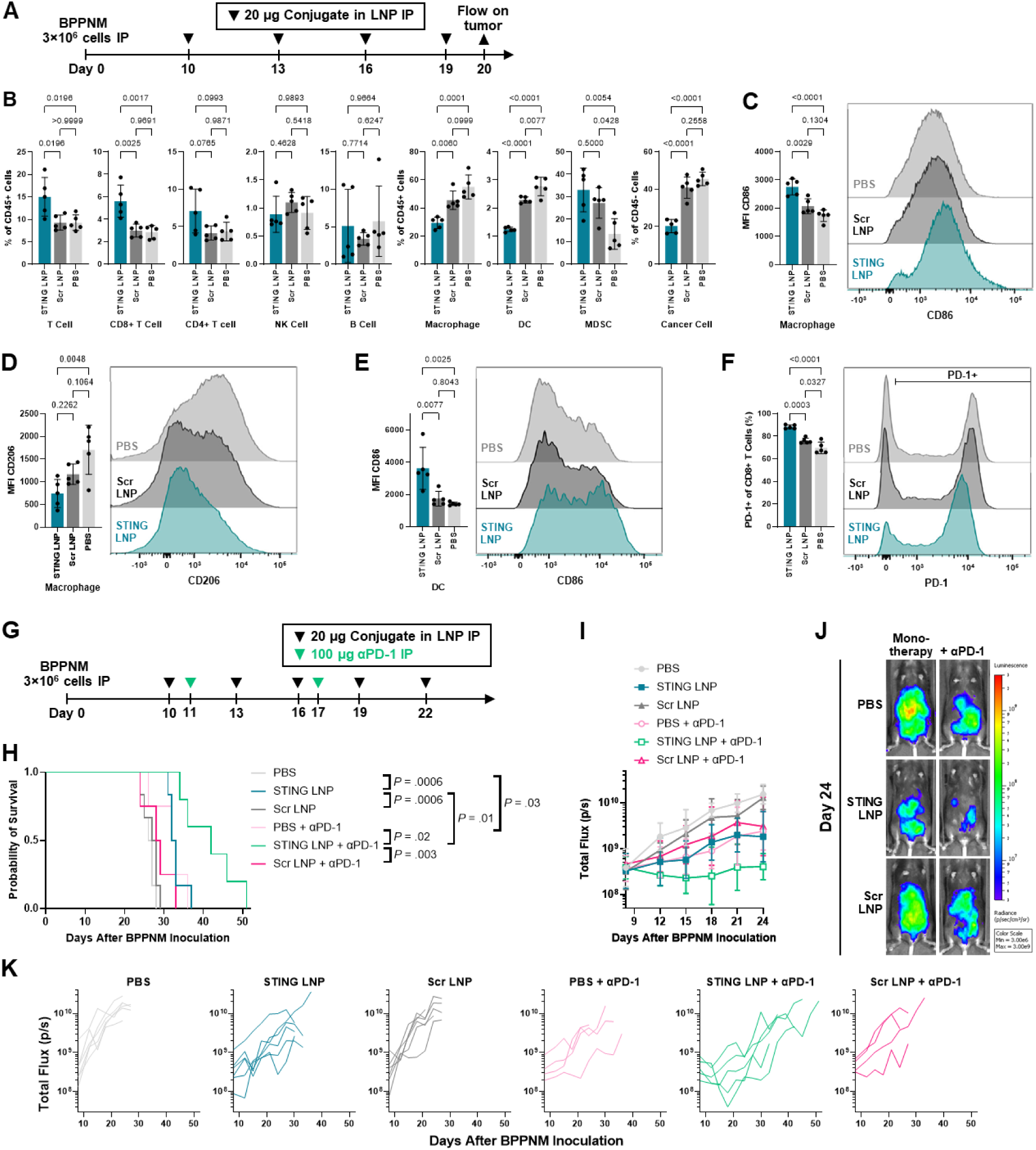
STING peptide-polyanion conjugate treatment repolarizes the ovarian tumor microenvironment to improve response to PD-1 checkpoint blockade. (**A**) Mice were inoculated with 3×10^6^ BPPNM cells IP and dosed with 20 μg of STING or Scr conjugate delivered by LNP IP at 10, 13, 16, and 19 days after inoculation. Omental tumor was collected on day 20 for analysis by flow cytometry. Groups included N = 5 mice. (**B**) Cell populations in tumor after treatment, displaying percentage of CD45^+^ cells made up by T cells (CD3^+^), CD8^+^ T cells, CD4^+^ T cells, NK cells (CD3^-^ NK1.1^+^), B cells (CD19^+^), Macrophages (F4/80^+^), DCs (CD11c^+^ MHCII^+^), and MDSCs (CD11b^+^ Gr-1^hi^). Percentage of CD45^-^ cells made up by BPPNM cancer cells (GFP^+^) are also displayed. (**C-D**) Polarization of Macrophages (F4/80^+^), showing MFI and representative distributions of (**C**) CD86 and (**D**) CD206. (**E**) Activation of DCs (CD11c^+^ MHCII^+^), showing MFI and representative distributions of CD86. (**F**) Expression of PD-1 on CD8^+^ T cells, showing percentage of cells that are PD-1^+^ and representative distributions of PD-1. *P* values computed with a one-way ANOVA followed by Tukey’s post-hoc test are displayed above each figure. All flow data is represented as mean ± SD. (**G**) Mice were inoculated with 3×10^6^ BPPNM cells IP and dosed with 20 μg of STING or Scr conjugate delivered by LNP IP at 10, 13, 16, 19, and 22 days after inoculation. A subset of groups were additionally treated with 100 μg αPD-1 antibody at 11 and 17 days after inoculation. Groups included N = 6 (PBS, STING LNP, Scr LNP), N = 5 (STING LNP + αPD-1), or N = 4 (PBS + αPD-1, Scr LNP + αPD-1) mice. (**H**) Survival plot with *P* values determined by log(rank) (Mantel–Cox) test. (**I-K**) Tumor burden measured by IVIS, displaying (**I**) geometric mean ± SD bioluminescent intensity over the treatment period, (**J**) an image of the mouse with the median bioluminescent intensity from each group at the measurement following the end of treatment (day 24), and (**K**) individual mouse bioluminescent intensity values.

Combination therapy with PD-1 checkpoint blockade was examined to leverage the T cell inflamed tumor microenvironment and overcome PD-1 expression induced by STING peptide-polyanion conjugate LNP treatment, using five doses of LNP with anti-PD-1 antibody (αPD-1) administered IP 1 day after the first and third dose (**Figure 7G**). As observed in prior work in this model,^73^ αPD-1 treatment alone resulted in a modest survival improvement compared to PBS control (*P* = .03), increasing median survival to 33 days compared to 27 days for the PBS control (**Figure 7H**). Combination of STING peptide-polyanion conjugate LNP with anti-PD-1 treatment resulted in a significant survival improvement compared to controls (*P* = .02 vs PBS + αPD-1, *P* = .003 vs Scr LNP + αPD-1), increasing median survival to 42 days compared to 33 or 29 days for PBS + αPD-1 or Scr + αPD-1 controls, respectively (**Figure 7H**). Combination treatment resulted in a partial response with a reduction in BPPNM luminescence that remained low over the course of treatment, however, tumor growth resumed after treatment ended (**Figure 7I-K**). Combination therapy was also well tolerated, with mice experiencing only small (<5-10%) and transient (1-2 days) drops in mass after each treatment, similar to STING peptide-polyanion conjugate monotherapy (**Figure S32**). These results show that STING peptide-polyanion conjugate LNP treatment complements PD-1 checkpoint blockade to further improve therapeutic outcomes beyond either treatment as a monotherapy.

## DISCUSSION

This work demonstrates that the design of STING mimicking therapeutics can be greatly simplified while maintaining strong therapeutic efficacy. High valency display of ligands with affinity for TBK1 and IRF3 in the cytosol is sufficient to activate IRF3. We were able to achieve this with a 39-amino acid peptide that contains short linear TBK1 and IRF3 interaction motifs, a peptide that is only 10% the length of the 378 amino acid STING protein or only 16% the length of the approximately 241 amino acid STING cytosolic domain commonly used in other STING mimicking theraputics.^43,47^ We anticipate that design lessons from this work for the simplification of STING mimics will be valuable for the design of next generation STING mimicking therapeutics as a whole, beyond just peptide conjugate systems. The STING peptide-polyanion conjugate activated IRF3 and NF-κB transcription factors at a similar ratio to a traditional STING agonist; however, a previously developed STING mimic reported a strong preference for IRF3 activation over NF-κB.^47^ We did not examine this discrepancy further, but it suggests that changing some aspect of STING mimic design may allow for tuning relative levels of IRF3 and NF-κB activation.

We designed our therapeutic molecule from the ground up with nanocarrier integration in mind, using a polyanionic backbone to promote encapsulation by existing nanocarriers that use cationic moieties to encapsulate negatively charged cargos. This greatly accelerated the development of our delivery system compared to prior work delivering the STING mimicking protein therapeutic STINGΔTM to the cytosol.^44^ Using the same HEK293T reporter-based functional delivery screen, STINGΔTM protein delivery was only effective with a fraction of tested vehicles and had poor functional delivery efficacy,^44^ while peptide-polyanion conjugate delivery in this work resulted in strong functional delivery efficacy with all vehicles tested. We only needed to perform a single round of LNP optimization to develop a formulation of small, monodisperse LNPs with high encapsulation efficiency and cytosolic delivery efficiency, that were ultimately effective for delivery of this novel peptide-polyanion conjugate cargo *in vivo*. Still, this screen revealed some differences in LNP formulation design rules when delivering peptide-polyanion conjugate compared to mRNA or siRNA, likely because of the high mass of hydrophilic and neutrally charged peptide that must also be encapsulated. Notably, a higher than typical proportion of phospholipid and lower proportion of cholesterol were required to achieve a formulation that retained a small and stable size as well as high encapsulation efficiency.

We demonstrated that the STING peptide-polyanion conjugate has promise as a therapeutic, initiating an on-target immune response in the tumor and improving survival in multiple challenging to treat models of metastatic ovarian cancer. This therapeutic efficacy was achieved with a proof-of-concept for a first-generation peptide-conjugate STING mimic. We anticipate that there is significant space for optimization of this system, including in the design of peptide-polyanion conjugate molecule, optimization of the LNP formulation, and introduction of targeting ligands. Finally, the availability of potent STING mimicking therapeutics will allow for STING activation in cancer cells to be explored as a therapeutic strategy, building on the large body of work focused on STING activation in immune cells in the STING agonist delivery field.^20,21^ There is certainly evidence that STING activation in cancer cells is beneficial to strengthen an anti-cancer immune response;^22–28^ however, more work is needed to show that STING activation in cancer cells leads to broad therapeutic benefit beyond existing evidence in a handful of animal models.^22^ Direct comparison of the therapeutic efficacies of STING agonists to STING mimics in a matched vehicle to equalize pharmacokinetics in animal models that match the epigenetic STING silencing found in human cancer would yield valuable insights to address this question. Regardless, STING peptide-polyanion conjugate is also effective in cells with intact STING signaling like KURAMOCHI or THP-1, suggesting that this STING mimicking therapy is capable of activating downstream signaling in host immune and stromal cells when needed.

## CONCLUSION

We demonstrated that the signaling biology of the activated STING-TBK1-IRF3 complex can be mimicked by display of short, linear protein interaction motifs on a multivalent polymer conjugate. This platform allowed us to therapeutically activate the innate immune transcription factors IRF3 and NF-κB, even in cell lines that are unresponsive to STING agonists. The peptide-polyanion conjugate could easily be loaded in a LNP carrier that allowed for its application as an immunotherapy and demonstrated efficacy in multiple mouse models of metastatic ovarian cancer. This work provides evidence that polyanion conjugates are a versatile tool to enable the delivery of therapeutic cargo into cells. The delivery of high valency protein-protein interaction motifs to the cytosol has potential to engage cellular signaling machinery in ways that are challenging with existing therapeutic options; for this reason, this work inspires the use of this strategy beyond applications in STING mimicking therapeutics.

## METHODS

### Materials

Poly(L-glutamic acid sodium salt, MW = 45,000 Da) 10% graft alkyne (PLE_300_-g-AK_10%_, #000-E300-g-AK010) and (poly-L-glutamic acid sodium salt, MW = 45,000 Da) with an alkyne function on the C-terminal (AK-PLE_300_, #000-AKE300) were purchased from Alamanda Polymers. Molecular weights and alkyne substitution from the lot specific specification sheet were used for all calculations. PLE_300_-g-AK_10%_ Lot #000-E300-g-AK010-101: number average molecular weight (M_n_) by NMR = 47,600 Da, number average degree of polymerization by NMR = 312, polydispersity index (PDI) by GPC = 1.02, alkyne functionalization by NMR = 11%. AK-PLE_300_ Lot #000-AKE300-101: M_n_ by NMR = 48,600 Da, DP_n_ by NMR = 320, PDI by GPC = 1.01, alkynyl substitution by NMR = 100%.

1,2-Dioleoyl-sn-glycero-3-phosphoethanolamine (DOPE), 1,2-distearoyl-sn-glycero-3-phosphocholine (DSPC), 1,2-dioleoyl-sn-glycero-3-phosphocholine (DOPC), cholesterol (ovine), and 6-((2-hexyldecanoyl)oxy)-N-(6-((2-hexyldecanoyl)oxy)hexyl)-N-(4-hydroxybutyl)hexan-1-aminium (ALC-0135) were purchased from Avanti Research. 8-[(2-hydroxyethyl)[6-oxo-6-(undecyloxy)hexyl]amino]-octanoic acid, 1-octylnonyl ester (SM-102) and 1,2-dimyristoyl-rac-glycero-3-methoxypolyethylene glycol-2000 (DMG-PEG2000) were purchased from Cayman Chemical Company. Dilinoleylmethyl-4-dimethylaminobutyrate (MC3) and 2’3’-c-di-AM(PS)2 (Rp,Rp) disodium salt (ADU-S100 disodium salt) were purchased from MedChemExpress. MRT67307 was purchased from InvivoGen. Sulfo-Cyanine5 azide was purchased from Lumiprobe. Copper(II) sulfate pentahydrate, (+)-sodium L-ascorbate, aminoguanidine hydrochloride, glycerol, 3M sodium acetate solution (pH 5.2), Tween 20, Triton X-100, 0.4% Trypan Blue solution, 0.01% poly-l-lysine solution, bovine serum albumin (BSA), Collagenase from Clostridium histolyticum Type V, DNase I, and Hyaluronidase Type I-S were purchased from Sigma-Aldrich. Sodium chloride and absolute (200 proof) ethanol were purchased from Fisher Scientific. Tris-hydroxypropyltriazolylmethylamine (THPTA) was purchased from Vector Laboratories. Tris-HCl was purchased from G-Biosciences. 1M sodium acetate solution (pH 4.5) and 16% Formaldehyde (w/v), methanol-free was purchased from Thermo Scientific. Opti-MEM, Fetal bovine serum (FBS), 5000 U/mL penicillin and 5000 µg/mL streptomycin solution, 0.25% Trypsin-EDTA, TrypLE Express Enzyme (1×), and ACK Lysis Buffer were purchased from Gibco. Dulbecco′s Modified Eagle′s Medium (DMEM), Roswell Park Memorial Institute (RPMI) 1640, and 1× Phosphate Buffered Saline (1× PBS) was purchased from Corning. DMEM/F12 1:1 Media was purchased from Cytiva. 0.5 M ethylenediaminetetraacetic acid (EDTA) (pH 8.0) was purchased from Growcells. Type I Ultrapure water generated with a Milli-Q IQ 7000 Ultrapure Lab Water System equipped with a Biopak polisher (Millipore) was used for all experiments.

### Conjugate Synthesis and Characterization

#### Peptide Design and Synthesis

The STING C-terminal tail peptide was based on mouse STING(340-378) (Uniprot Q3TBT3): VTMNAPMTSV APPPSVLSQE PRLLISGMDQ PLPLRTDLI. The scramble (Scr) control peptide was randomly rearranged: LLRNRPVTSG ESMPVPISQA PADLLLPPLQ MMPVSTDTI. An azidolysine residue was included on the N-terminus of both peptides. Peptides were custom synthesized and characterized by GenScript Biotech. Peptide molecular weight was consistent with the theoretical molecular weight of 4341.13 Da by electrospray ionization (ESI) mass spectrometry (MS). Purity was ≥98% by HPLC. Endotoxin levels were <10 EU/mg by Limulus Amebocyte Lysate (LAL) assay.

#### Fluorescent labeling of poly(L-glutamate) graft alkyne

Poly(L-glutamate)-graft-alkyne was modified with sulfo-cyanine5 azide using copper catalyzed azide–alkyne cycloaddition. The reaction was performed in water by adding sulfo-cyanine5 azide and poly(L-glutamate)-graft-alkyne to a mixture with a final concentration of 0.1 mM Poly(L-glutamate)-graft-alkyne, 1 mM copper (II) sulfate, 10 mM sodium ascorbate, and 1 mM THPTA. Sulfo-cyanine5 azide was added at 2 molar equivalents to poly(L-glutamate)-graft-alkyne for all experiments except for confocal microscopy, where a highly-labeled batch was prepared with 10 molar equivalents of sulfo-cyanine5 azide. The reaction was carried out at room temperature (RT) protected from light for 1 hour and then at 4 °C overnight. The reaction mixture was dialyzed (3.5 kDa MWCO Slide-A-Lyzer Dialysis Cassette) against 5 mM EDTA, followed by 50 mM sodium chloride, followed by water and then lyophilized (Labconco FreeZone Freeze Dryer, <-85 °C and <0.3 mbar) to generate a blue solid.

The removal of unreacted fluorophore was confirmed with thin layer chromatography (TLC) using silica gel 60 F_254_ (Sigma) as the stationary phase and methanol as the mobile phase. Unreacted sulfo-cyanine5 azide migrated near the solvent front, while the purified sulfo-cyanine5 azide-labeled poly(L-glutamate)-graft-alkyne did not migrate and showed no signs of unreacted dye. Fluorescent tag incorporation was quantified by comparing sulfo-cyanine5 absorbance at 646 nm (ε_dye_ = 271,000 M^−1^ cm^−1^, reported by vendor) to poly(L-glutamate)-graft-alkyne absorbance at 205 nm (ε_PLE300-g-AK10%_ = 1,020,000 M^−1^ cm^−1^, measured for stock solution), correcting for dye absorbance at 205 nm using the correction factor (CF_205_ = 0.14, measured for stock solution), using a NanoDrop One^C^ ultraviolet–visible (UV–vis) spectrophotometer (Thermo Fisher).

#### Multivalent peptide poly(L-glutamate) conjugate synthesis

Sulfo-cyanine5-labeled poly(L-glutamate)-graft-alkyne was modified with azidolysine containing peptides using copper catalyzed azide–alkyne cycloaddition. The reaction was performed in water by adding azidolysine containing peptide and poly(L-glutamate)-graft-alkyne to a mixture with a final concentration of 1 mM alkyne, 1.2 mM azide, 1 mM copper (II) sulfate, 10 mM sodium ascorbate, 1 mM THPTA, and 10 mM aminoguanidine. The reaction was carried out at RT protected from light for 1 hour and then at 4 °C overnight. The conjugate was purified by repeated centrifugal filtration (10 kDa Amicon Ultra Centrifugal Filter) at 1500×g, washing first with 5 mM EDTA, followed by 50 mM sodium chloride, and then water to reach a total dilution factor of >1,000,000. Conjugate solution in water was sterilized with a 0.2 μm cellulose acetate filter and stored at −80 °C until use.

Conjugate was quantified by measuring sulfo-cyanine5 absorbance of the labeled poly(L-glutamic acid) backbone at 646 nm (ε_dye_ = 271,000 M^−1^ cm^−1^) using a NanoDrop One^C^ ultraviolet–visible (UV–vis) spectrophotometer. Complete reaction was confirmed by measuring the disappearance of detectable alkynes using a 3-azido-7-hydroxycoumarin-based alkyne quantification kit (ProteinMods) following vendor directions. NMR (Bruker AVANCE, 500 MHz ^1^H, D_2_O) was used to confirm the disappearance of the poly(L-glutamate)-graft-alkyne alkyne-adjacent methylene δ 3.95 (-C**H2**-C≡CH) protons. Successful reaction was also confirmed by the appearance of a triazole δ 7.89 proton in the conjugate. Removal of unreacted excess peptide was confirmed by size-exclusion chromatography (Zorbax GF-250 column (Agilent) with 1× PBS mobile phase). Endotoxin levels were quantified with the HEK-Blue mTLR4 reporter cell assay (InvivoGen) following vendor instructions, with all conjugates having <2 EU/mg.

#### Monovalent peptide poly(L-glutamate) conjugate synthesis

End-functionalized alkyne-poly(L-glutamate) was modified with azidolysine containing peptides using copper catalyzed azide–alkyne cycloaddition. The reaction was performed in water by adding azidolysine containing peptide and alkyne-poly(L-glutamate) to a mixture with a final concentration of 0.1 mM alkyne, 0.12 mM azide, 1 mM copper (II) sulfate, 10 mM sodium ascorbate, 1 mM THPTA, and 10 mM aminoguanidine. The reaction was carried out at RT protected from light for 1 hour and then at 4 °C overnight. The conjugate was purified by repeated centrifugal filtration (3 kDa Amicon Ultra Centrifugal Filter) at 12,000×g, washing first with 5 mM EDTA, followed by 50 mM sodium chloride, and then water to reach a total dilution factor of >1,000,000. Conjugate solution in water was sterilized with a 0.2 μm cellulose acetate filter and stored at −80 °C until use. Conjugate was quantified by measuring peptide backbone absorbance at 205 nm (ε = 1,000,000 M^−1^ cm^−1^, estimated based on amino acid composition^84^) using a NanoDrop One^C^ ultraviolet–visible (UV–vis) spectrophotometer. NMR (Bruker AVANCE, 500 MHz ^1^H, D_2_O) was used to confirm the disappearance of the alkyne-poly(L-glutamate) alkyne-adjacent methylene δ 4.24 (-C**H2**-C≡CH) protons.

#### Circular Dichroism

Samples of azido lysine-mSTING(340-378) peptide (25 µg/mL), poly(L-glutamate)-graft-alkyne polymer (50 µg/mL), and a multivalent poly(L-glutamate)-graft**-**mSTING(340-378) conjugate (50 µg/mL) were prepared in 1× PBS. Circular dichroism (CD) spectra were obtained with a Jasco J-1500 CD Spectropolarimeter in the Biophysical Instrumentation Facility at MIT. Samples (250 μL) were loaded into a 1 mm quartz cuvette and analyzed at ambient temperature. Spectra were collected from 190 to 250 nm with a scan speed of 50 nm/min, a digital integration time of 4 s, a data pitch of 0.5 nm, and a bandwidth of 1 nm. Reported spectra are the average of 3 measurements. To process data, signal from a 1× PBS blank was subtracted and then raw millidegrees (mdeg) of rotation (m°) were converted to mean residual ellipticity (MRE) using the equation MRE = m° × M / (10 × L × C × R), where M is the molar mass of the construct in g/mol, L is the pathlength of the cuvette in cm, C is the concentration of the construct in mg/mL, and R is the average number of amino acid residues in each construct.

### LNP Formation and Characterization

#### LNP Formation

DOPC was dried from a chloroform stock under nitrogen and then dissolved in 100% ethanol. ALC-0305, MC3, DOPE, DSPC, Cholesterol, and DMG-PEG2000 were directly dissolved in 100% ethanol.

LNPs were formed by bulk mixing using a previously described method.^85^ LNPs were prepared with varying lipid components and compositions as described in **Table S1**. Peptide-polyanion conjugate was diluted into 25 mM sodium acetate at pH 4.5 in a glass scintillation vial charged with a polytetrafluoroethylene (PTFE) stir bar. To 4 volumes of peptide-polyanion conjugate under magnetic stirring (700 rpm), 1 volume of lipid mixture in 100% ethanol was pipetted in rapidly. The solution was allowed to mix for 10 s, then rested without stirring for 5 min. Magnetic stirring was resumed (700 rpm), the solution was diluted with 5 volumes of water and allowed to mix for 10 s. LNPs were purified and concentrated by repeated centrifugal filtration (100 kDa Amicon Ultra Centrifugal Filter) at 2,650×g, washing with water. LNPs were diluted to reach a final concentration of 1× PBS. LNP solutions were stored at 4 °C until use.

#### Dynamic Light Scattering and Zeta Potential Measurement

The Z-average diameter and polydispersity index (PDI) of the nanoparticles were characterized by dynamic light scattering (DLS) and the zeta potential by laser doppler anemometry, using a Malvern Zetasizer Pro Red (Malvern Panalytical). The measurements were performed at 25 °C with a red laser (wavelength = 633 nm) and a detection angle of 173°. Each data point reported is the mean of technical replicates for an independently assembled nanoparticle formulation.

#### Conjugate Quantification in LNP

Peptide-polyanion conjugate concentration in purified LNP samples was quantified by measuring the fluorescence intensity of the Cyanine5-labeled polyanion backbone compared to a peptide-polyanion conjugate standard at a known concentration. LNPs were lysed by adding 1 volume of 4% (v/v) Trition X-100 in water to 1 volume of LNP or standard sample, followed by 10 min incubation at 37 °C with orbital shaking. Fluorescence intensity was read with an Infinite MPlex Plate Reader (Tecan), with excitation at 640 nm and emission at 680 nm.

#### Encapsulation Efficiency Assay

Peptide-polyanion conjugate encapsulation in LNPs was measured using native polyacrylamide gel electrophoresis (PAGE), where soluble conjugate can migrate freely into the gel but encapsulated conjugate remains trapped in the well. Each LNP sample was divided into two containers to allow for direct comparison of intact to lysed LNPs. To 1 volume of LNP sample, 1 volume of water (intact condition) or 1 volume of 2% (v/v) Triton X-100 (lysed condition) was added. Samples were incubated at 37 °C for 10 min with orbital shaking. Each sample was then mixed with 2 additional volumes of Native PAGE Sample Buffer (62.5 mM Tris-HCl, pH 6.8, 40% (v/v) glycerol), mixed thoroughly by pipette, and loaded onto a gel (Mini-PROTEAN TGX Precast Protein gel, Bio-Rad). Native PAGE was run in 25 mM Tris, 192 mM glycine, pH 8.3 buffer at 100 V for 1 h using a Mini-PROTEAN Tetra Cell (Bio-Rad) and a PowerPac™ HC High-Current Power Supply (Bio-Rad). Gels were imaged using the Cyanine5 method to detect Cyanine5-labeled polyanion backbone using a ChemiDoc MP imaging system (Bio-Rad). Image Lab software (Bio-Rad) was used to quantify the Cyanine5 fluorescence signal from free peptide-polyanion conjugate in each lane using local background subtraction. Encapsulation efficiency (EE%) was calculated by comparing the fluorescence signal of free conjugate in lysed LNPs (F_lysed_) to intact LNPs (F_inact_) using the following formula:

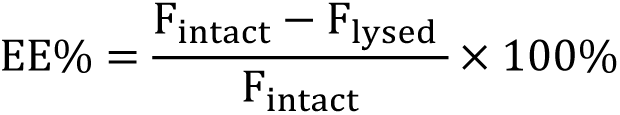

#### Effective pKa Assay

Solutions of 20 mM sodium phosphate, 20 mM ammonium acetate, 25 mM citrate, and 150 mM sodium chloride were titrated to pH values of 2.0, 3.0, 3.5, 4.0, 4.5, 5.0, 5.5, 6.0, 6.5, 7.0, 7.5, 8.0, 8.5, 9.5, 10.0, and 11.0. 2-p-Toluidinonaphthalene-6-sulfonic acid (TNS) was dissolved in DMSO to a working concentration of 300 μM. In a black 96-well plate, LNPs containing ALC-0315 were incubated in the buffer solutions described above. TNS was added to each well such that the final concentration of LNP ionizable lipid was 20 μM and TNS was 6 μM at a total volume of 100 μL. The plate was covered and placed on an orbital shaker at 300 rpm for 1 min, then fluorescence intensity was read with an Infinite MPlex Plate Reader, with excitation at 322 nm and emission at 431 nm. The pK_a_ was determined by fitting a four-parameter logistic curve to the data using GraphPad Prism.

#### Transmission Electron Microscopy

For cryo-transmission electron microscopy (Cryo-TEM), 3 µL of the nanoparticle sample (exchanged into water using 100 kDa MWCO Amicon centrifugal filter at 2650×g) was applied to copper grids coated with a continuous carbon film. The grids were pre-treated with oxygen plasma using a Solarus 950 Gatan Advanced Plasma System. Excess sample on the grid was gently blotted using the Gatan Cryo Plunge III, followed by rapid plunging into liquid ethane to vitrify the sample. The grid was then mounted on a Gatan 626 single tilt cryo-holder, which was subsequently inserted into the TEM column. Both the specimen and the holder tip were maintained at cryogenic temperatures with liquid nitrogen to ensure preservation throughout the transfer and imaging process. Imaging was performed on a JEOL 2100 FEG microscope using a minimum dose method to reduce electron beam damage to the sample. The microscope was operated at 200 kV, with magnifications between 10,000× and 60,000× to evaluate particle size and distribution. All images were captured using a Gatan 2k × 2k UltraScan CCD camera.

### *In Vitro* Experiments

#### Cell Lines

HEK293T pGL4.45 cells are a reporter derivative of HEK293T (ATCC CRL-3216) generated previously in our lab.^43^ Briefly, HEK293T were stably transfected with the pGL4.45[*luc2P/ISRE/Hygro*] vector (Promega) to allow for luciferase induction in response to IRF3 activation. THP1-Dual (thpd-nfis) and HEK-Blue mTLR4 (hkb-mtlr4) reporter cells were purchased from InvivoGen. KURAMOCHI mCherry^+^ Luciferase^+^ cells were a gift from the laboratory of Ronny Drapkin (University of Pennsylvania). A2780 cells were a gift from the laboratory of Stephen Howell (UC San Diego). BPPNM and KPCA.C cells were a gift from the laboratory of Robert Weinberg (MIT).^73^ BPPNM cells stably expressing luciferase, a gift from the laboratory of Stefani Spranger (MIT), were generated by lentiviral transduction of a plasmid containing the sequence of firefly luciferase cloned into the backbone vector pLV-EF1a-IRES-Blast (Addgene plasmid #85133) and selected using 15 µg/mL blasticidin. KPCA.C cells stably expressing mCherry-SIY and luciferase, a gift from the laboratory of Stefani Spranger (MIT), were generated by lentiviral transduction of a plasmid containing the sequence mCherry-SIY-P2A-Luciferase (mCherry with the SIY antigen fused to its C-terminus, followed by the “self-cleaving” peptide P2A, then firefly luciferase) was cloned into the backbone vector pLV-Ef1a-IRES-Puro (Addgene plasmid #85132) and selected using 2.5 µg/mL puromycin.

#### Cell Maintenance

HEK293T pGL4.45 were cultured in DMEM supplemented with 10% FBS, 50 U/mL penicillin, and 50 µg/mL streptomycin (P/S). HEK-Blue mTLR4 cells were cultured in DMEM supplemented with 10% heat inactivated (56 °C for 30 min) FBS and P/S. A2780 cells were cultured in RPMI 1640 supplemented with 10% FBS and P/S. THP1-Dual were cultured in RPMI 1640 supplemented with 10% heat inactivated (56 °C for 30 min) FBS and P/S. KURAMOCHI were cultured in DMEM/ F12 1:1 supplemented with 10% FBS and P/S. BPPNM and KPCA.C cells were cultured in DMEM supplemented with 4% heat inactivated FBS (Millipore Sigma #F4135), 1% insulin–transferrin–selenium (Thermo Fisher #41400045), and P/S. Adherent cells were dissociated by tapping the flask (HEK293T pGL4.45 and HEK-Blue mTLR4) or by incubation in 0.25% Trypsin-EDTA (KURAMOCHI, A2780, BPPNM, and KPCA.C). All cells were incubated at 37 °C at 100% humidity and 5% CO_2_ atmosphere for maintenance and during all assay incubation periods unless otherwise specified. Cells were used at less than 20 passages. Live cells were counted using a Trypan Blue stain with a Cellometer Auto 1000 (Nexcelom). Cell lines were tested upon receipt and routinely during culture for mycoplasma at the Koch Institute’s Preclinical Modeling Core Facility (MIT) using the MycoAlert PLUS Assay (Lonza Biosciences). All cell lines tested negative. All cell lines were tested using the short tandem repeat (STR) profiling service at American Type Culture Collection, with the exception of commercial reporter cell lines (HEK-Blue mTLR4, THP1-Dual) which were used as received from vendors.

#### Transfection Complex Preparation

TransIT-X2 Dynamic Delivery System (Mirus Bio) is a polymer-based delivery system that was used for routine transfection of peptide-polyanion conjugate following vendor directions. TransIT-X2 was warmed to RT and vortexed gently, then added to the cargo in Opti-MEM at a ratio of 1 μL TransIT-X2 per μg of cargo, with thorough pipet mixing. The mixture was incubated at RT for 15 to 30 min to allow for complex formation and then added to cells. Transfection grade, linear, MW 25,000, polyethylenimine (PEI) (Polysciences) is a polymeric transfection reagent. PEI was dissolved in 25 mM sodium acetate at pH 5.2 at a concentration to allow 2 μg of PEI per μg of cargo. The cargo was also diluted in 25 mM sodium acetate at pH 5.2, and then mixed with the PEI solution at a 1:1 volume ratio. The mixture was incubated at RT for 15 min, then added to cells. The PBAE Poly2 is a polymeric transfection reagent. Poly2 from the same batch of polymer synthesized and characterized in previous work was used.^56^ Poly2 was dissolved in 25 mM sodium acetate at pH 5.2 at a concentration to allow 8 μg of Poly2 per μg of cargo. The cargo was also diluted in 25 mM sodium acetate at pH 5.2, and then mixed with the Poly2 solution at a 1:1 volume ratio. The mixture was incubated at RT for 15 min, then added to cells.

#### Reporter Cell Assays

Assays with HEK293T pGL4.45 reporter cells were performed as described previously.^43,86^ Briefly, 3×10^5^ cells/mL were added to a 96-well plate in 100 μL of DMEM with 10% FBS and P/S, then allowed to incubate for 24 h. To each well, 20 μL of treatment specified in each figure caption was added, then allowed to incubate for 24 h. Luciferase was quantified using the Firefly Luciferase Assay Kit (Biotium). The supernatant was aspirated, 25 μL of Lysis Buffer was added to each well, and then the plate was allowed to incubate for 15 min with orbital shaking. 25 μL of each sample was transferred to an opaque, white, 96-well plate, then 50 μL of Assay Buffer with 0.2 mg/mL D-luciferin was added, and luminescence was measured immediately using an Infinite MPlex Plate Reader (Tecan), with an integration time of 1000 ms. Data was analyzed after subtracting the luminescence of a blank (Lysis and Assay Buffer with 0.2 mg/mL D-luciferin in a well with no cells), and reported relative to a buffer-treated control.

Assays with THP1-Dual reporter cells were performed based on vendor instructions. Briefly, 5.6×10^5^ cell/mL were added to a 96-well plate in 180 µL RPMI with 10% heat inactivated FBS and P/S cells along with 20 μL of treatment specified in each figure caption, then allowed to incubate for 24 h. To detect IRF3 controlled Lucia luciferase reporter signal, 10 µL of treated cell supernatant was transferred to an opaque, white, 96-well plate, then 50 µL of QUANTI-Luc Reagent (InvivoGen) was added and luminescence was measured immediately using an Infinite MPlex Plate Reader, with an integration time of 100 ms. Data was analyzed after subtracting the luminescence of a blank (QUANTI-Luc Reagent plus fresh media), and reported relative to a buffer treated control. To detect NF-κB controlled secreted embryonic alkaline phosphatase (SEAP) reporter signal, 20 µL of treated cell supernatant was added to 180 µL QUANTI-Blue Reagent (InvivoGen) in a clear 96-well plate. The plate was incubated at 37 °C with orbital shaking until a visible color difference was observed. Absorbance at 640 nm was measured using an Infinite MPlex Plate Reader. Data was analyzed after subtracting the luminescence of a blank (QUANTI-Blue Reagent plus fresh media).

#### Western Blot

HEK293T pGL4.45 reporter cells were added to a 6-well plate at 3×10^5^ cells/mL in 3 mL of DMEM with 10% FBS and P/S, then allowed to incubate for 24 h. Cells were treated with 600 μL of the treatment specified in each figure caption, then allowed to incubate for 6 h. Cells were washed twice with ice-cold 1× PBS and then lysed in RIPA lysis buffer (Thermo Fisher) supplemented with 1× Halt Protease and Phosphatase Inhibitor Cocktail (Thermo Fisher) and 5 mM EDTA (Thermo Fisher) for 10 min at 4 °C with gentle orbital shaking. Cell lysates were centrifuged at 15,000×g, 4 °C for 15 min, the supernatant was recovered, and the protein concentration was quantified by DC (detergent-compatible) protein assay (Bio-Rad). The sample was diluted to reach a final composition of 1× Laemmli Sample Buffer (Bio-Rad) and 25 mM DTT and then heated to 95 °C for 5 min. All lysates were stored at −80 °C until analysis.

Samples were loaded at 25 μg of total protein. Samples were run on a 4–20% Mini-PROTEAN TGX Precast protein gel alongside Precision Plus Protein Dual Color Standards (Bio-Rad) and transferred to a 0.2 μm nitrocellulose membrane (Bio-Rad) using a Trans-Blot SD Semi-Dry Transfer Cell (Bio-Rad). The membrane was blocked with 5% (w/v) Nonfat Dry Milk (Cell Signaling) in tris-buffered saline and 0.1% (v/v) Tween 20 (TBST) for 1 h at RT with gentle orbital shaking. The membrane was incubated with primary antibodies anti-STING (1:1000 dilution, Cell Signaling #13647), anti-TBK1 (1:1000, Cell Signaling #3504), anti-phospho-TBK1 (Ser172) (1:1000, Cell Signaling #5483), anti-IRF3 (1:1000, Cell Signaling #4302), or anti-phospho-IRF3 (Ser396) (1:1000, Cell Signaling #29047) in 5% (w/v) BSA in TBST overnight at 4 °C with gentle orbital shaking. The membrane was incubated with the secondary antibody anti-rabbit IgG, HRP (1:2000, Cell Signaling #7074) in 5% (w/v) Nonfat Dry Milk in TBST for 1 h at RT with gentle orbital shaking. The membrane was incubated with anti-β-actin, HRP (1:25,000, Abcam ab49900) in 5% (w/v) Nonfat Dry Milk in TBST for 1 h at RT with gentle orbital shaking. Western Lightning Ultra (Revvity) chemiluminescent substrate was added to the membrane, and it was imaged using a ChemiDoc MP imaging system (Bio-Rad).

#### Immunocytochemistry

HEK293T pGL4.45 reporter cells were added to a Lab-Tek II Chamber Slide (precoated in 0.01% (w/v) poly-L-lysine solution for 5 min) at 3×10^5^ cells/mL in 400 µL of DMEM with 10% FBS and P/S, then allowed to incubate for 24 h. Cells were treated with 50 μL of the treatment specified in each figure caption, then allowed to incubate for 6 h. Cells were fixed with 4% formaldehyde in 1× PBS for 10 min, permeabilized by 0.1% (v/v) Triton X-100 in 1× PBS 10 min, blocked with 1% (w/v) BSA and 0.05% (v/v) Tween 20 in 1× PBS for 30 min, and stained with a primary rabbit anti-TBK1 antibody (1:66, Abcam #ab235253) at 4 °C overnight. Cells were then stained with secondary Alexa Fluor 488-conjugated donkey anti-rabbit IgG (1:400, Thermo Fisher #A-32790) and simultaneously with Dylight 554 Phalloidin (1:400, Cell Signaling #13054S) at RT for 1 h. ProLong Gold Antifade Mountant with DNA Stain DAPI (Thermo Fisher #P36941) was used to mount a coverslip and was allowed to cure for 24 h at RT. Imaging was performed on a FV4000 confocal laser scanning microscope (Evident) equipped with 405, 488, 561, and 640 nm lasers in the Koch Institute’s Microscopy Core Facility (MIT). Images were acquired with a 100× silicone oil immersion objective. All images were acquired at consistent laser settings. A single Z-slice containing features of interest is displayed. Images were pseudocolored using ImageJ software.

#### Enzyme-Linked Immunosorbent Assays (ELISA)

DuoSet ELISA kits (R&D Systems) for human CXCL10 (#DY266), human IFN-β (#DY814), mouse CXCL10 (#DY466), mouse IFN-β (#DY8234), mouse IL-6 (#DY406), mouse TNF-α (#DY410), and mouse IFN-γ (#DY485) were used with 1-Step TMB ELISA Substrate Solution (Thermo Scientific) following vendor instructions. Cytokine concentrations were calculated using a four-parameter logistic curve-fit of a standard curve, with the bottom parameter constrained at the mean absorbance of the blank (GraphPad PRISM software). The limit of detection (LOD) was calculated by adding 3 standard deviations to the mean absorbance of the blank and converting this to a concentration using the standard curve. Conditions where analyte was below the LOD were labeled as not detected (ND).

#### *In vitro* Cytokine Induction

KURAMOCHI or A2780 cells at 5.6×10^4^ cell/mL were added to a 96-well plate in 180 µL of the media used to culture each cell line (described above), then allowed to incubate for 24 h. Cells were treated with 20 µL of the treatment specified in each figure caption, then allowed to incubate for 24 h. Plates were centrifuged at 500×g, then the supernatant was sampled and stored at −20°C until analysis. Human CXCL10 and IFN-β levels were measured by ELISA, as described above.

#### mRNA Sequencing

KURAMOCHI cells were added to a 6-well plate at 1×10^5^ cells/mL in 3 mL of DMEM/F12 1:1 with 10% FBS and P/S, then allowed to incubate for 24 h. Cells were treated with 600 μL of the treatment specified in each figure caption, then allowed to incubate for 6 h. Cells were washed with 1× PBS and then detached using 100 μL TrypLE Express Enzyme. Cells were diluted with 200 μL DMEM/F12 1:1 with 10% FBS and P/S and then counted with a Cellometer Auto 1000.

The cell suspension was centrifuged at 500×g for 5 min and then the pellet was resuspended in DNA/RNA Shield (Zymo Research Corporation) at 2×10^5^ cells/mL to preserve RNA.

Preserved cells in DNA/RNA Shield were shipped to Plasmidsaurus for mRNA-sequencing. Briefly, total RNA was extracted, converted to complementary DNA via reverse transcription and second-strand synthesis, followed by tagmentation, library indexing, and amplification. Libraries were sequenced using an Ilumina NovaSeq instrument. A 3’ end counting approach was used to capture differential gene expression. Sequencing data was processed using the default Plasmidsaurus pipeline. Quality was assessed using FastQC v0.12.1. Reads were quality filtered using fastp v0.24.0 with poly-X tail trimming, 3’ quality-based tail trimming, a minimum Phred quality score of 15, and a minimum length of 50 bp. Quality-filtered reads were aligned to the reference genome using STAR aligner v2.7.11 with non-canonical splice junction removal and output of unmapped reads, followed by coordinate sorting using samtools v1.22.1. PCR and optical duplicates were removed using UMI-based deduplication with UMIcollapse v1.1.0, resulting in a range of 5.1-15.1 million deduplicated reads per replicate. Gene-level expression quantification was performed using featureCounts (subread package v2.1.1) with strand-specific counting, multi-mapping read fractional assignment, exons and three prime UTR as the feature identifiers, and grouped by gene_id. Differential expression was done with edgeR v4.0.16 after filtering for low-expressed genes with edgeR::filterByExprwith default values. Functional enrichment was performed using gene set enrichment analysis with gseapy v0.12 using the MSigDB Hallmark gene set, normalized enrichment score and familywise-error rate adjusted *P* values are reported.

### Animal Experiments

#### Animal Care

All animal experiments were approved by the MIT Committee on Animal Care (CAC, protocol #2404000660) and were conducted under the oversight of the Division of Comparative Medicine (DCM). Female C57BL/6 mice were purchased from Jackson Laboratory and housed at the Koch Institute for Integrative Cancer Research at MIT animal facility in cages of no more than five animals with controlled temperature (25 °C), 12 h light–dark cycles, and free access to food and water.

#### Tumor Models

For the BPPNM model, 3×10^6^ BPPNM cells (expressing luciferase) were suspended in 200 µL 1× PBS and then injected IP in 8-9 week-old female C57BL/6 mice. For the KPCA.C model, 1×10^6^ KPCA.C cells (expressing mCherry-SIY and luciferase) were suspended in 200 µL 1× PBS and then injected IP in 8-9 week-old female C57BL/6 mice. Tumor burden was monitored using 2D IVIS bioluminescence imaging. Mice were injected IP with 200 µL of 15 mg/kg D-luciferin, sodium salt (GoldBio, #LUCNA), and luminescence was measured 15 min later using the IVIS Spectrum In Vivo Imaging System (Perkin Elmer). Images were analyzed with Living Image software. Prior to each experiment, tumor burden was measured by IVIS bioluminescence and mice were divided into groups with comparable distributions of total tumor bioluminescence signal. Mice with no detectable tumor or poorly dispersed tumor found only at the injection site were excluded from subsequent experiments.

#### Biodistribution

The BPPNM model was initiated as described above, with mice moved onto AIN-93M Maintenance Purified Diet (TestDiet #1810541) at least one week prior to biodistribution measurements. Mice were administered a 20 μg dose of STING peptide-polyanion conjugate either without a carrier or loaded in the C3 LNP formulation IP in 200 µL of 1× PBS 15 days after tumor inoculation. 200 µL of 1× PBS was administered on the same schedule as a control. Mice were injected IP with 200 µL of 15 mg/kg D-luciferin, sodium salt, luminescence was measured 15 min later using the IVIS Spectrum In Vivo Imaging System, and then mice were euthanized at 4 hours after initial treatment. Immediately following, necropsies were performed to harvest upper genital tract (UGT), omentum, liver, kidneys, spleen, intestines, heart and lungs from each mouse. Organs were immersed in RPMI 1640 media in 24-well plates and placed on ice until imaging. Organs were imaged using bioluminescence to track BPPNM tumors. STING peptide-polyanion conjugate was imaged using the fluorescence of its sulfo-cyanine5-label using a 640 excitation and 680 emission filter. Living Image Software was used to measure the bioluminescent total flux in [p/s], bioluminescent average radiance in [p/s/cm²/sr], fluorescent total radiant efficiency in [p/s] / [µW/cm²], fluorescence average radiant efficiency in [p/s/cm²/sr] / [µW/cm²], and area in [cm²] of each organ. To correct for background fluorescence, average fluorescence of the organ in the vehicle 1× PBS treated mice was subtracted from each measurement. The fraction of recovered fluorescence was computed as the total radiant efficiency of the organ divided by sum of the total radiant efficiency for all organs recovered from an individual mouse.

#### Acute Toxicity

The BPPNM model was initiated as described above. Mice were administered a 20 μg dose of STING or Scr peptide-polyanion conjugate loaded in the C3 LNP formulation IP in 200 µL of 1× PBS 14 days after tumor inoculation. 200 µL of 1× PBS was administered on the same schedule as a control. 24 h after treatment, blood was collected via submandibular bleeding into EDTA Microtubes (Sarstedt) for complete blood counts or Serum Gel CAT Microtubes (Sarstedt) for serum chemistry. Analysis was performed by the Division of Comparative Medicine Comparative Pathology Laboratory (MIT), with complete blood counts analyzed using a HemaVet 950FS (Drew Scientific) and serum chemistry analyzed using a custom IDEXX panel.

#### Pharmacokinetics and Pharmacodynamics

A 20 μg dose of STING peptide-polyanion conjugate loaded in the C3 LNP formulation was administered IP in 200 µL of 1× PBS to 11 week old female C57BL/6 mice. Blood was collected from 3 randomly selected mice out of 5 treated mice at 0 h (immediately), 1 h, 3 h, 6 h, 10 h, 24 h, and 50 h after treatment. At each time point, approximately 50 μL blood was collected via submandibular bleeding into Serum Gel CAT Microtubes (Sarstedt). Serum Gel CAT Microtubes were centrifuged at 10,000×g for 5 min, serum was collected from the topmost fraction and stored at −20 °C until analysis.

STING peptide-polyanion conjugate was quantified in serum by measuring fluorescence of its sulfo-cyanine5 label. 5 μL of serum was diluted with 20 μL of 2% (v/v) Triton X-100 in water, followed by 10 min incubation at 37 °C with orbital shaking. Cyanine5 fluorescence intensity was read in black 384-well flat-bottom plates on an Infinite MPlex Plate Reader (Tecan), with excitation at 640 nm and emission at 680 nm. STING peptide-polyanion conjugate concentrations were calculated using a four-parameter logistic curve-fit of a standard curve, prepared by diluting LNP stock in serum from an untreated mouse (GraphPad PRISM software). The limit of detection (LOD) was calculated by adding 3 standard deviations to the mean absorbance of the blank and converting this to a concentration using the standard curve. Conditions where analyte was below the LOD were labeled as not detected (ND). Mouse CXCL10 levels in serum were measured by ELISA, as described above.

#### *In Vivo* Cytokine Profiling

The BPPNM model was initiated as described above. Mice were administered a 20 μg dose of STING or Scr peptide-polyanion conjugate loaded in the C3 LNP formulation IP in 200 µL of 1× PBS 14 days after tumor inoculation. 6 h after treatment, blood was collected via submandibular bleeding into Serum Gel CAT Microtubes (Sarstedt). Serum Gel CAT Microtubes were centrifuged at 10,000×g for 5 min, serum was collected from the topmost fraction, and supplemented with 1× Halt Protease and Phosphatase Inhibitor Cocktail (Thermo Fisher). Immediately after bleeding, mice were euthanized and ascites was collected via peritoneal lavage with 1 mL 1× PBS. Ascites were centrifuged at 4,000×g for 15 min at 4 °C, the supernatant was collected and supplemented with 1× Halt Protease and Phosphatase Inhibitor Cocktail. Tumor nodules located on the omentum were collected and washed with ice-cold 1× PBS. Approximately 60 mg of tumor (range 53-63 mg) was added to 1 mL RIPA lysis buffer (Thermo Fisher) and supplemented with 1× Halt Protease and Phosphatase Inhibitor Cocktail. Tumor was diced with scissors, homogenized with a syringe plunger, and incubated at 4 °C for 15 min with occasional brief sonication. Tissue homogenate was centrifuged at 15,000×g for 15 min at 4 °C, and the supernatant was collected. All samples were stored at −80 °C until analysis. Mouse CXCL10, IFN-β, IL-6, TNF-α, and IFN-γ levels were measured by ELISA, as described above. Cytokine levels in ascites and tumor were normalized to total protein levels in each sample, measured by DC protein assay (Bio-Rad) following vendor instructions.

#### Therapeutic Efficacy Studies

The BPPNM model was initiated as described above. In a monotherapy study, mice were administered a 20 μg dose of STING or Scr peptide-polyanion conjugate loaded in the C3 LNP formulation IP in 200 µL of 1× PBS on days 10, 13, and 16 after tumor inoculation. 200 µL of 1× PBS was administered on the same schedule as a control. In a combination therapy study, mice were administered a 20 μg dose of STING or Scr peptide-polyanion conjugate loaded in the C3 LNP formulation IP in 200 µL of 1× PBS on days 10, 13, 16, 19, and 22 after tumor inoculation. 200 µL of 1× PBS was administered on the same schedule as a control. In groups specified in figure captions, mice were also administered a 100 μg dose of anti-mouse PD-1 antibody (Bio X Cell #BE0273 InVivoMAb, Clone 29F.1A12) in 200 µL of 1× PBS on days 11 and 17 after tumor inoculation. Weight and tumor bioluminescence were measured throughout the study. Mice were monitored and euthanized when body condition score dropped below 2, weight loss exceeded 20%, or poor responsiveness was observed.

The KPCA.C model was initiated as described above. Mice were administered a 20 μg dose of STING or Scr peptide-polyanion conjugate loaded in the C3 LNP formulation IP in 200 µL of 1× PBS on days 7, 10, 13, and 16 after tumor inoculation. 200 µL of 1× PBS was administered on the same schedule as a control. Weight and tumor bioluminescence were measured throughout the study. Mice were monitored and euthanized when body condition score dropped below 2, weight loss exceeded 20%, or poor responsiveness was observed.

#### Immunophenotyping

The BPPNM model was initiated as described above. Mice were administered a 20 μg dose of STING or Scr peptide-polyanion conjugate loaded in the C3 LNP formulation IP in 200 µL of 1× PBS on days 10, 13, 16, and 19 after tumor inoculation. 200 µL of 1× PBS was administered on the same schedule as a control. On day 20 after tumor inoculation (12 h after the final dose), mice were euthanized to collect organs.

Tumor, localized on the omentum, was collected and stored in RPMI media on ice. To the gentleMACS™ C Tubes (Miltenyi #130-096-334), 2.5 mL digestion mixture (1 mg/mL collagenase type V, 0.25 mg/mL hyaluronidase, 0.1 mg/mL DNAse I in RPMI media) and tumor were added. Tumor was minced using forceps and kept on ice. Tumors were digested using a gentleMACS Octo Dissociator (Miltenyi), m_imp-tumor-02 program. Tubes were incubated under shaking (150 rpm) at 37 °C for 30 min. Tubes were returned to ice and digested using m_impTumor_03 program. Digested tumors were diluted 1:1 with FACS buffer (1% BSA and 2 mM EDTA) to final concentration of 1 mM EDTA and 0.5% BSA as EDTA quenches the enzymatic reaction. Dissociated tumors were filtered through a 70 μm filter (Miltenyi #130-110-916) into a new 15 mL tube to obtain a single cell suspension and the filter was rinsed with 2 mL FACS buffer to complete the transfer. Cells were collected by spinning 15 mL tubes at 300×g for 5 min at 4 °C. Supernatant was decanted and 2 mL ACK lysis buffer was added, then the tube was gently vortexed to resuspend cells and incubated for 2 min to lyse red blood cells. To quench the ACK, 5 mL FACS buffer was added and the tube was spun at 300×g for 5 min. This process was repeated until a white cell pellet was obtained. Supernatant was removed and cells resuspended in 200 μL of FACS buffer, counted, and left on ice until flow staining.

Ascites was collected using a 3 mL syringe attached with an 18G needle (Air-Tite #ML3181). Cold RPMI (2.5 mL) was injected into the peritoneal cavity, the peritoneal cavity was gently massaged and then the fluid was collected in the syringe and transferred to 15 mL tube and kept on ice. Tubes were centrifuged at 300×g for 5 min at 4 °C, supernatant decanted, and cells resuspended in 1 mL of enzyme digestion mixture by gentle vortex. For the digestion, cells were then incubated at 37 °C for 30 min on an orbital shaker incubator under agitation (150 rpm). Following enzymatic digestion, 1 mL of FACS buffer was added to quench the enzymatic reaction. Cells were centrifuged at 300×g for 5 min at 4 °C. Supernatant was decanted and 2 mL ACK lysis buffer was added, then the tube was gently vortexed to resuspend cells and incubated for 2 min to lyse red blood cells. To quench the ACK, 5 mL FACS buffer was added and the tube spun at 300×g for 5 min. Cells were resuspended in 200 μL FACS buffer by vigorous pipetting and passed through a 70 μm filter into a fresh tube to obtain a single cell suspension. Cells were counted and left on ice until flow staining.

Based on an average cell count, 10^6^ cells of each tumor sample and all of the ascites cells (typically 5×10^5^-10^6^ cells) were transferred to a 96-well V bottom plate for staining. The plate was centrifuged (500×g for 3 min at 4 °C for this and subsequent steps), dumped, and cells resuspended in 50 μL Zombie Live/Dead stain (1:100) diluted in PBS and incubated for 10 min at room temperature. Cells were washed with 150 μL FACS buffer then Fc receptors were blocked with anti-CD16/32 (Biolegend #101339) at 1 μg per well in 50 μL for 15 min at 4 °C. Cells were washed with 150 μL FACS buffer then stained with antibodies in 50 μL, diluted in FACS buffer and Brilliant Stain Buffer (BD #BDB563794) at specified dilutions in **Table S2,** and incubated for

30 min at 4 °C. Cells were washed with 150 μL FACS buffer then fixed in 200 μL of 2% formaldehyde in 1× PBS for 20 min at 4 °C. Cells were centrifuged and resuspended in FACS buffer and stored at 4 °C until analyzed. Compensation was calculated with UltraComp eBeads™ Plus Compensation Beads and single stained cells where appropriate. Flow cytometry was performed using a BD Symphony A3 equipped with HTS (BD Biosciences, New Jersey), and data analyzed in FlowJo V10.

### Data Analysis and Statistics

All statistical tests were performed using GraphPad PRISM 10. Comparisons between two groups were performed via unpaired *t*-tests. Comparisons between multiple groups were performed with one-way ANOVA followed by Tukey’s post-hoc test or two-way ANOVA followed by Šídák’s post-hoc test. All tests were two-tailed. For survival studies, comparisons were performed using the log(rank) (Mantel–Cox) test. For mRNA sequencing data analysis, results were considered significant at *P* ≤ 0.05. When an assay determined that an analyte was below the limit of detection (LOD), the result was highlighted as not detected (ND). If any replicates were ND, summary statistics were not computed and statistical comparisons were not performed. Data is presented as mean ± standard deviation (SD) if presented on a linear scale or geometric mean ± geometric SD if presented on a logarithmic scale.

## AUTHOR CONTRIBUTION STATEMENT

J.A.K. conceived of the project, designed experiments, performed experiments, analyzed data, and wrote the first draft of the paper. J.A.K., J.T., V.F.G. and A.K. performed animal experiments. J.T. assisted in conjugate synthesis, LNP optimization and characterization, and carried out *in vitro* experiments. A.G. assisted in LNP optimization and characterization, produced LNPs, and carried out *in vitro* experiments. V.F.G. led the design, execution, and analysis of flow cytometry studies. Z.K.T. performed CD and HPLC experiments to characterize peptide-conjugates. M.M.B. assisted in the design of the LNP optimization screen. P.T.H and J.A.K. acquired funding to support this project. P.T.H. provided project oversight and critical review of this manuscript. All authors discussed results and contributed to manuscript review and editing.

## ACKNOWLEDGEMENTS

We thank the Koch Institute’s Robert A. Swanson (1969) Biotechnology Center for technical support, specifically the Peterson (1957) Nanotechnology Materials, Preclinical Modeling, Imaging & Testing, Flow Cytometry, Microscopy, Hope Babette Tang (1983) Histology, and Biopolymers & Proteomics Core Facilities. We also acknowledge technical support from the MIT Department of Biology Biophysical Instrumentation Facility and Department of Chemistry Instrumentation Facility. We also acknowledge support from the MIT Division of Comparative Medicine and the Comparative Pathology Laboratory. We thank DongSoo Yun for assistance with TEM imaging, Magalie Boucher for discussion of acute toxicity data, and Namita Nabar for assistance with LNP generation. We also thank Grace Wolczanski and Stefani Spranger for providing luciferized BPPNM and KPCA.C cells. Finally, we are grateful to the entire Hammond lab for advice and support, particularly Ivan Pires and Apoorv Shanker for helpful discussion of this work. We thank Eva Cai for critical review of this manuscript.

## DECLARATION OF COMPETING INTREST

J.A.K. and P.T.H. are inventors on a patent filed by the Massachusetts Institute of Technology relating to STING peptide-polyanion conjugate therapeutics. P.T.H. is a member of the Board of Alector Therapeutics and the Board of Sail Biomedicine, a Flagship company, and a former member of the Scientific Advisory Board of Moderna Therapeutics and the Board of LayerBio. All other authors report no competing interests.

