## Supplementary Material for "A multivalent peptide-polymer conjugate material mimics STING to therapeutically activate innate immune signaling"

**This file includes:**

Figures S1 – S32
Tables S1 – S2

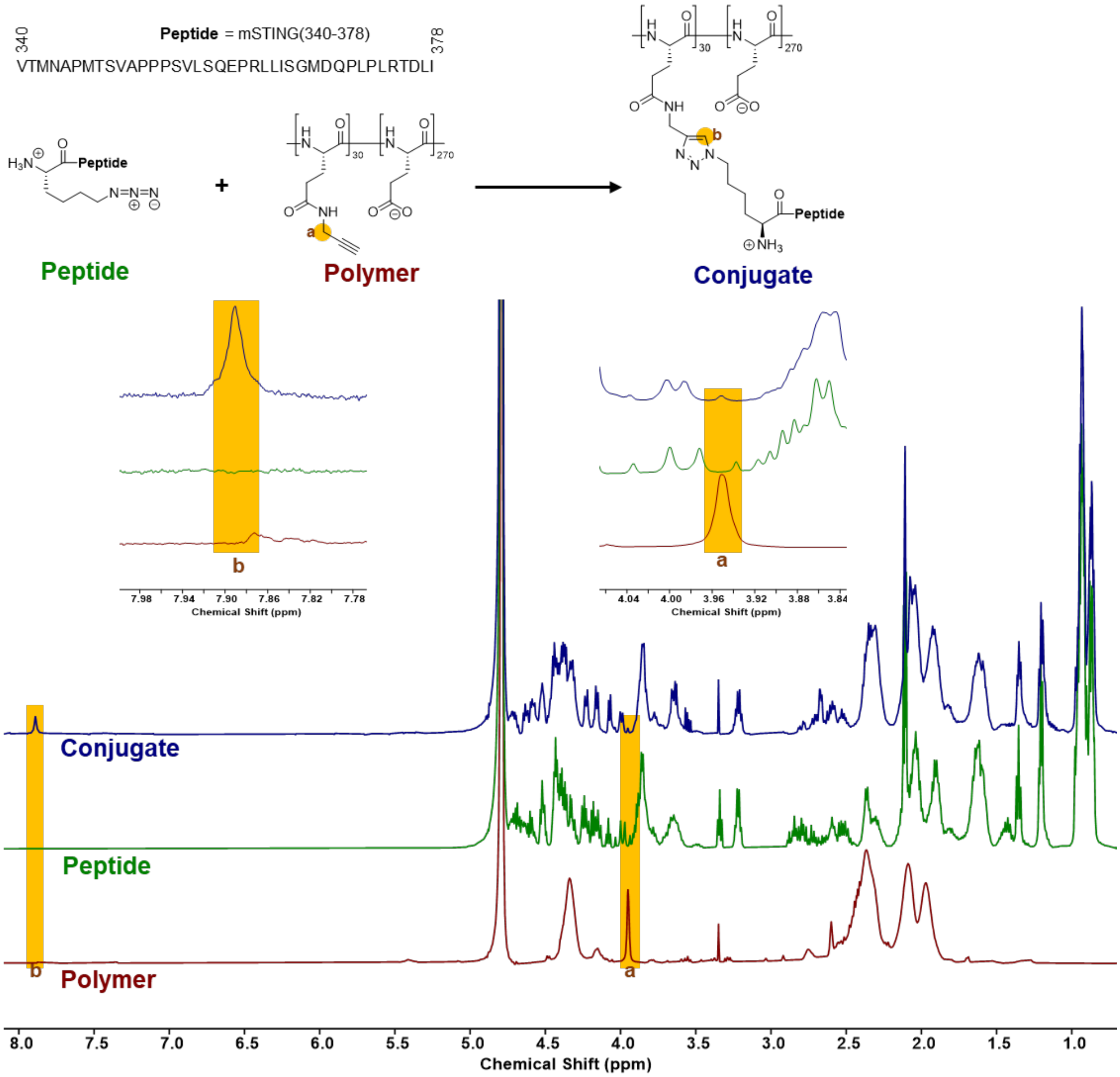


**Figure S1: $^1$H NMR of multivalent peptide-polyanion conjugate synthesis reaction.** 500 MHz NMR in $D_2O$. Reactants poly(L-glutamate)-graft-alkyne (sulfo-Cyanine5-labeled) ("Polymer" in red) and azido-lysine-mSTING(340-378) peptide ("Peptide" in green) are compared to purified product poly(L-glutamate)-graft-mSTING(340-378) peptide conjugate. Key changes that indicate quantitative conversion of polymer alkynes are highlighted in yellow including the complete disappearance of the alkyne-adjacent methylene peak ("a") as well as the appearance of a triazole proton peak ("b") in the conjugate.

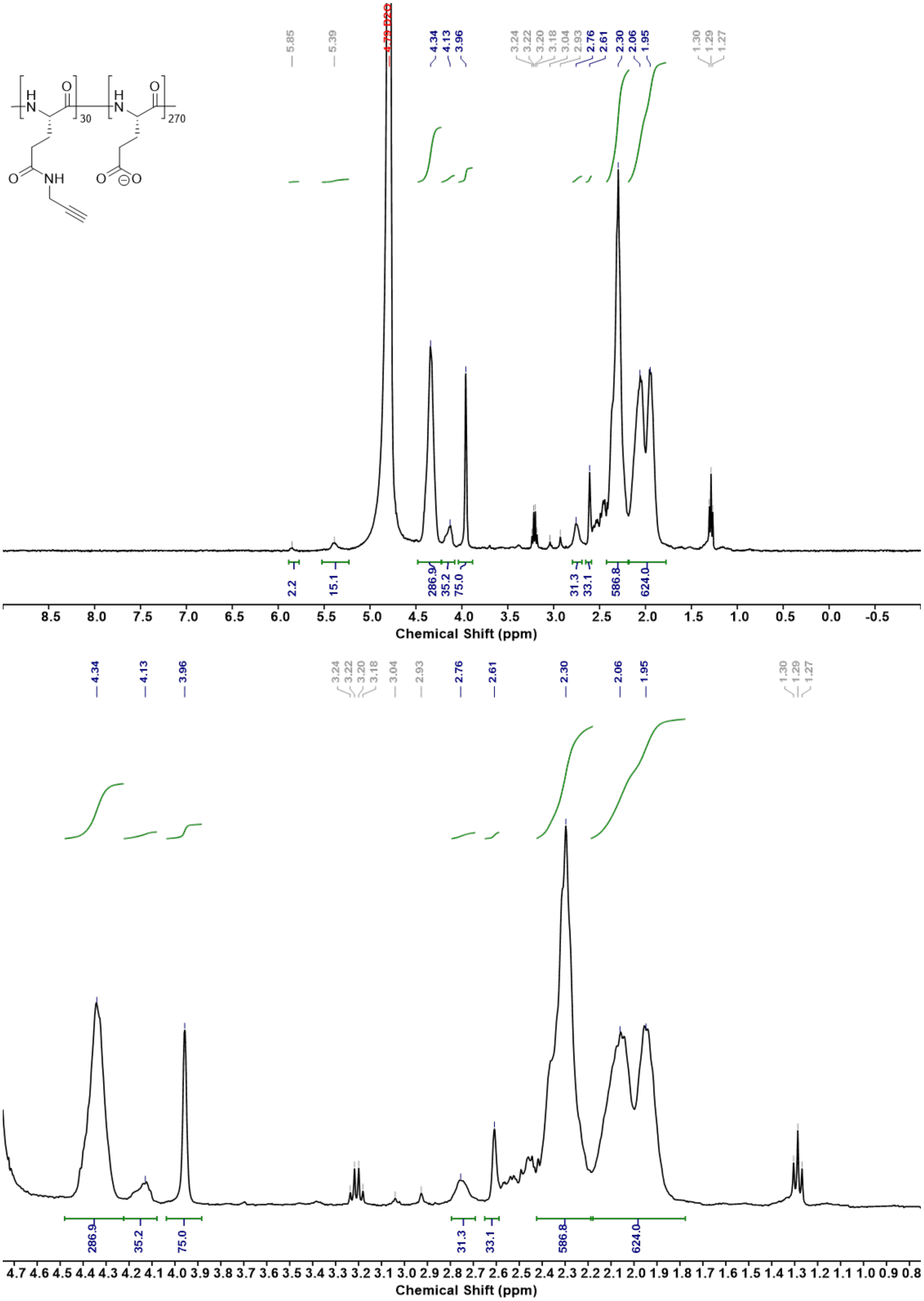


**Figure S2: [1]H NMR of poly(L-glutamate)-graft-alkyne.** 1 mg/mL solution in $D_2O$ on 400 MHz NMR.

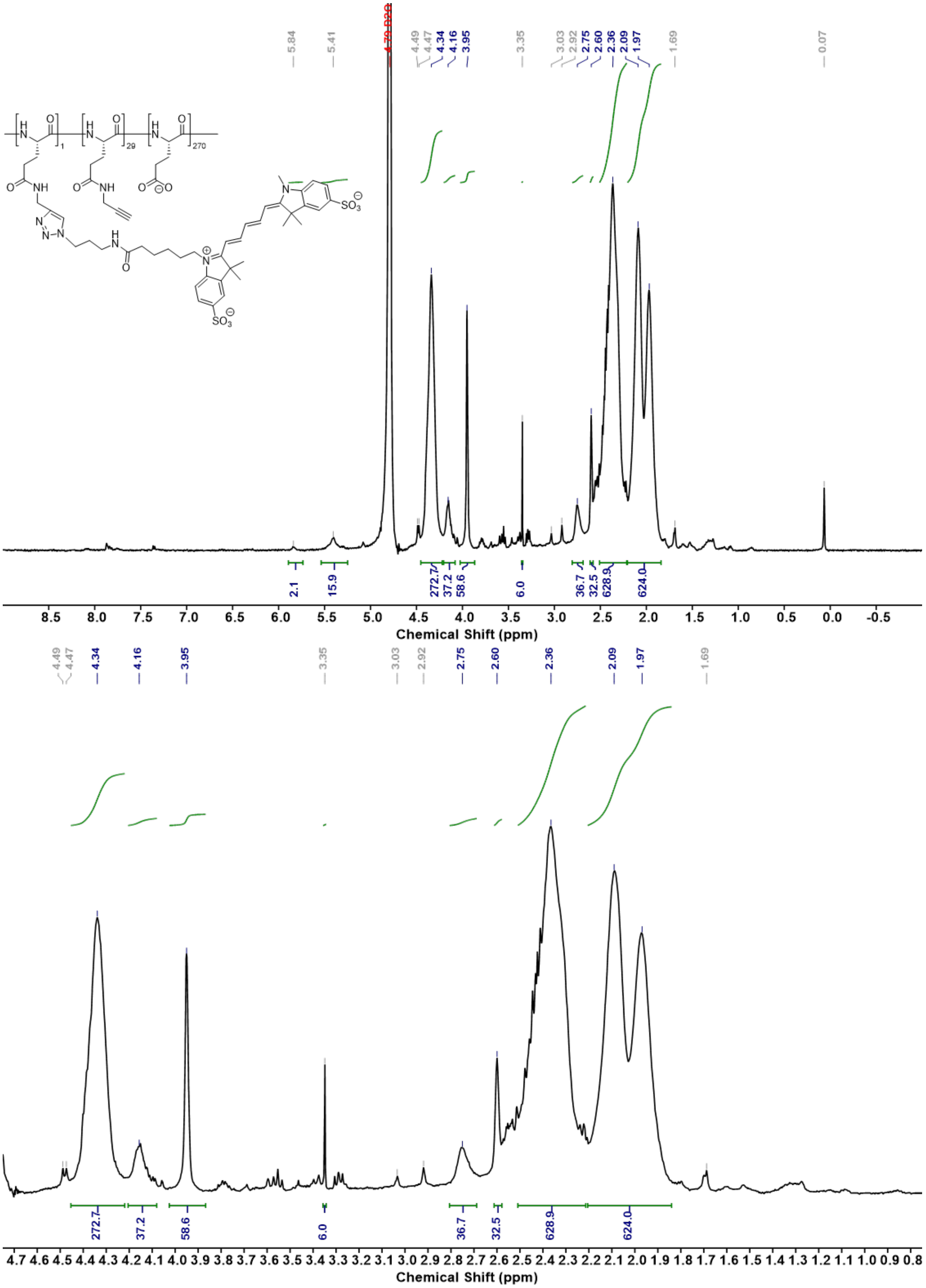


**Figure S3: [1]H NMR of poly(L-glutamate)-graft-alkyne (sulfo-Cyanine5-labeled).** 1 mg/mL solution in $D_2O$ on 500 MHz NMR.

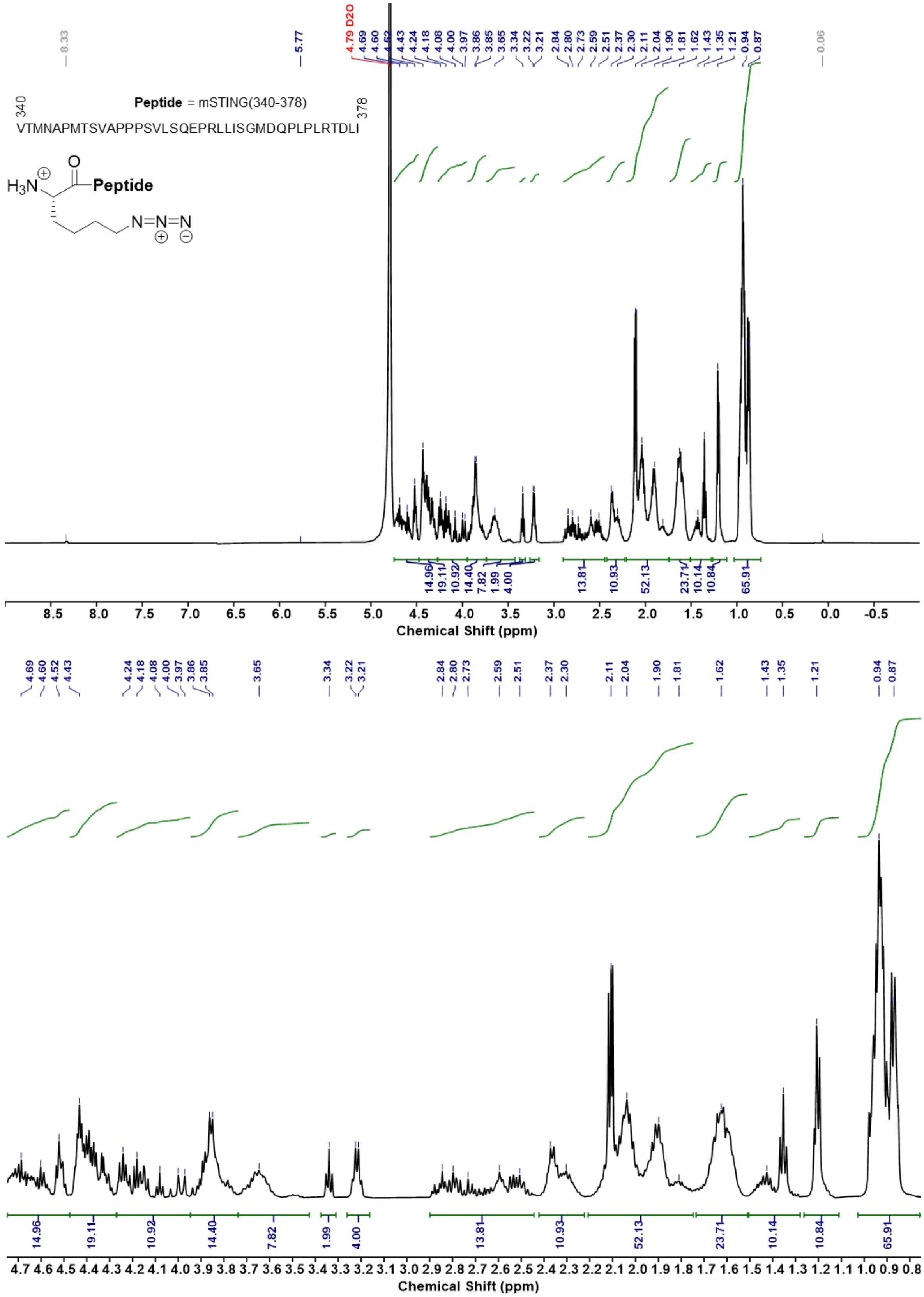


**Figure S4: [1]H NMR of azido-lysine-mSTING(340-378) peptide.** 1 mg/mL solution in $D_2O$ on 500 MHz NMR.

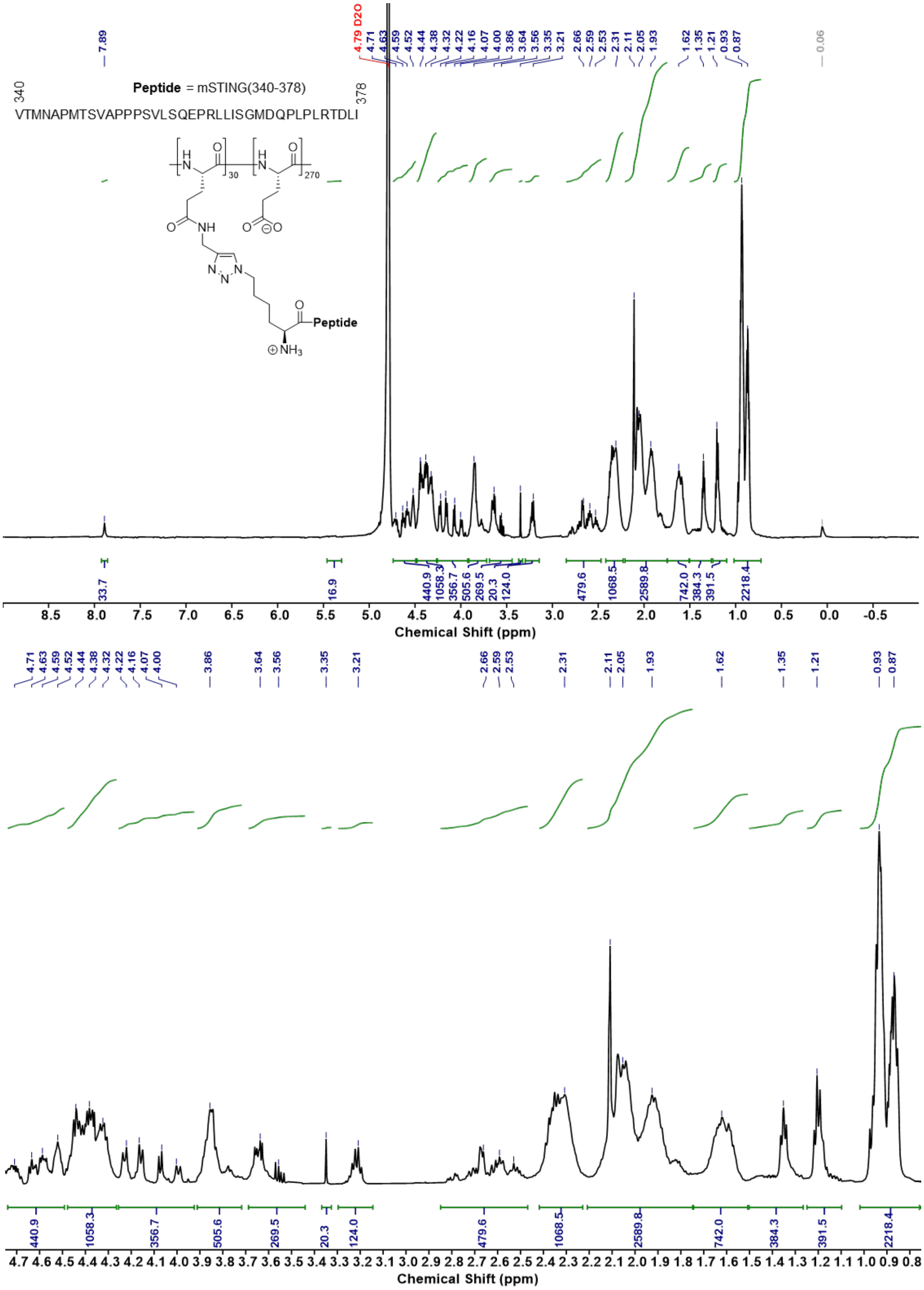


**Figure S5: [1]H NMR of poly(L-glutamate)-graft-mSTING(340-378) peptide conjugate (sulfo-Cyanine5-labeled).** 1 mg/mL solution in $D_2O$ on 500 MHz NMR.

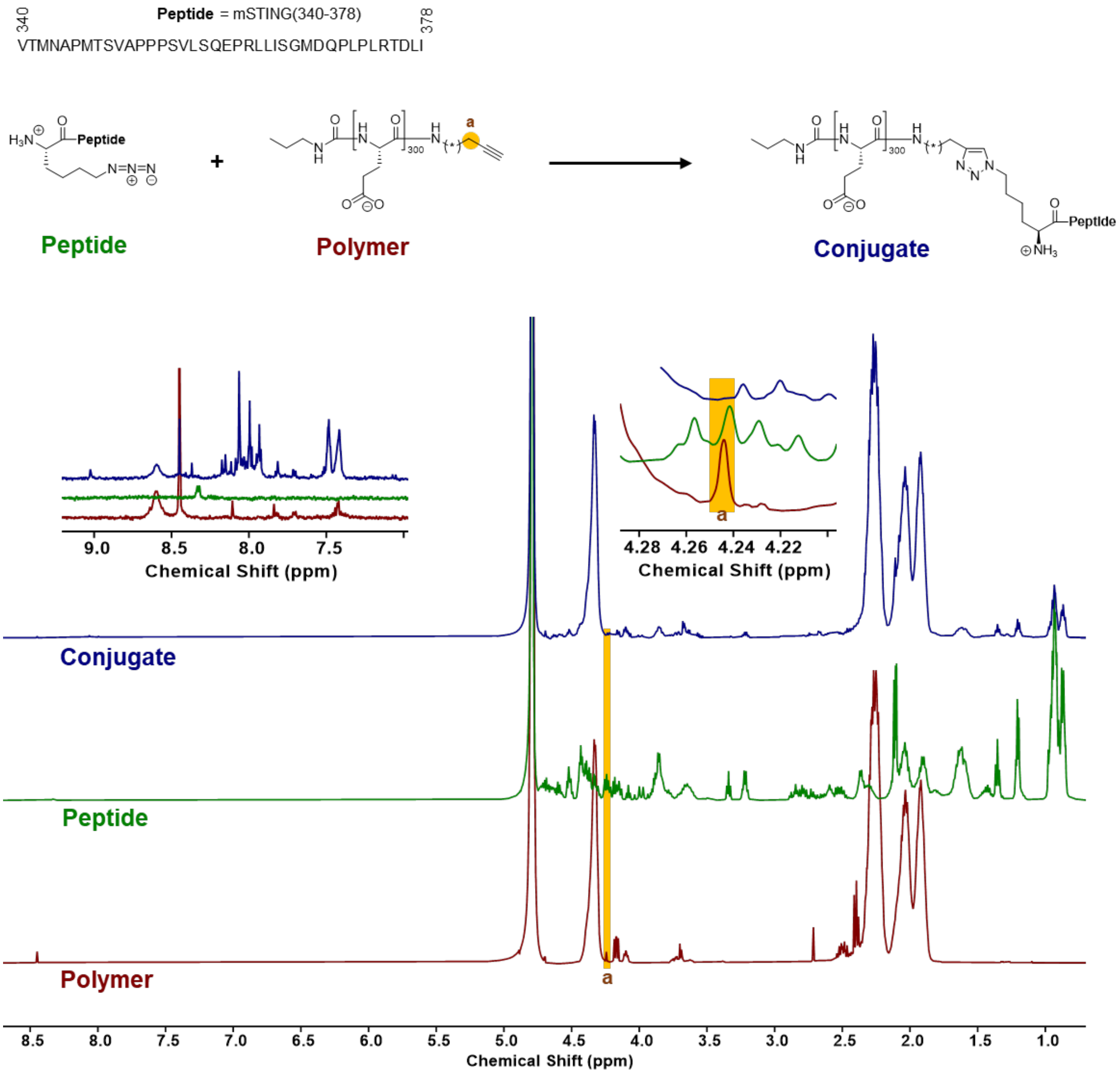


**Figure S6: [1]H NMR of monovalent peptide-polyanion conjugate synthesis reaction.** 500 MHz NMR in $D_2O$. Reactants end-functionalized alkyne-poly(L-glutamate) ("Polymer" in red) and azido-lysine-mSTING(340-378) peptide ("Peptide" in green) are compared to purified product mSTING(340-378)-poly(L-glutamate) peptide conjugate. Key changes that indicate quantitative conversion of polymer alkynes are highlighted in yellow including the complete disappearance of alkyne-adjacent methylene peak ("a") in the conjugate.

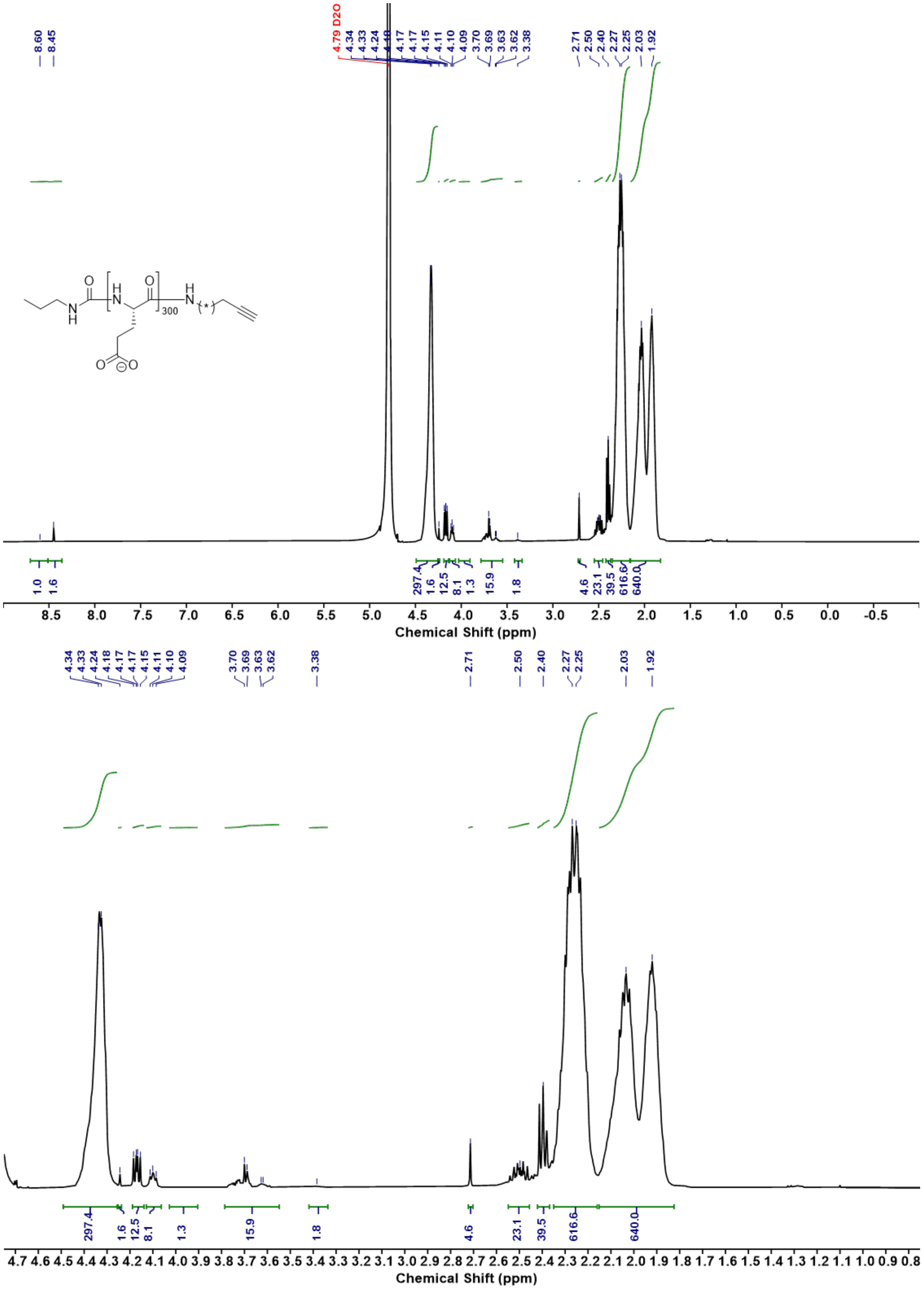


**Figure S7: [1]H NMR of end-functionalized alkyne-poly(L-glutamate).** 1.5 mg/mL solution in $D_2O$ on 500 MHz NMR.

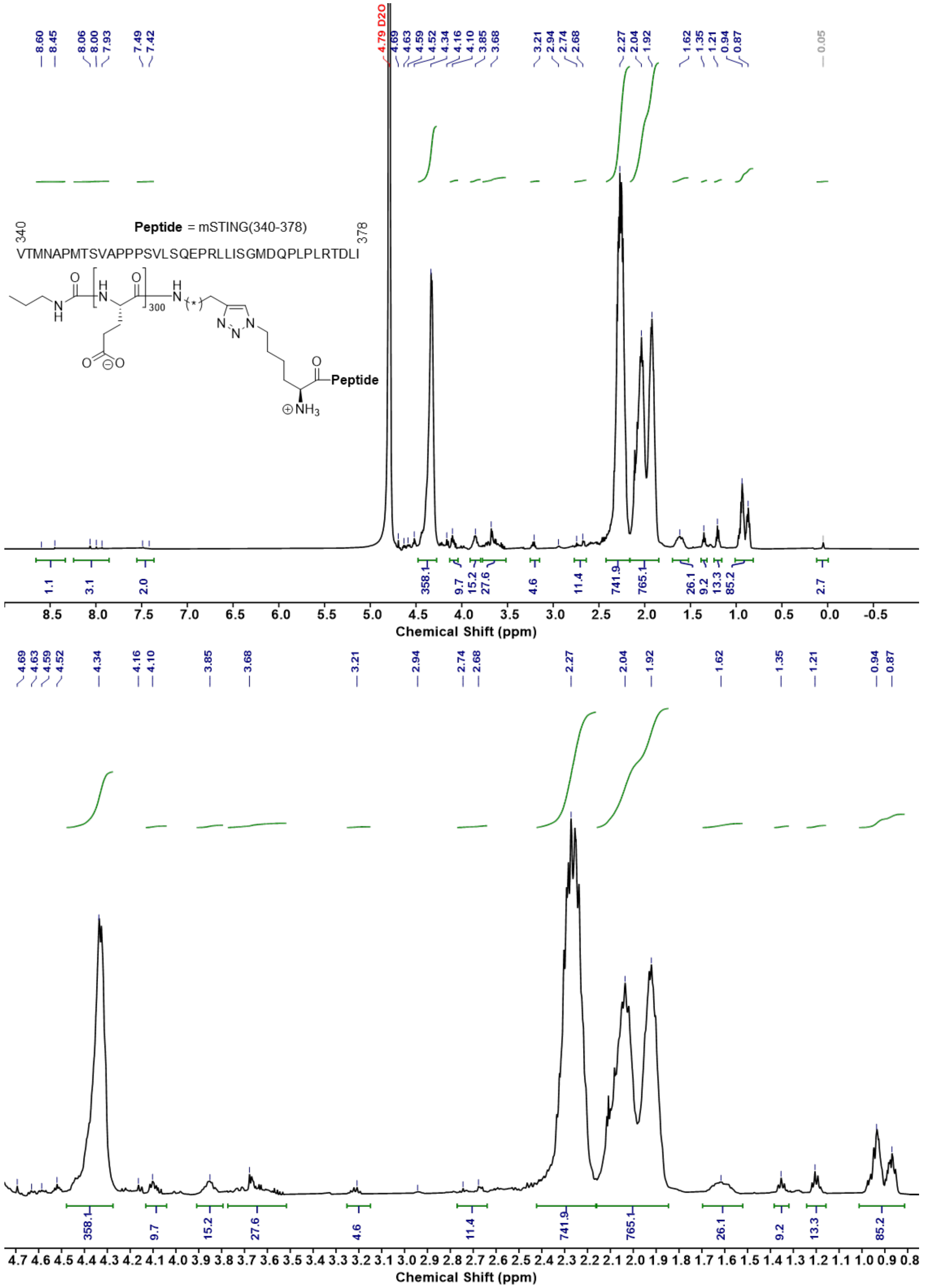


**Figure S8: $^{1}$H NMR of end-functionalized (monovalent) mSTING(340-378)-poly(L-glutamate peptide conjugate.** 1.5 mg/mL solution in $D_2O$ on 500 MHz NMR.

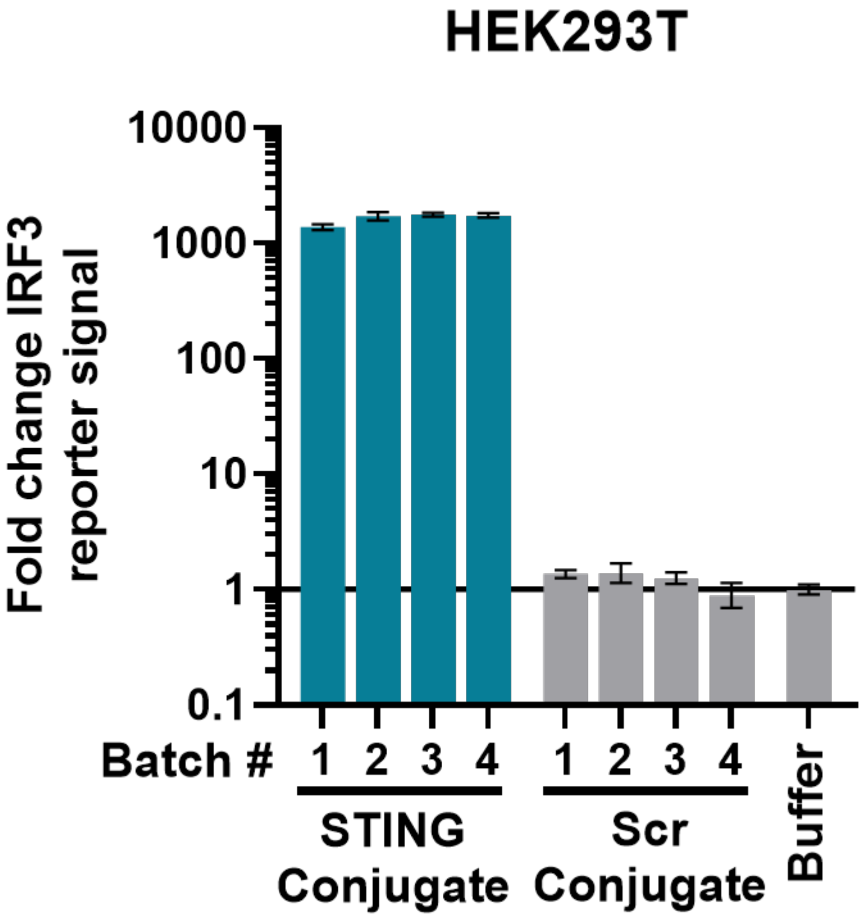


**Figure S9: Peptide-polyanion conjugate has consistent bioactivity across batches.** IRF3 reporter signal relative to buffer treatment for HEK293T reporter cells treated with 8.3 μg/mL STING or Scr conjugate delivered using TransIT-X2, measured 24 h post treatment. N = 4 independently synthesized batches of conjugate, displaying geometric mean ± SD of N = 3 technical replicate activity measurements for each batch.

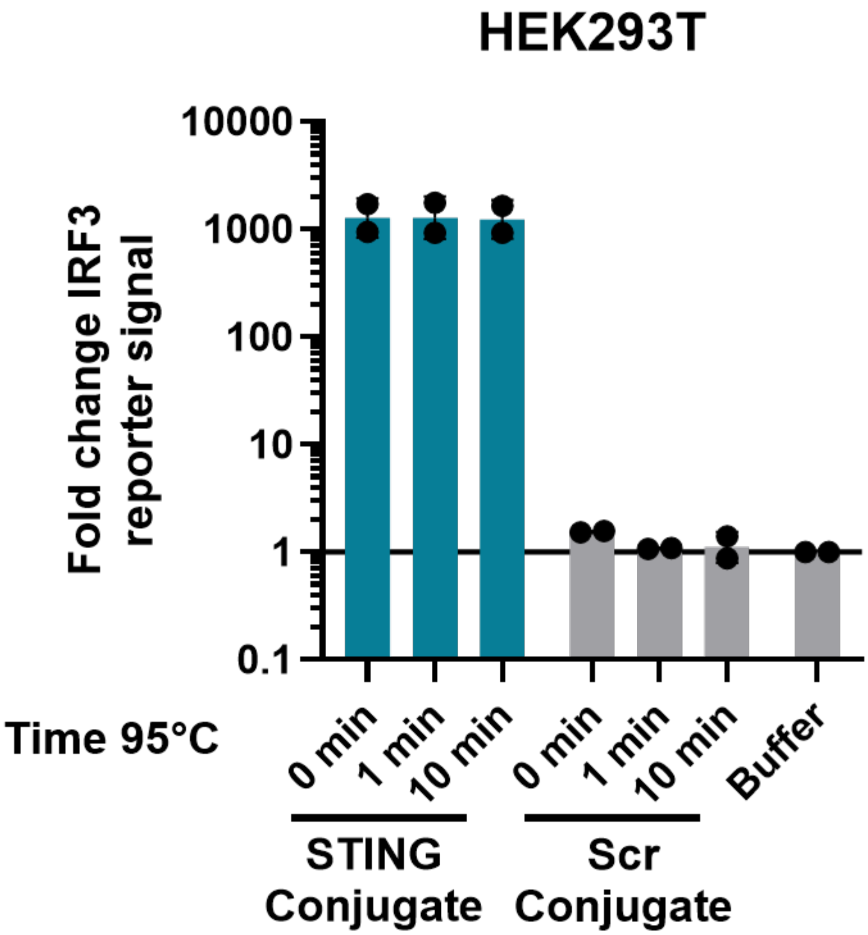


**Figure S10: Peptide-polyanion conjugate activity is stable to thermal denaturation.** STING or Scr peptide-polyanion conjugate was held at 95°C for specified time, allowed to cool to room temperature, then complexed with TransIT-X2 to assay activity. IRF3 reporter signal relative to buffer treatment for HEK293T reporter cells treated with 8.3 µg/mL STING or Scr conjugate delivered using TransIT-X2, measured 24 h post treatment (N = 2 biological replicates). Data represented as geometric mean ± SD.

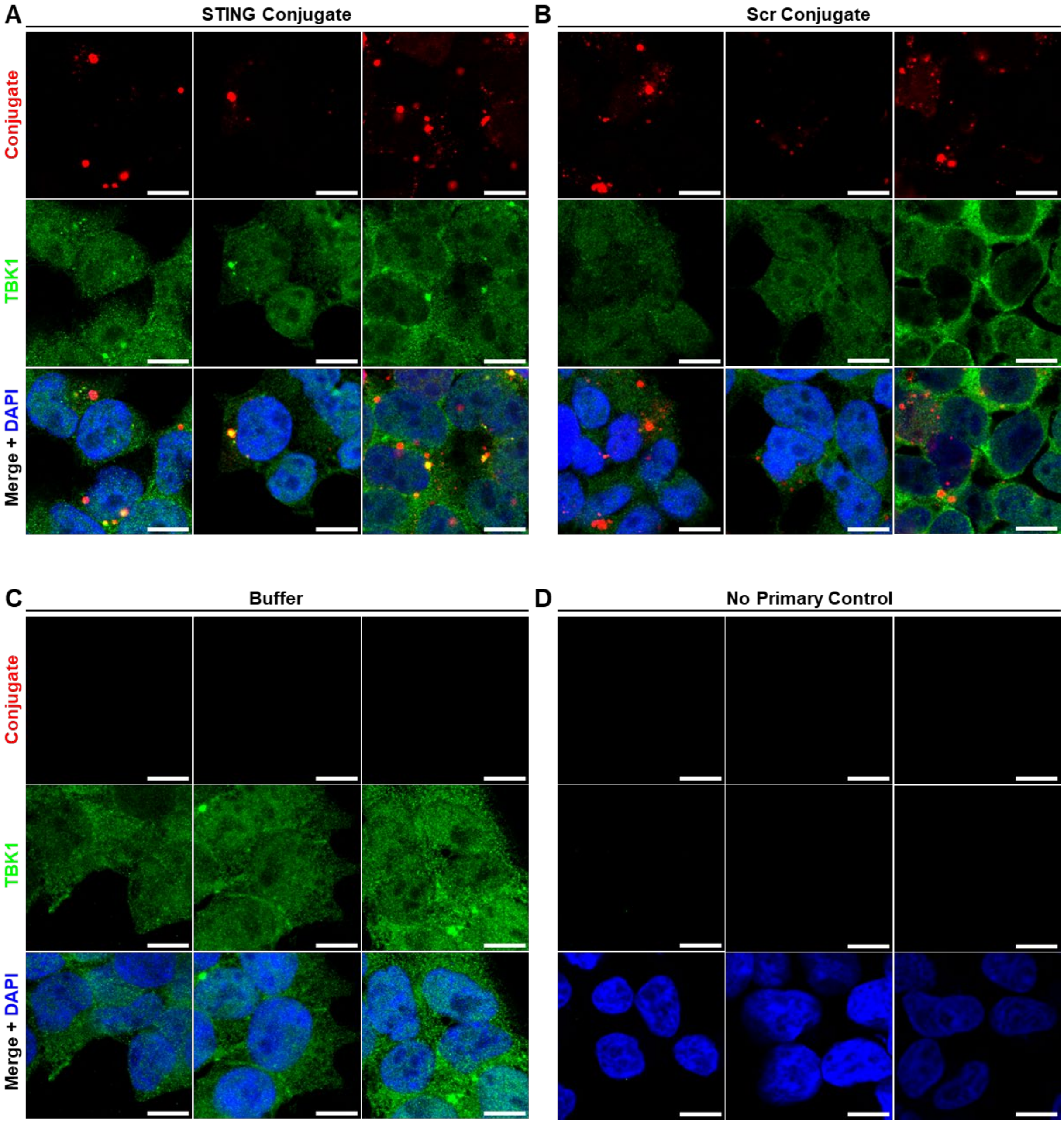


**Figure S11: Biological replicates for confocal imaging experiments.** HEK293T reporter cells were treated with 6.0 µg/mL (**A**) STING or (**B**) Scr peptide-polyanion conjugate using TransIT-X2 as a vehicle, or a (**C**) buffer control, and cells were imaged by confocal microscopy at 6 h post treatment to examine conjugate colocalization with target TBK1. (**D**) A buffer treated control with no primary anti-TBK1 antibody but the same secondary antibody is also included. A representative image from each of N = 3 biological replicates is displayed for each experiment. Scale bar is 10 µm.

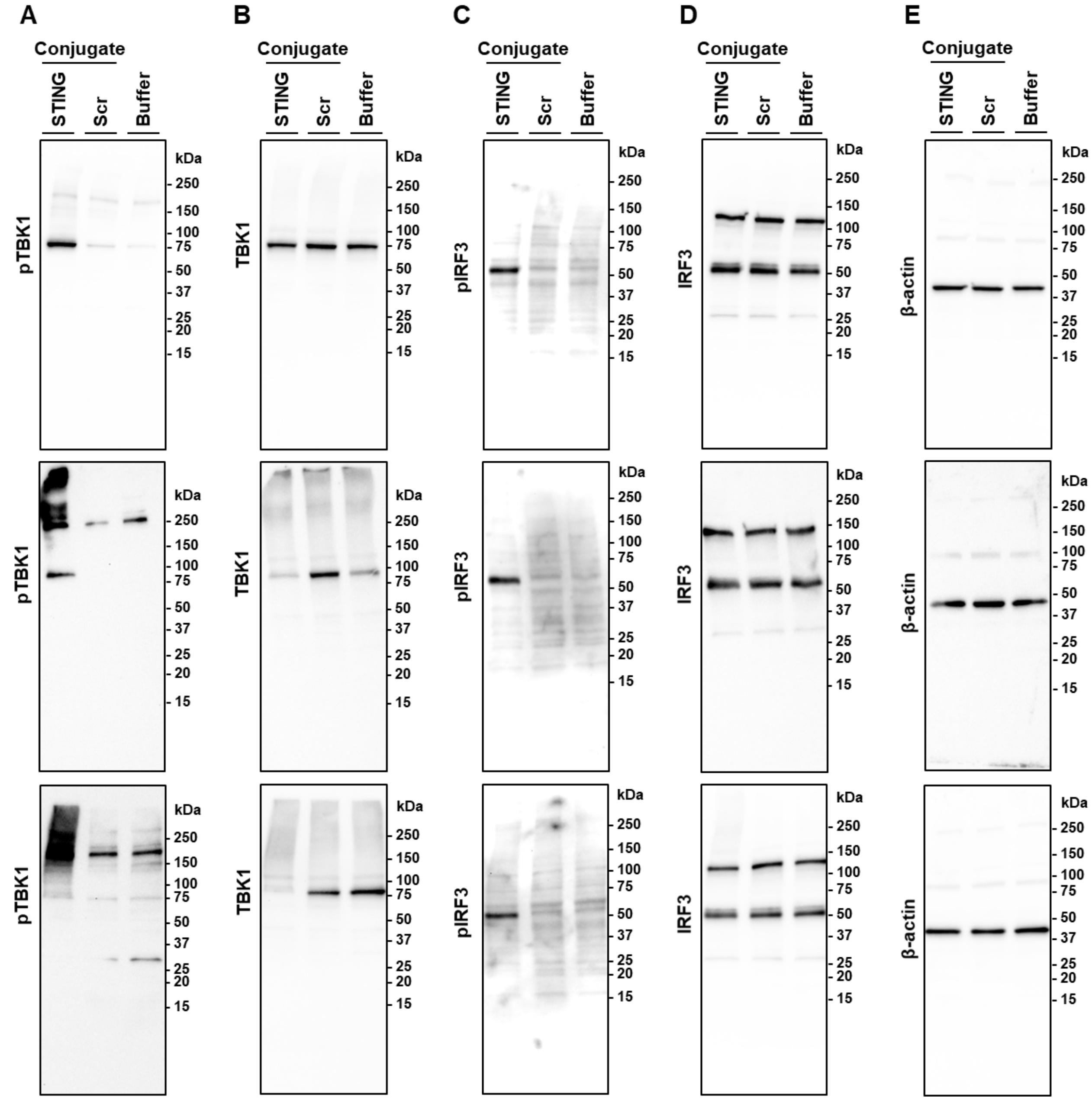


**Figure S12: Full gels of biological replicates for Western blot activity experiment.** HEK293T reporter cells were treated with 8.3 µg/mL STING or Scr peptide-polyanion conjugate using TransIT-X2 as a vehicle, Western blot was performed at 6 h post treatment to examine STING signaling activation. Full gel images showing staining for (**A**) pTBK1 (Ser172), (**B**) TBK1, (**C**) pIRF3 (Ser396), (**D**) IRF3, and (**E**) β-actin. N = 3 biological replicate experiments are shown. We suspect that the higher molecular weight smear observed when staining for pTBK1 in some replicates is a result of binding interactions between pTBK1 and the STING peptide-polyanion conjugate slowing migration through the gel.

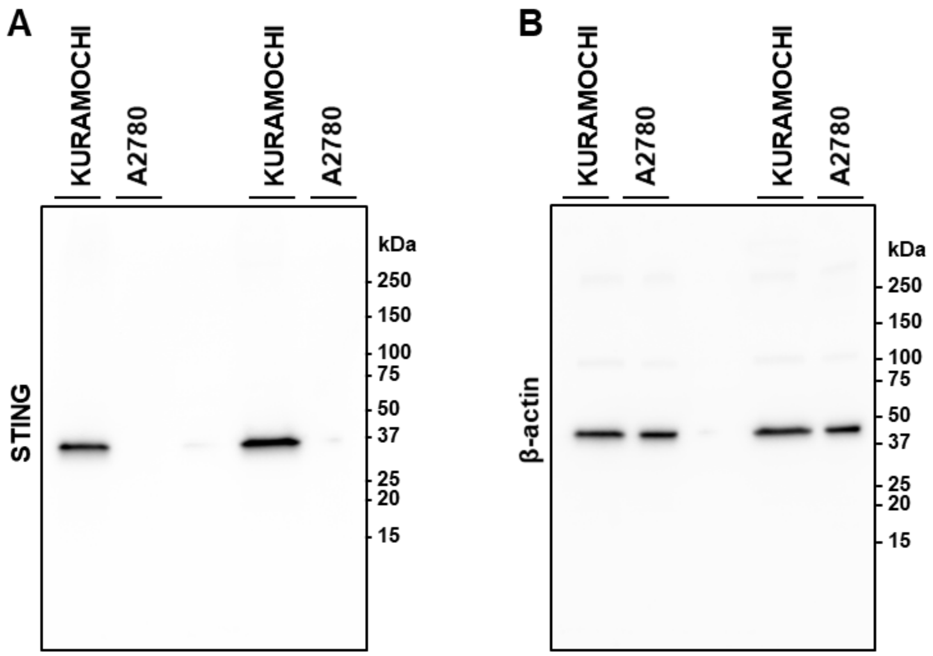


**Figure S13: Full gels of biological replicates for Western blot of ovarian cancer cell lines.** A2780 and KURAMOCHI cell lines were examined. Full gel images showing staining for (**A**) STING and (**B**) β-actin. N = 2 biological replicate experiments are shown.

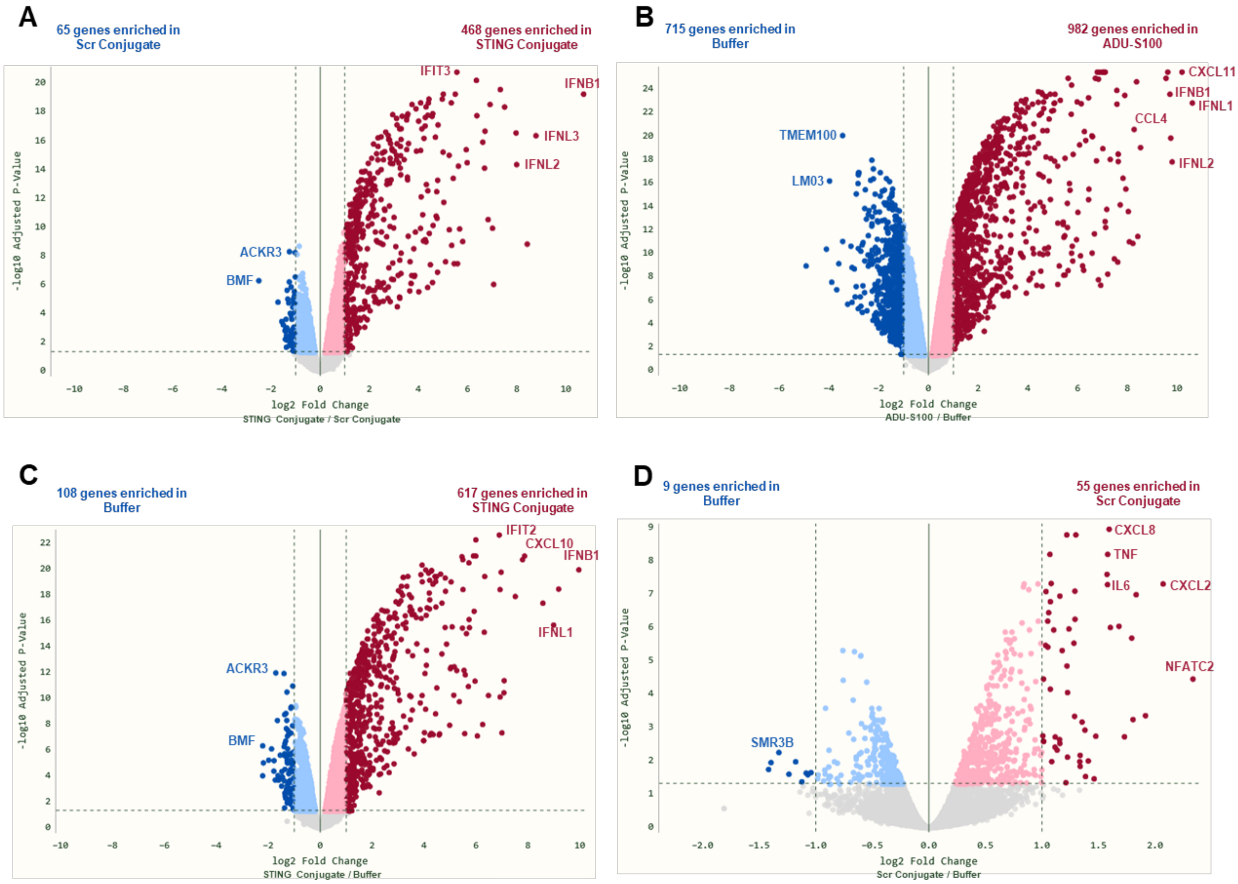


**Figure S14: Volcano plots from mRNA sequencing experiment.** KURAMOCHI cells were treated with 5.0 µg/mL STING or Scr conjugate using TransIT-X2 as a vehicle or 50 µM ADU-S100, mRNA sequencing was performed at 6 h post treatment (N = 4 biological replicates). Volcano plots were generated for (**A**) STING Conjugate compared to Scr Conjugate, (**B**) ADU-S100 compared to Buffer, (**C**) STING Conjugate compared to Buffer, and (**D**) Scr Conjugate compared to Buffer. Counts of differentially expressed genes for each comparison are displayed, using a threshold of $|\log_2 FC| \geq 1$ and Adjusted *P* value ≤ .05. Selected differentially expressed genes are labeled.

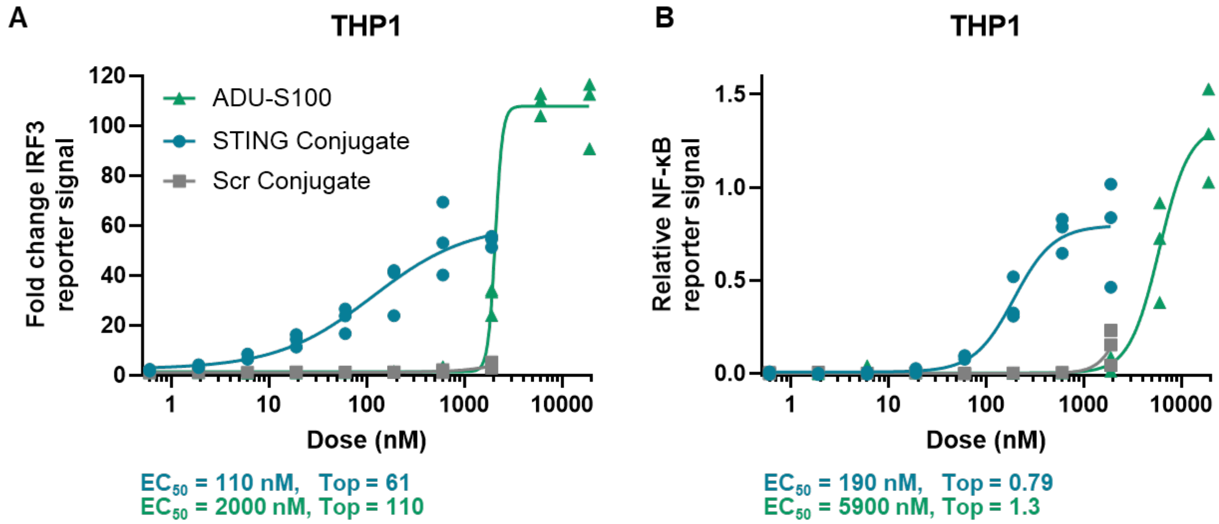


**Figure S15: STING peptide-polyanion conjugate activates IRF3 and NF-κB transcription factors in THP1 monocytes.** (**A**) IRF3 reporter signal relative to buffer treatment and (**B**) relative NF-κB reporter signal for THP1 dual reporter cells treated with STING or Scr conjugate delivered at specified dose of peptide using TransIT-X2, or STING agonist ADU-S100, measured 24 h post treatment (N = 3 biological replicates). A variable slope Hill equation was fit to data, with the $EC_{50}$ and Top reported underneath each plot for the active treatments STING conjugate and ADU-S100.

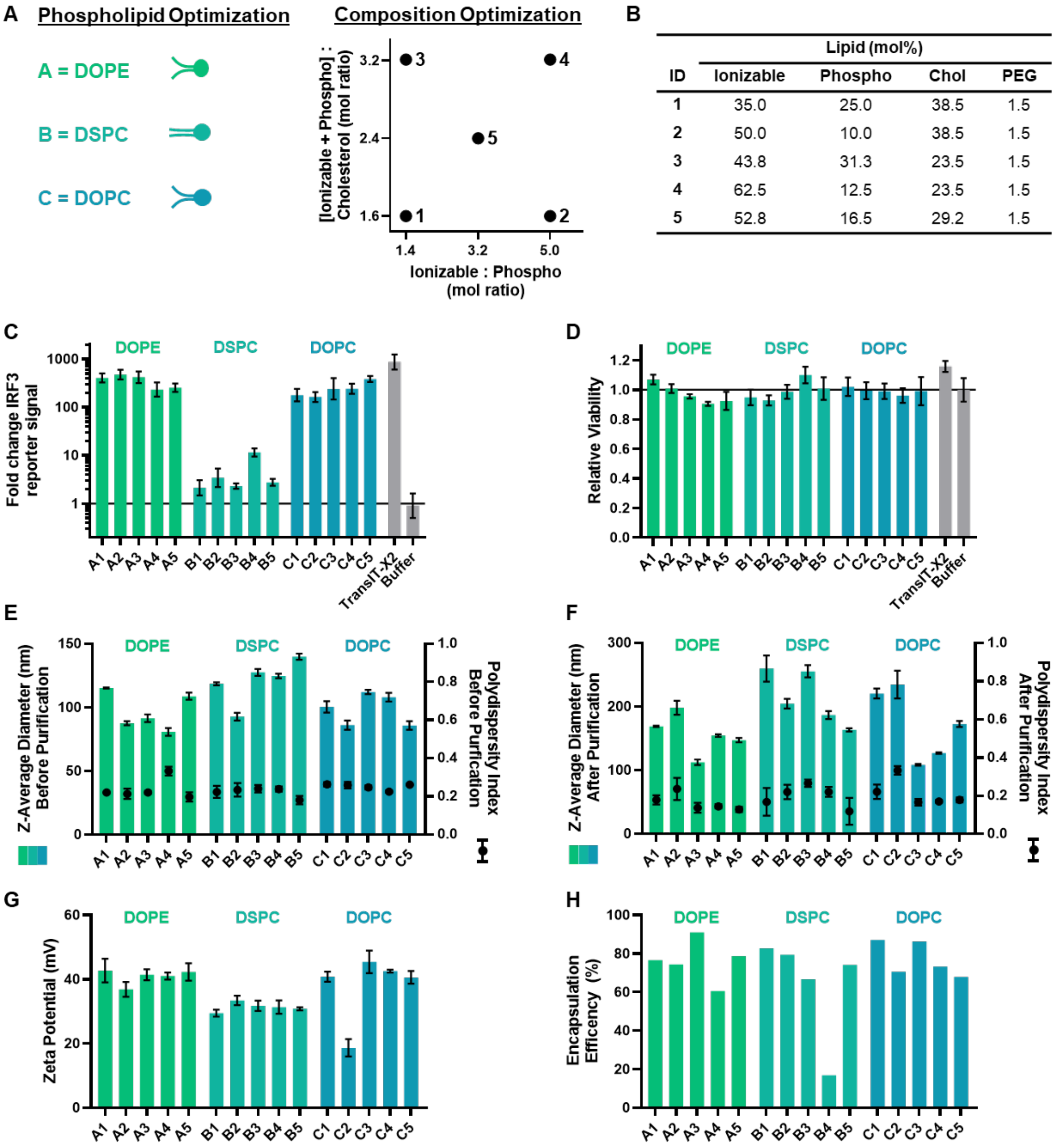


| | Lipid (mol%) | | | |
|---|---|---|---|---|
| ID | Ionizable | Phospho | Chol | PEG |
| 1 | 35.0 | 25.0 | 38.5 | 1.5 |
| 2 | 50.0 | 10.0 | 38.5 | 1.5 |
| 3 | 43.8 | 31.3 | 23.5 | 1.5 |
| 4 | 62.5 | 12.5 | 23.5 | 1.5 |
| 5 | 52.8 | 16.5 | 29.2 | 1.5 |

**Figure S16: Optimization of LNP formulation for multivalent peptide-polyanion conjugate delivery.** (**A**) The identity of the phospholipid component and ratios of ionizable lipid, phospholipid, and cholesterol were varied to optimize LNP stability, encapsulation, and size. The ionizable lipid ALC-0315 was used for all formulations, the mole percentage of PEG lipid was held at 1.5%, and the mass ratio of total lipid to peptide-polyanion conjugate was held at 20:1. (**B**) Mole percentage of each lipid in five compositions examined. (**C**) IRF3 reporter signal and (**D**) viability (resazurin assay) relative to buffer treatment for HEK293T reporter cells treated with 3.3 µg/mL conjugate delivery by each LNP formulation or TransIT-X2 control, 24 h post treatment (N = 3 technical replicates). LNP Z-Average diameter and polydispersity index measured by DLS (**E**) immediately after particle formation (**F**) after purification by centrifugal filtration (N = 3 technical replicates). (**G**) LNP zeta potential measured in water (N = 3 technical replicates). (**H**) Encapsulation efficiency of peptide-polyanion conjugate in LNP measured by native PAGE assay. Data represented as mean ± SD for data on linear scale and geometric mean ± SD for data on log-scale.

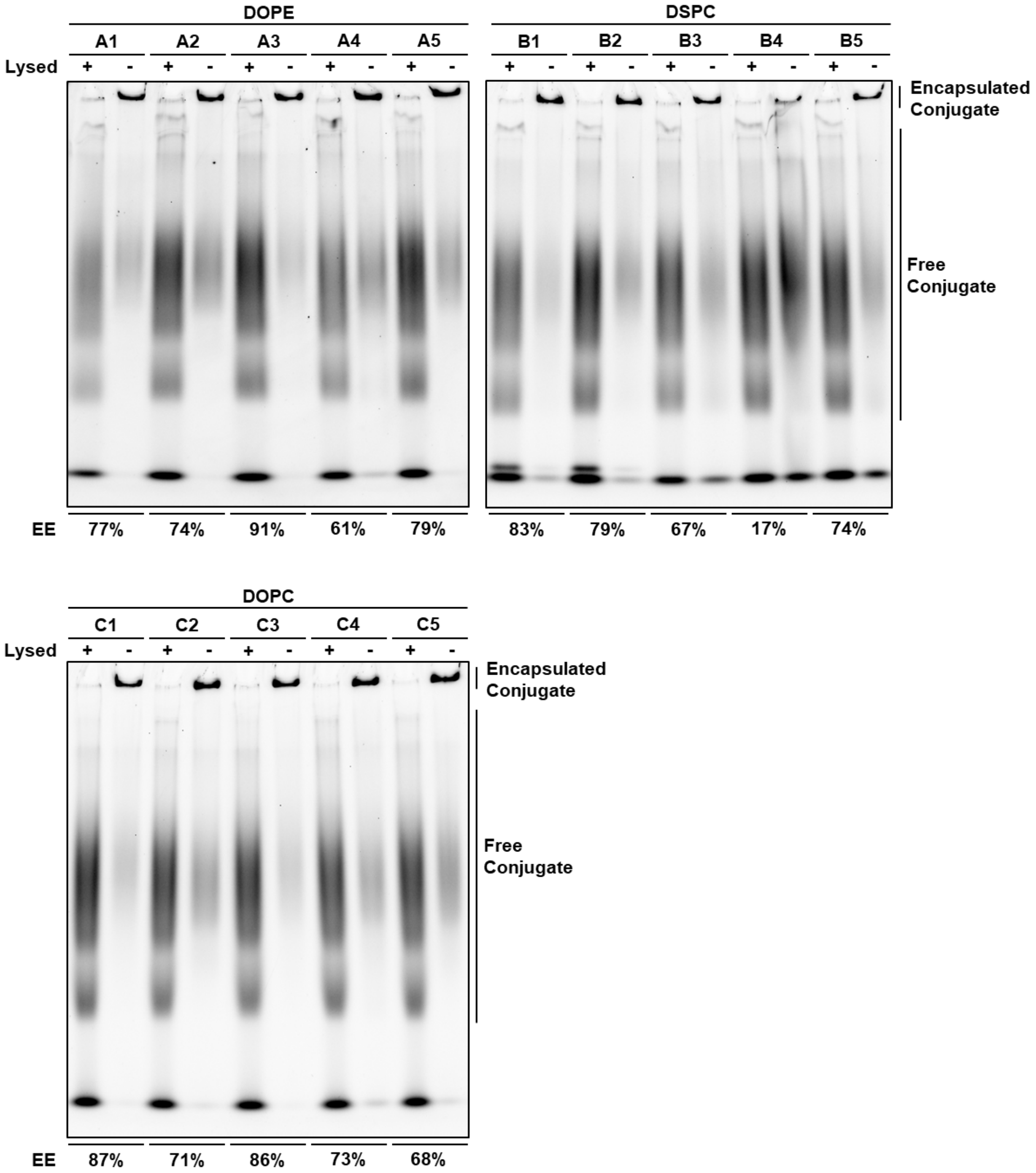


**Figure S17: Native PAGE encapsulation assay results for LNP optimization screen.** LNPs from the screen were loaded into Native PAGE gel intact or after lysis with Triton X-100. Gels were imaged using the Cy5 channel to detect Cy5 tag on peptide-polyanion conjugate. Free conjugate is able to migrate in the gel up to the labeled point, while encapsulated conjugate is trapped in the well. Encapsulation efficiency (EE) was calculated by comparing the total Cy5 signal of free conjugate in lysed particles to intact particles, and is listed underneath the image.

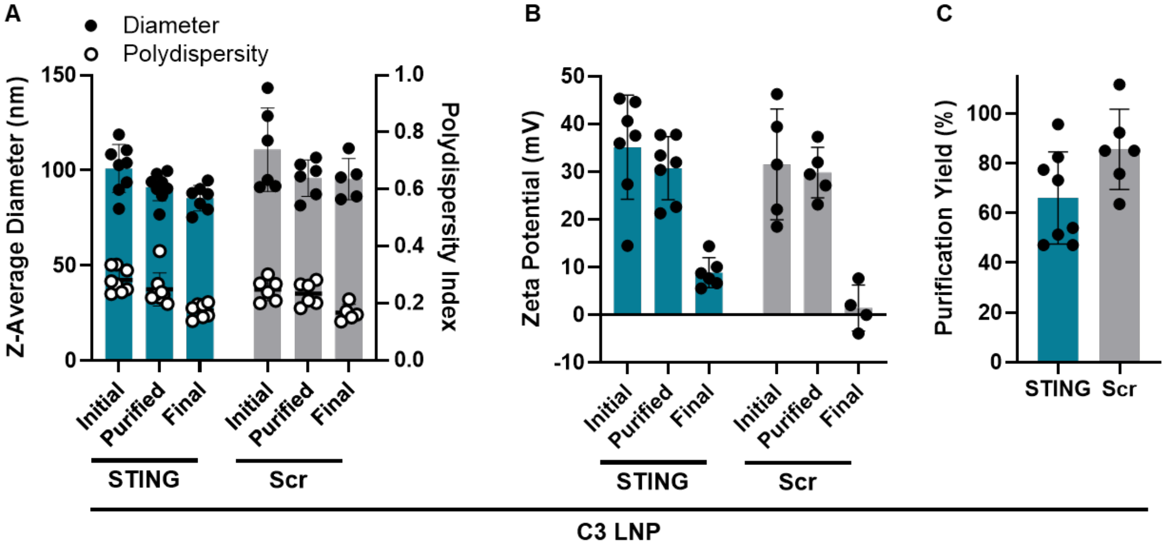


**Figure S18: Characterization of optimized C3 LNP formation process.** (**A**) Z-average diameter, polydispersity index, and (**B**) zeta potential measurements (N = 4-8 independent LNP batches) were taken during the process of generating LNPs loaded with STING or Scr peptide-polyanion conjugate. The "Initial" measurement was taken immediately upon LNP formation, the "Purified" measurement was taken after purification and concentration of LNPs by centrifugal filtration into water, and the "Final" measurement was taken after dilution to a final conjugate concentration of 100 µg/mL in the 1x PBS buffer used for dosing. (**C**) Conjugate yield after purification and concentration of LNPs by centrifugal filtration (N = 6-8 independent LNP batches). Data represented as mean ± SD.

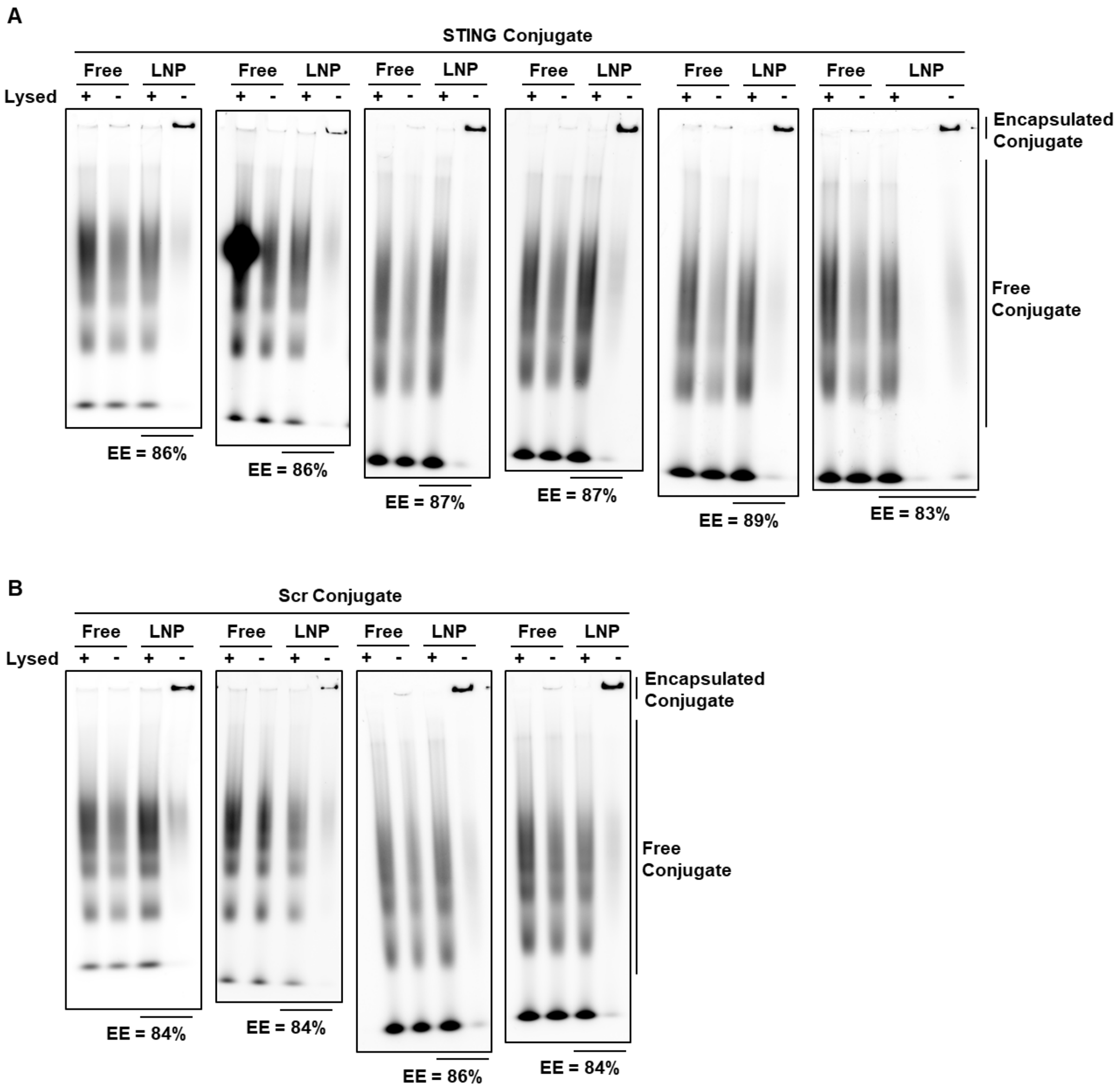


**Figure S19: Native PAGE encapsulation assay results for optimized C3 LNP formulation.** Final purified C3 LNP formulation containing (**A**) STING peptide conjugate (N = 6 independent LNP batches) or (**B**) Scr peptide conjugate (N = 4 independent LNP batches) were loaded into Native PAGE gel intact or after lysis with Triton X-100. Concentration matched free-conjugate that was unencapsulated in a LNP was loaded as a control. Gels were imaged using the Cy5 channel to detect Cy5 tag on peptide-polyanion conjugate. Free conjugate is able to migrate in the gel up to the labeled point, while encapsulated conjugate is trapped in the well. Encapsulation efficiency (EE) was calculated by comparing the total Cy5 signal of free conjugate in lysed particles to intact particles, and is listed underneath the image.

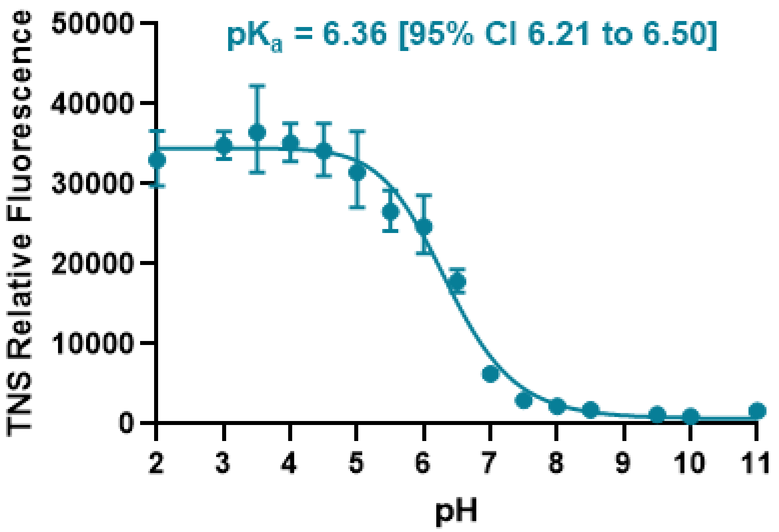


**Figure S20: TNS titration curve of C3 LNPs loaded with STING peptide-polyanion conjugate.** Apparent $pK_a$ was determined by conducting a four-parameter logistic regression in GraphPad Prism and determining pH at 50% normalized TNS fluorescence. (N = 3 independent LNP batches).

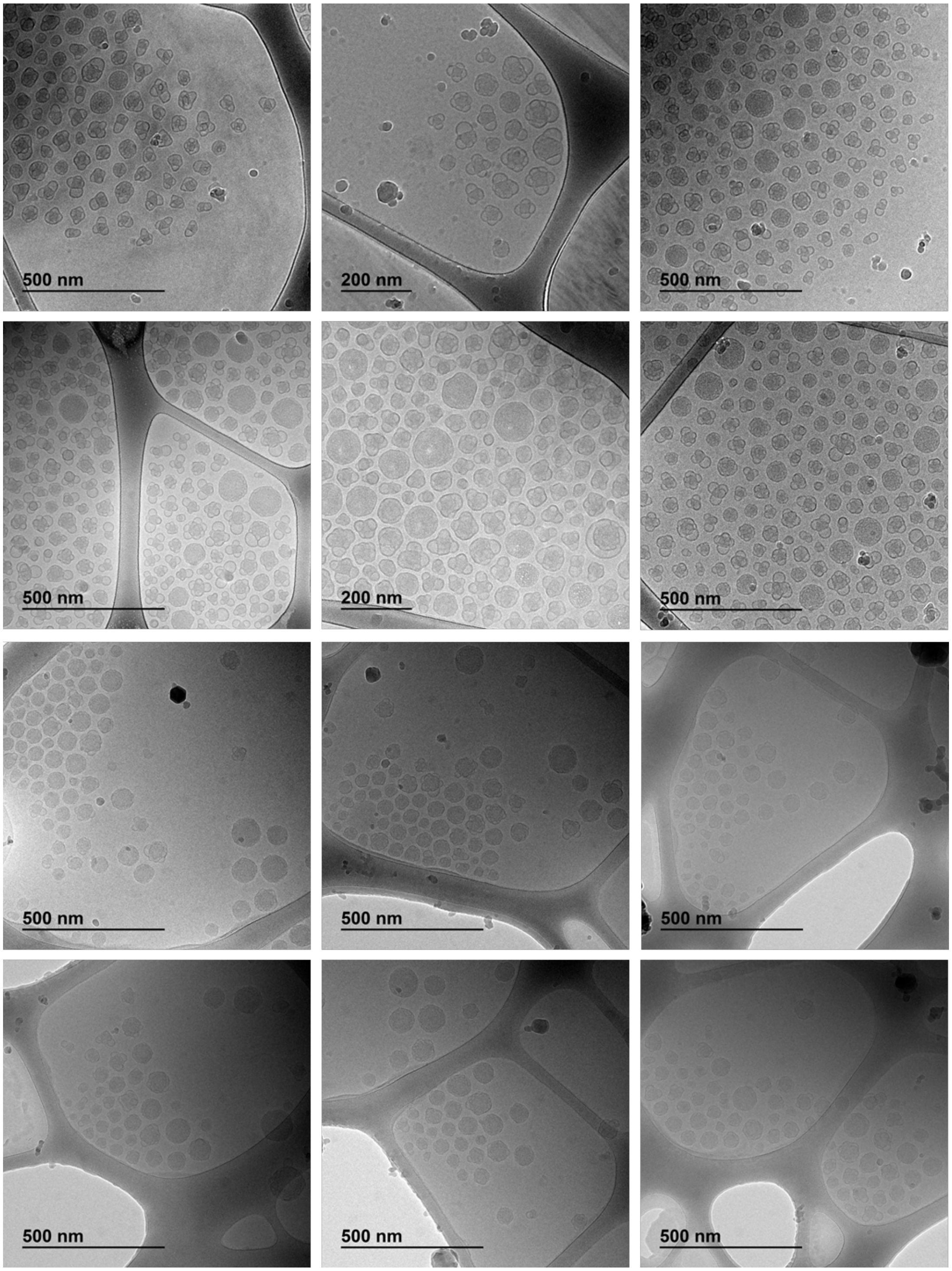


**Figure S21: Cryo-TEM Images of STING peptide-polyanion conjugate LNP formulation.** Cryo-TEM image of C3 LNP formulation containing STING conjugate (representative of N = 2 independent LNP batches, showing N = 6 fields from each batch).

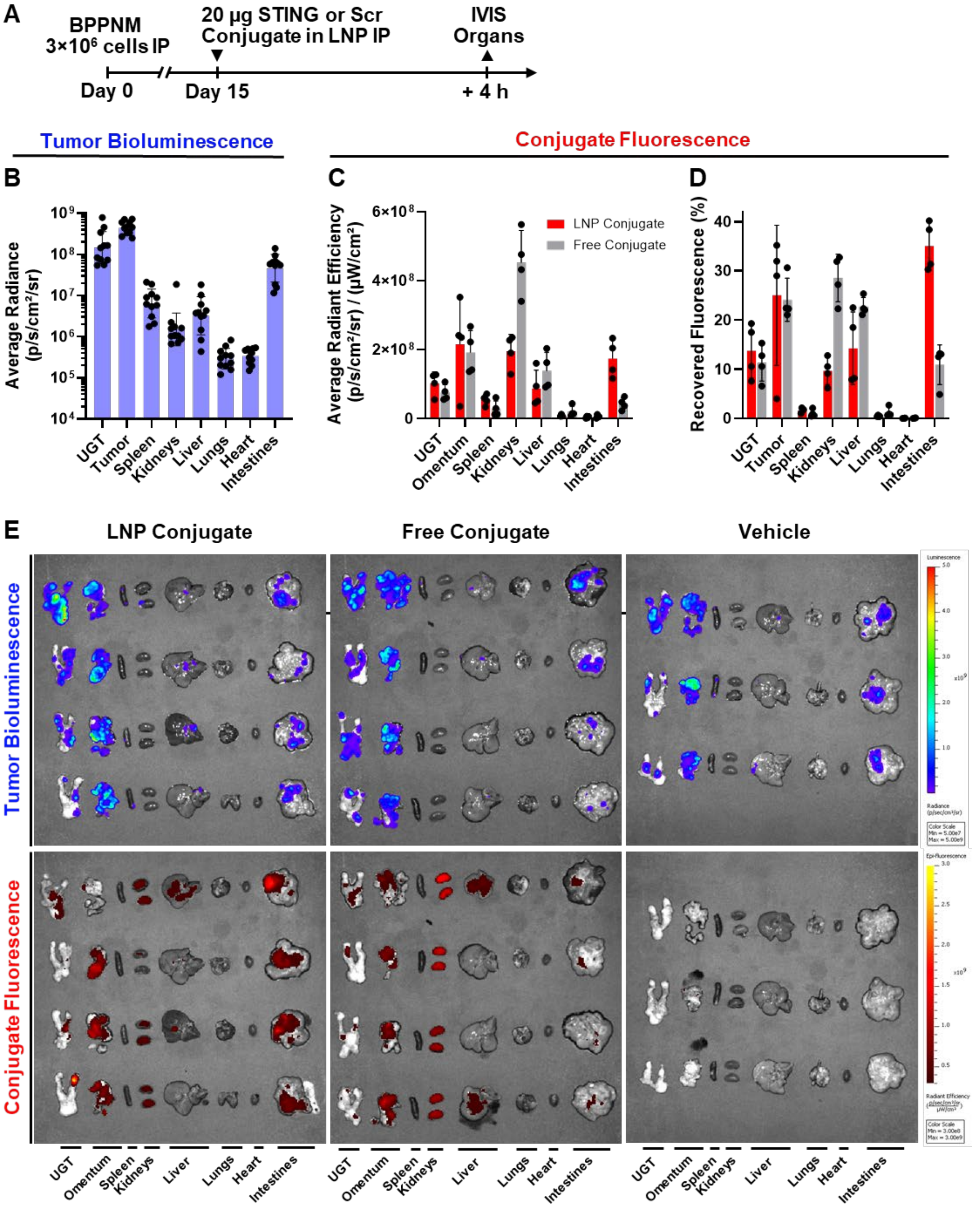


**Figure S22: Biodistribution of STING peptide-polyanion conjugate in ovarian cancer model.** (**A**) Mice were inoculated with 3×10$^6$ BPPNM cells IP and dosed with 20 µg of STING peptide conjugate delivered either by LNP or unencapsulated ("free") (N = 4 mice) or PBS vehicle (N = 3 mice) delivered by LNP IP at 15 days after inoculation. Organs were collected 4 h after dosing for *ex vivo* IVIS. (**B**) Tumor bioluminescence measured as average radiance in each organ, pooled for N = 11 mice in all groups. (**C-D**) Conjugate fluorescence measured as (**C**) average radiant efficiency and (**D**) as a fraction of total fluorescence signal recovered from all organs. (**E**) IVIS images showing tumor bioluminescence and conjugate fluorescence in each organ. Data is represented as mean ± SD.

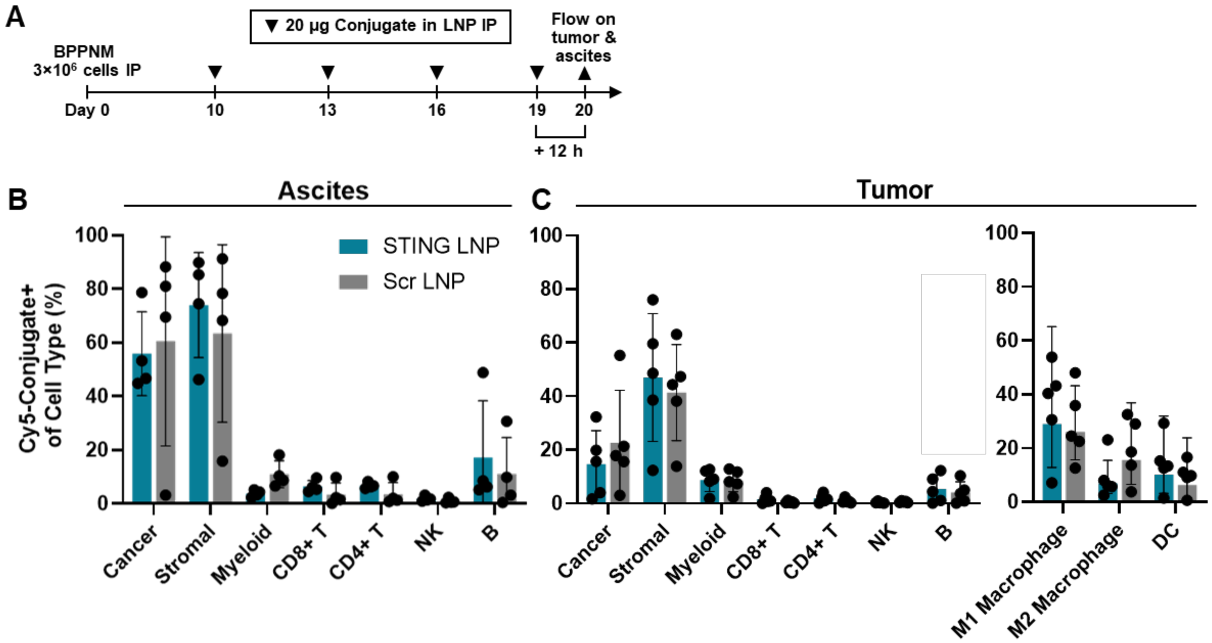


**Figure S23: Cellular biodistribution of peptide-polyanion conjugate LNP in tumor and ascites.** (**A**) Mice were inoculated with 3×10$^6$ BPPNM cells IP and dosed with 20 µg of STING or Scr peptide conjugate delivered by LNP IP at 10, 13, 16, and 19 days after inoculation. Omental tumor and ascites (peritoneal fluid) was collected on day 20, 12 h after the final dose, for analysis by flow cytometry. Groups included N = 5 (All tumors, PBS ascites) or N = 4 (STING LNP ascites, Scr LNP ascites) mice. (**B-C**) The percentage of each listed cell type that is positive for Cy5-labeled peptide-polyanion conjugate was measured in the (**B**) ascites and (**C**) tumor. Cell populations examined include BPPNM cancer cells (CD45$^-$ GFP$^+$), stromal cells (CD45$^-$ GFP$^-$), myeloid cells (CD11b$^+$), CD8$^+$ and CD4$^+$ T cells (CD3$^+$), NK cells (CD3$^-$ NK1.1$^+$), and B cells (CD19$^+$). Specific myeloid cell populations were examined in the tumor including M1 (CD86$^{hi}$ CD206$^{low}$) and M2 (CD86$^{low}$ CD206$^{hi}$) polarized macrophages (F4/80$^+$) and DCs (CD11c$^+$ MHCII$^+$). Data is represented as mean ± SD.

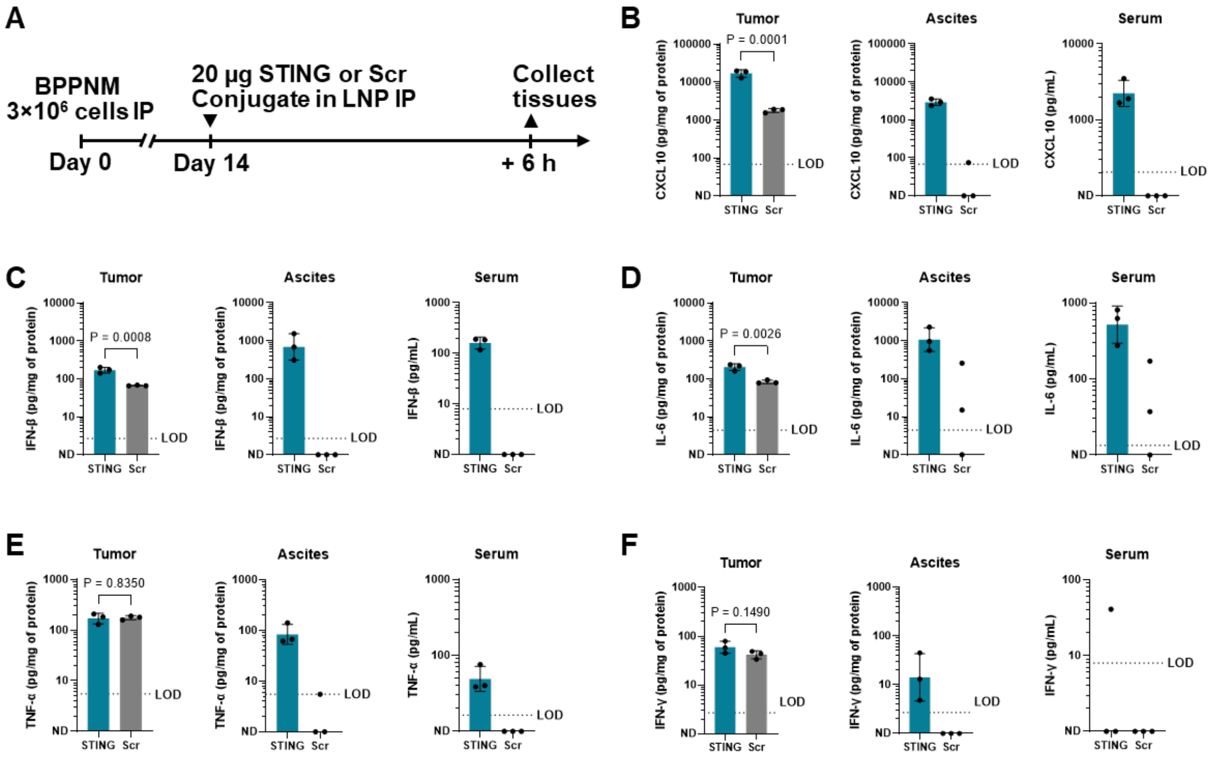


**Figure S24: Analysis of cytokines in tissues after STING peptide-polyanion conjugate treatment.** (**A**) Mice were inoculated with 3×10$^6$ BPPNM cells IP and dosed with 20 µg of STING or Scr peptide conjugate (N = 3 mice) delivered by LNP IP at 14 days after inoculation. Omental tumor, ascites, and serum were collected 6 h after dosing. Concentrations of (**B**) CXCL10, (**C**) IFN-β, (**D**) IL-6, (**E**) TNF-α, and (**F**) IFN-γ in each tissue were measured by ELISA and are reported relative to total protein concentration in tumor and ascites or relative to volume in serum. Conditions where analyte was below the LOD are labeled as ND. *P* values computed on log-transformed concentrations using an unpaired t test. Where cytokine level was ND for any replicate, no statistical tests were performed.  Data represented as geometric mean ± SD.

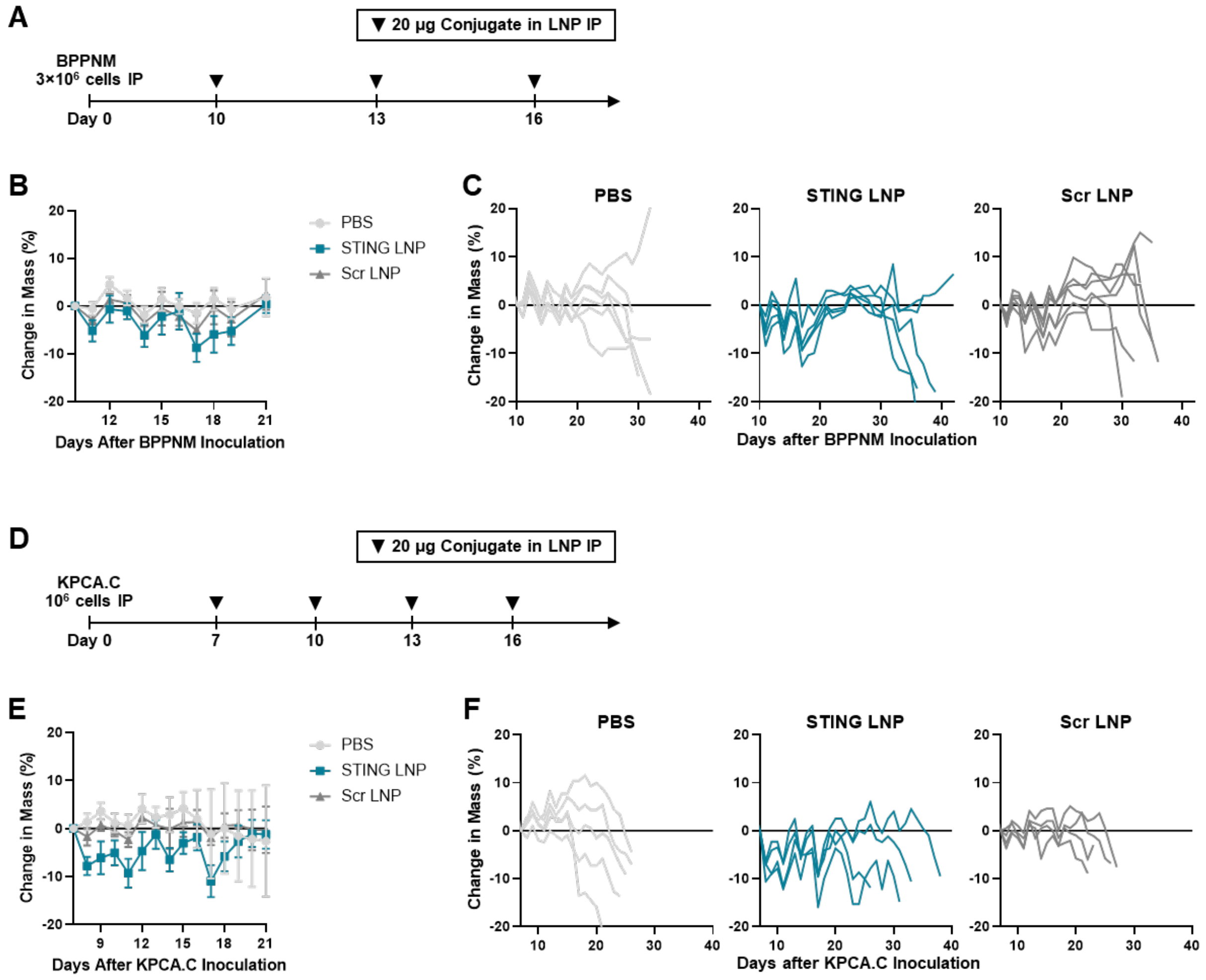


**Figure S25: Change in mouse mass during monotherapy tumor efficacy studies.** (**A**) Mice were inoculated with $3\times10^6$ BPPNM cells IP and dosed with 20 µg of STING or Scr peptide-polyanion conjugate delivered by LNP IP at 10, 13, and 16 days after inoculation. Groups included N = 6 (PBS, Scr LNP) or N = 5 (STING LNP) mice. (**B-C**) Change in mouse mass was measured, displaying (**B**) mean ± SD over the treatment period and (**C**) individual mouse changes in mass. (**D**) Mice were inoculated with $10^6$ KPCA.C cells IP and dosed with 20 µg of STING or Scr peptide-polyanion conjugate delivered by LNP IP at 7, 10, 13, and 16 days after inoculation. Groups included N = 5 (PBS), or N = 4 (STING LNP, Scr LNP) mice. (**E-F**) Change in mouse mass was measured, displaying (**E**) mean ± SD over the treatment period and (**F**) individual mouse change in mass.

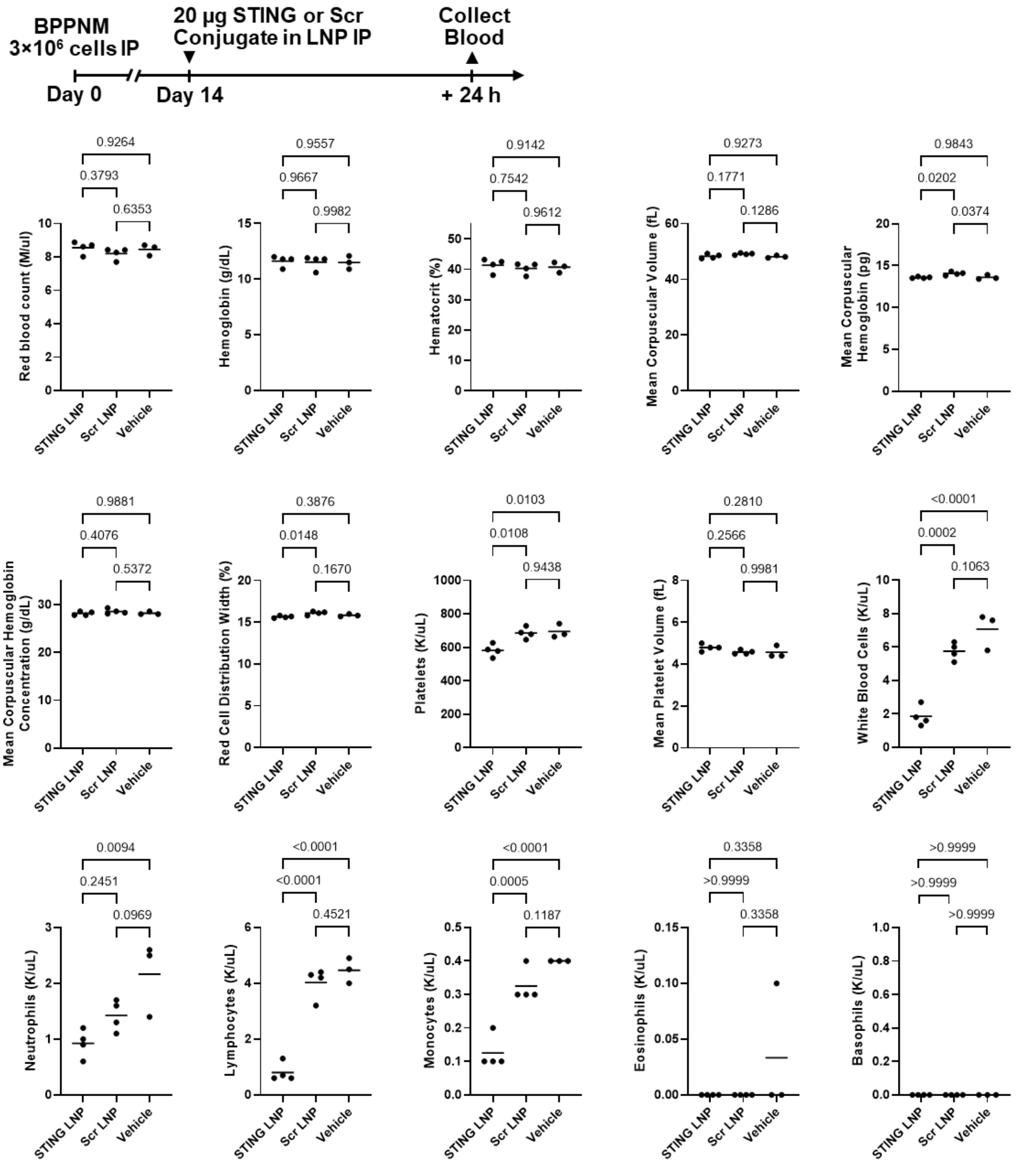


**Figure S26: Acute toxicity study hematology measurements.** Mice were inoculated with 3×10$^6$ BPPNM cells IP and dosed with 20 µg of STING or Scr peptide-polyanion conjugate delivered by LNP IP at 14 days after inoculation. Groups included N = 4 (STING LNP, Scr LNP) or N = 3 (PBS) mice. 24 h after treatment, blood was collected for complete blood counts. *P* values computed with a one-way ANOVA followed by Tukey's post-hoc test are displayed above each figure. Data represented as mean ± SD.

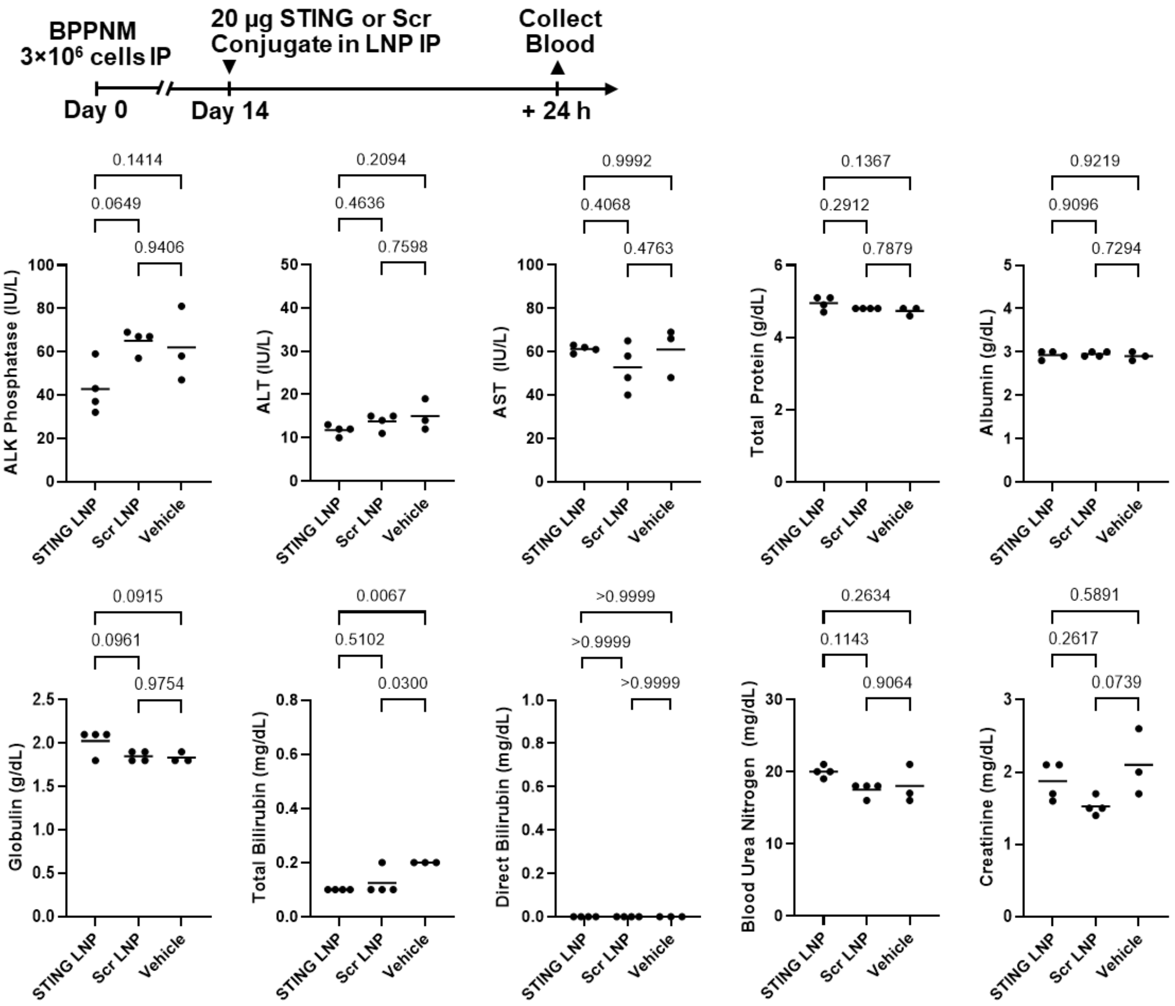


**Figure S27: Acute toxicity study serum chemistry measurements.** Mice were inoculated with 3×10$^6$ BPPNM cells IP and dosed with 20 µg of STING or Scr peptide-polyanion conjugate delivered by LNP IP at 14 days after inoculation. Groups included N = 4 (STING LNP, Scr LNP) or N = 3 (PBS) mice. 24 h after treatment, blood was collected for analysis of serum chemistry. *P* values computed with a one-way ANOVA followed by Tukey's post-hoc test are displayed above each figure. Data represented as mean ± SD.

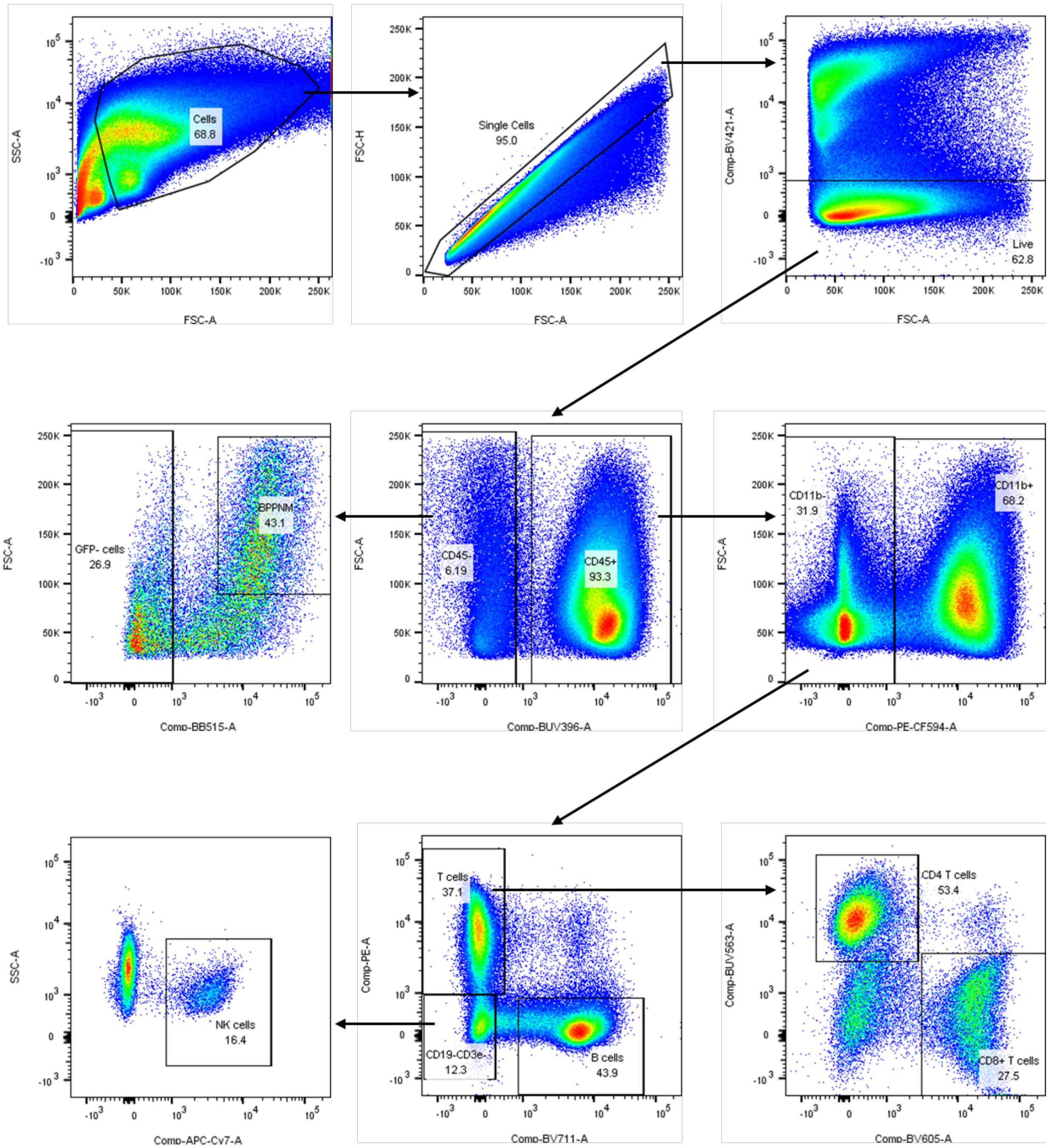


**Figure S28: Gating strategy used for flow cytometry lymphoid panel.**

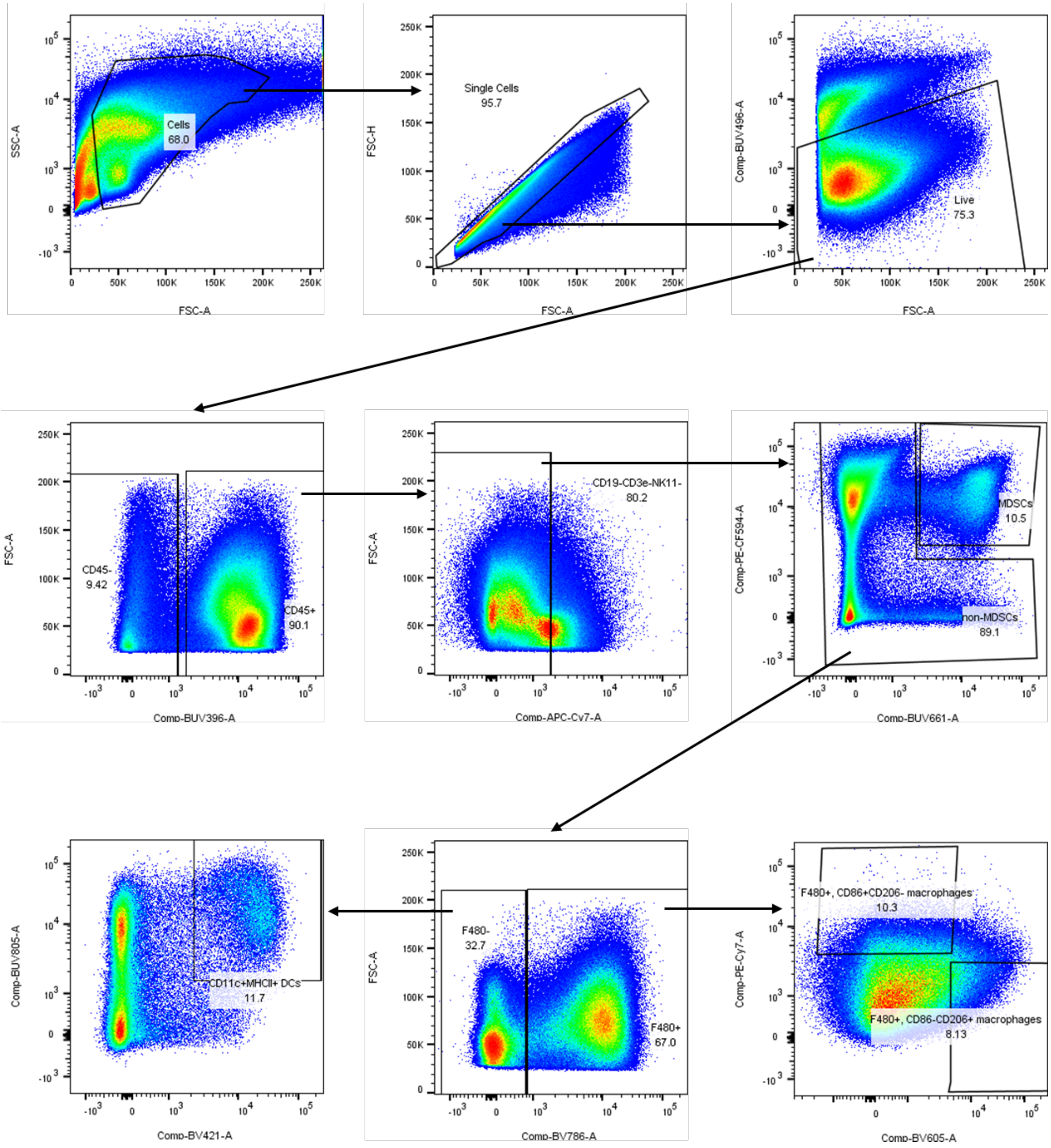


**Figure S29: Gating strategy used for flow cytometry myeloid panel.**

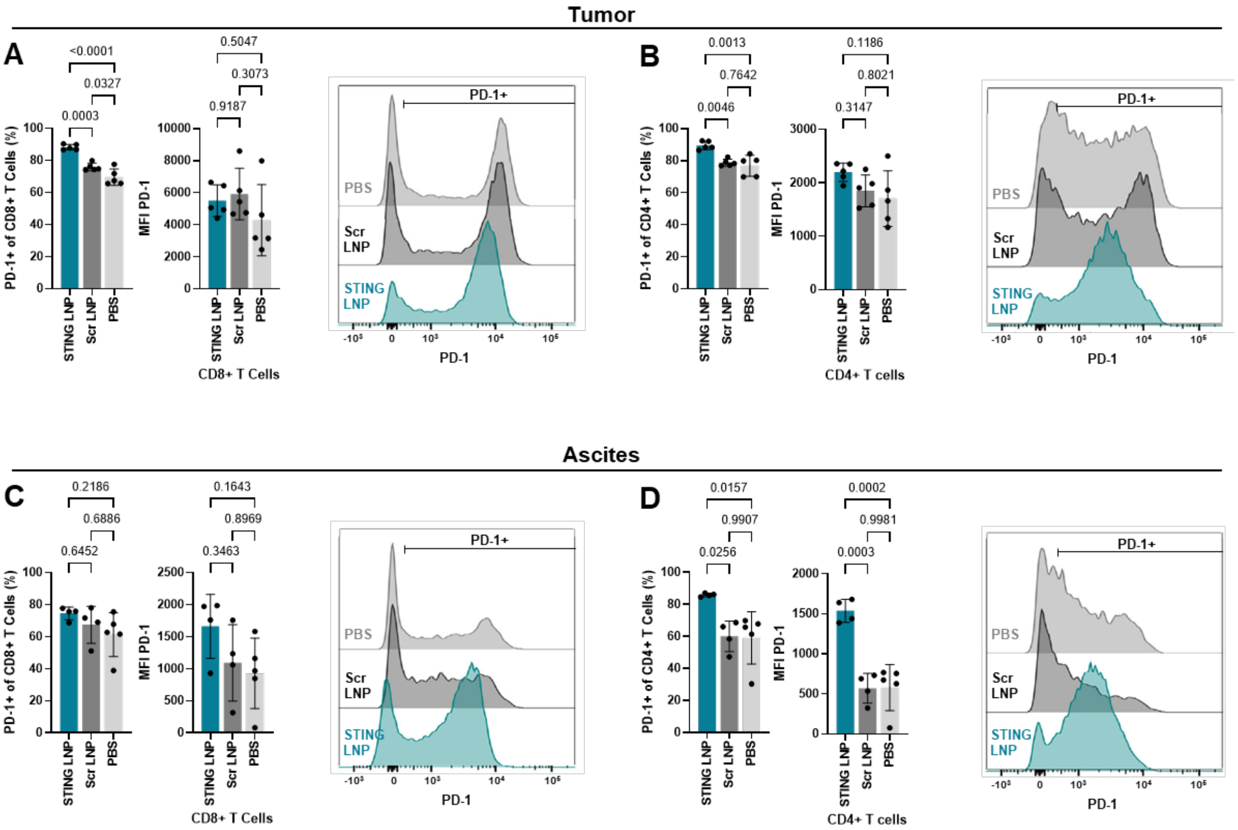


**Figure S30: STING peptide-polyanion conjugate treatment impact on T cell PD-1 expression.** Mice were inoculated with $3\times10^6$ BPPNM cells IP and dosed with 20 µg of STING or Scr peptide-polyanion conjugate delivered by LNP IP at 10, 13, 16, and 19 days after inoculation. Omental tumor and ascites (peritoneal fluid) was collected on day 20 for analysis by flow cytometry. Groups included N = 5 (All tumors, PBS ascites) or N = 4 (STING LNP ascites, Scr LNP ascites) mice. Expression of PD-1 on (**A**) $CD8^+$ and (**B**) $CD4^+$ T cells in the tumor as well as (**C**) $CD8^+$ and (**D**) $CD4^+$ T cells in the ascites, showing percentage of cells that are PD-$1^+$, PD-1 MFI, and representative distributions of PD-1. *P* values computed with a one-way ANOVA followed by Tukey's post-hoc test are displayed above each figure. Data represented as mean ± SD.

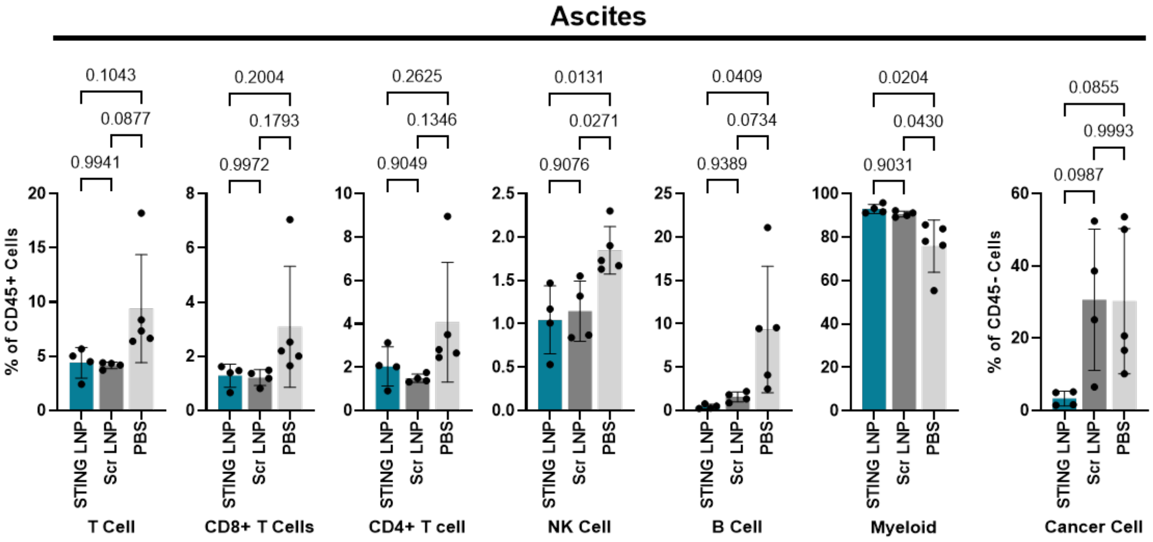


**Figure S31: STING peptide-polyanion conjugate treatment impact on ascites cell populations.** Mice were inoculated with $3\times10^6$ BPPNM cells IP and dosed with 20 µg of STING or Scr peptide-polyanion conjugate delivered by LNP IP at 10, 13, 16, and 19 days after inoculation. Ascites (peritoneal fluid) was collected on day 20 for analysis by flow cytometry. Groups included N = 5 (PBS) or N = 4 (STING LNP, Scr LNP) mice. Cell populations in tumor after treatment, displaying percentage of $CD45^+$ cells made up by T cells ($CD3^+$), $CD8^+$ T cells, $CD4^+$ T cells, NK cells ($CD3^-$ $NK1.1^+$), B cells ($CD19^+$), or Myeloid cells ($CD11b^+$). Percentage of CD45- cells made up by BPPNM cancer cells ($GFP^+$) are also displayed. *P* values computed with a one-way ANOVA followed by Tukey's post-hoc test are displayed above each figure. Data represented as mean ± SD.

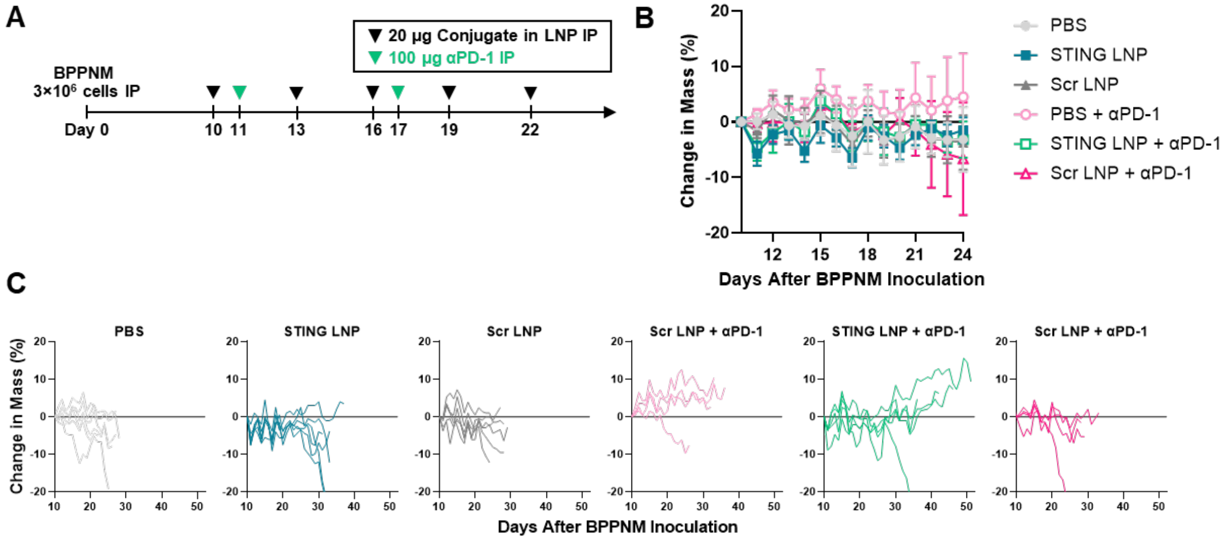


**Figure S32: Change in mouse mass during combination therapy tumor efficacy studies.** (**A**) Mice were inoculated with 3×10$^6$ BPPNM cells IP and dosed with 20 µg of STING or Scr peptide-polyanion conjugate delivered by LNP IP at 10, 13, 16, 19, and 22 days after inoculation. A subset of groups were additionally treated with 100 µg αPD-1 antibody at 11 and 17 days after inoculation. Groups included N = 6 (PBS, STING LNP, Scr LNP), N = 5 (STING LNP + αPD-1 ), or N = 4 (PBS + αPD-1, Scr LNP + αPD-1) mice. (**B-C**) Change in mouse mass was measured daily, displaying (**B**) mean ± SD over the treatment period and (**C**) individual mouse change in mass.

**Table S1: Compositions of all LNP formulations examined**

| Formulation | Total lipid to cargo (mass ratio) | Ionizable lipid (mol %) | Phospholipid (mol %) | Cholesterol (mol %) | PEG lipid (mol %) |
|---|---|---|---|---|---|
| **ALC-0315** | 16.8* | ALC-0315 (50.0) | DOPE (10.0) | Cholesterol (38.5) | DMG-PEG2000 (1.5) |
| **MC3** | 18.2* | DLin-MC3-DMA (50.0) | DOPE (10.0) | Cholesterol (38.5) | DMG-PEG2000 (1.5) |
| **SM-102** | 17.3* | SM-102 (50.0) | DOPE (10.0) | Cholesterol (38.5) | DMG-PEG2000 (1.5) |
| **A1** | 20.0 | ALC-0315 (35.0) | DOPE (25.0) | Cholesterol (38.5) | DMG-PEG2000 (1.5) |
| **A2** | 20.0 | ALC-0315 (50.0) | DOPE (10.0) | Cholesterol (38.5) | DMG-PEG2000 (1.5) |
| **A3** | 20.0 | ALC-0315 (43.8) | DOPE (31.3) | Cholesterol (23.5) | DMG-PEG2000 (1.5) |
| **A4** | 20.0 | ALC-0315 (62.5) | DOPE (12.5) | Cholesterol (23.5) | DMG-PEG2000 (1.5) |
| **A5** | 20.0 | ALC-0315 (52.8) | DOPE (16.5) | Cholesterol (29.2) | DMG-PEG2000 (1.5) |
| **B1** | 20.0 | ALC-0315 (35.0) | DSPC (25.0) | Cholesterol (38.5) | DMG-PEG2000 (1.5) |
| **B2** | 20.0 | ALC-0315 (50.0) | DSPC (10.0) | Cholesterol (38.5) | DMG-PEG2000 (1.5) |
| **B3** | 20.0 | ALC-0315 (43.8) | DSPC (31.3) | Cholesterol (23.5) | DMG-PEG2000 (1.5) |
| **B4** | 20.0 | ALC-0315 (62.5) | DSPC (12.5) | Cholesterol (23.5) | DMG-PEG2000 (1.5) |
| **B5** | 20.0 | ALC-0315 (52.8) | DSPC (16.5) | Cholesterol (29.2) | DMG-PEG2000 (1.5) |
| **C1** | 20.0 | ALC-0315 (35.0) | DOPC (25.0) | Cholesterol (38.5) | DMG-PEG2000 (1.5) |
| **C2** | 20.0 | ALC-0315 (50.0) | DOPC (10.0) | Cholesterol (38.5) | DMG-PEG2000 (1.5) |
| **C3** | 20.0 | ALC-0315 (43.8) | DOPC (31.3) | Cholesterol (23.5) | DMG-PEG2000 (1.5) |
| **C4** | 20.0 | ALC-0315 (62.5) | DOPC (12.5) | Cholesterol (23.5) | DMG-PEG2000 (1.5) |
| **C5** | 20.0 | ALC-0315 (52.8) | DOPC (16.5) | Cholesterol (29.2) | DMG-PEG2000 (1.5) |

*Initial comparison between ionizable lipids was carried out at a fixed 10:1 mass ratio of ionizable lipid to cargo.

**Table S2: Antibody panels used for flow cytometry experiments**

| Marker | Fluorophore | Clone | Product Info | Dilution |
|---|---|---|---|---|
| ***Lymphoid Panel*** | | | | |
| CD45 | BUV 395 | 30-F11 | BD Biosciences #564279 | 40 |
| CD11b | PE-Dazzle 594 | M1/70 | BioLegend #101256 | 400 |
| NK1.1 | APC-Cy7 | PK136 | BioLegend #108724 | 40 |
| CD3e | PE | 17A2 | BioLegend #100206 | 20 |
| CD19 | BV711 | 6D5 | BioLegend #115555 | 80 |
| CD4 | BUV563 | GK1.5 | BD Biosciences #612923 | 40 |
| CD8 | BV605 | 53–6.7 | BioLegend #100744 | 40 |
| PD-1 | PE-Cy7 | clone 29F.1A12 | BioLegend #135216 | 60 |
| Zombie Violet | | | BioLegend #423113 | 100 |
| ***Myeloid Panel*** | | | | |
| CD45 | BUV 395 | 30-F11 | BD Biosciences #564279 | 40 |
| MHC-II (I-A/I-E) | BUV805 | M5/114.15.2 | BD Biosciences #748844 | 60 |
| CD11c | BV421 | N418 | BioLegend #117330 | 20 |
| F4/80 | BV785 | BM8 | BioLegend #123141 | 20 |
| CD11b | PE-Dazzle 594 | M1/70 | BioLegend #101256 | 400 |
| Ly6C/Ly6G (Gr-1) | BUV 661 | RB6-8C5 | BD BioSciences #741470 | 400 |
| CD206 | BV605 | C068C2 | BioLegend #141721 | 20 |
| CD86 | PE-Cy7 | GL-1 | BioLegend #105014 | 20 |
| CD19 | APC-Cy7 | 6D5 | BioLegend #115530 | 40 |
| CD3e | APC-Cy7 | 17A2 | BioLegend #100222 | 40 |
| NK1.1 | APC-Cy7 | PK136 | BioLegend #108724 | 40 |
| Zombie UV | | | BioLegend #423107 | 100 |
